# Poseidon – A framework for archaeogenetic human genotype data management

**DOI:** 10.1101/2024.04.12.589180

**Authors:** Clemens Schmid, Ayshin Ghalichi, Thiseas C. Lamnidis, Dhananjaya B. A. Mudiyanselage, Wolfgang Haak, Stephan Schiffels

## Abstract

The study of ancient human genomes, archaeo- or palaeogenetics, has accelerated in the last ten years, with now thousands of new ancient genomes being released each year. Operating at the interface of genetics, anthropology and archaeology, this data includes features from all three fields, including rich meta- and context-data, for example regarding spatiotemporal provenience. While archives and standards for genetic sequencing data already exist, no such infrastructure exists for combined genetic and meta-data that could ensure FAIR principles across the field. Here, we present Poseidon, a framework for open and FAIR data handling in archaeogenetics, including a specified package format, software tools, and public, community-maintained online archives. Poseidon emphasises human- and machine-readable data storage, the development of convenient and interoperable command line software, and a high degree of source granularity to elevate the original data publication to the main unit of long-term curation.

## 2 Introduction

The technology to extract and sequence DNA from human remains thousands of years old has revolutionised the study of the human past. This is documented by groundbreaking new insights, from our evolutionary relationships to distant relatives like Neanderthals [1, 2] to prehistoric [3, 4] and historic migrations [5, 6]. Since the sequencing of the first ancient modern human genome in 2010 [7], hundreds of studies have been published, accompanied by massive datasets of ancient human DNA sequences. A drop in sequencing costs and new technologies like hybridisation capture [8, 4, 9] have in fact lead to an acceleration of new published ancient genomes, with data now coming out faster than individual researchers typically can keep track of and co-analyze. Recently, the threshold of genome-wide data for 10,000 ancient human individuals has been surpassed [10].

To make all this new data publicly available, researchers can partly rely on existing infrastructure for the archival and distribution of modern genetic data, such as the Sequence Read Archive (SRA) [11], the European Nucleotide Archive (ENA) [12] or other INSDC databases (https://www.insdc.org). However, this infrastructure has not been prepared to also capture the rich context-data ranging from archaeological field observations to radiocarbon dating that accompanies ancient samples. Nor is there a standardised archive yet for derived genotype data that is routinely used to substantiate most if not all of the conclusions in archaeogenetic papers. This raises multiple concrete issues:

- Ancient individuals only constitute meaningful observations if their spatiotemporal provenience is known. Current practice renders it difficult to maintain the connection between archaeological context and sampled genomic data, as this information is generally kept separately.
- Specific results of typical archaeogenetic analyses (e.g. PCA, F-Statistics, kinship estimation) can only be fully reproduced with genotype-level data. But current practice is not to include this data with a publication, be it for its unwieldy size or the lack of a central repository to easily share it.
- Meta-analyses involving large amounts of data require tedious curation. Despite the fact that archaeogenetic genotype data is largely standardized, and common practices for reporting processing steps, data quality indicators, as well as spatial and temporal sample origin have emerged, combining information from different papers is still hindered by severe structural variation.

A major project addressing some of these problems in human archaeogenetics is the Allen Ancient DNA Resource (AADR), which is a curated dataset of public ancient DNA data assembled by the Reich Lab at Harvard University [13]. While the AADR clearly fills a gap in the field, there continues to be a need for open standardization and the creation of a community-maintained archive. Both standard and archive should be well-specified to be human and machine-readable, fully open and transparent, and permanently downloadable with their entire version history. They should be geared towards scientific practice and include well-documented interfaces for research software. To ensure fairness, i.e. equal access for users and contributors from different backgrounds, and long-term reliability, they should be as independent as possible from specific institutions, key persons [14] or infrastructure providers, and ideally controlled by the community-of-practice as a whole.

Here, we present ‘*Poseidon*’ (https://www.poseidon-adna.org), a computational framework including an open data format, software, and community-maintained archives, to enable this standardised and FAIR handling of archaeogenetic data. The name *Poseidon* is inspired by the notion of a *sea of data* benefiting from structure and governance. We imagine Poseidon to simplify a variety of concrete workflows and mitigate technical challenges practitioners in the field of human archaeogenetics regularly face:

### Data storage

Archaeogenetic samples/ancient individuals (we often use these terms synonymously below, see Supplementary Text 8 for a definition) can only be effectively analysed with context data. The Poseidon package format allows one to store archaeogenetic genotype data with arbitrary spatiotemporal or archaeological information on a per-sample level. The package format enforces human- and machine-readability for the context data to enable its computational co-analysis with the genomic data, while also maintaining a high level of flexibility and extensibility to accommodate arbitrary additional variables.

### Data acquisition

Research in archaeogenetics is strongly dependent on incorporating published reference data. Poseidon features public archives with per-article packages that can be downloaded through an open web API. The packages include genotypes, context data and machine-readable citation information. To ensure computational reproducibility, Poseidon supports version tracking for these packages, with old versions directly available through the web interface. Beyond hosting data, the web infrastructure also provides options to report issues and suggest transparent data updates.

### Data analysis

Working with common software tools in human archaeogenetics (e.g. smartpca [16, 17], ADMIXTURE [18], qpAdm [4, 19]) requires frequent merging, sub-setting and file format conversion for the samples of interest. The Poseidon core team develops the software tool *trident* to simplify these operations both with Poseidon packages and unpackaged genotype data. The unified package format allows for new analysis tools with flexible data detection, low-memory stream processing and on-the-fly aggregation of widely used genome-wide statistics as demonstrated in the *xerxes* software tool also developed by the core team.

### Data publication

For each sample, human archaeogenetic papers should include genotypes, context information and quality-control data, so e.g. estimates for the aDNA damage or contamination. Poseidon offers a standardized and reusable way to share this data directly with the publication and/or through our public archives.

## 3 Overview

Poseidon consists of three major components: A data format, software, and public archives (see Figure 1). At the foundation stands a data format to store genotype data together with context information. The software implements this specification, relies on and validates its promises, and builds convenient, useful functionality on top of it, both for individual users and for our cloud infrastructure. Finally, the public archives store data using the data format and employ the software for validation, modification and book-keeping.

**Figure 1:**
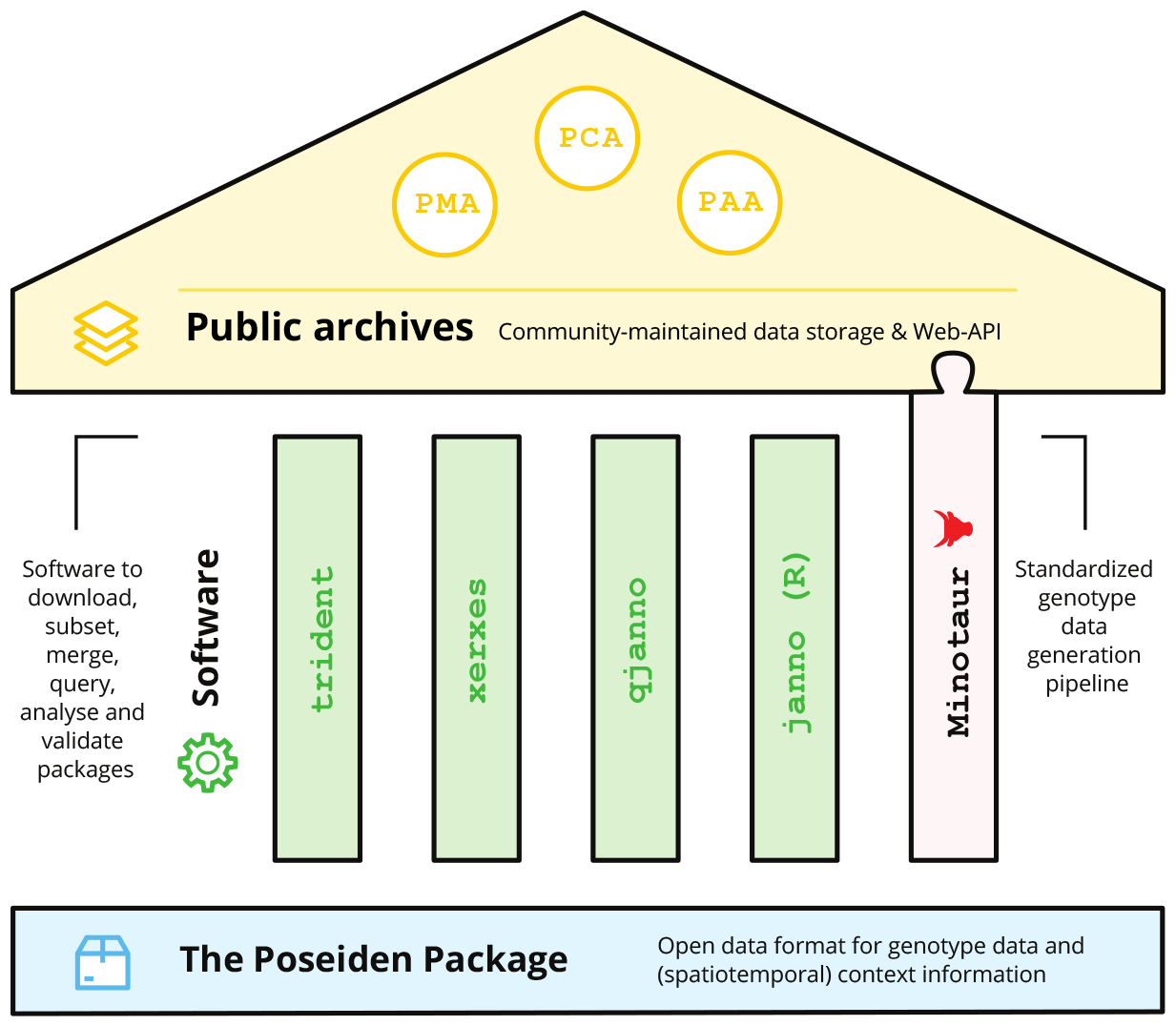
Schematic overview of the core components of the Poseidon framework. The Poseidon package specification forms the foundation on which various open source software tools and the Minotaur workflow are based. They, in turn, underpin and enable the public Poseidon data archives.

The Poseidon schema (Supplementary Text 1) defines the structure and format of a Poseidon package (Figure 2). This includes the general layout and purpose of the main files, the POSEIDON.yml, the .janno and the .ssf file, and detailed definitions of the variables within them, both regarding semantics and syntax (Supplementary Text 2). The short definitions in the schema are explained in more detail on the Poseidon website, which also serves as a central hub for all components of the framework.

**Figure 2:**
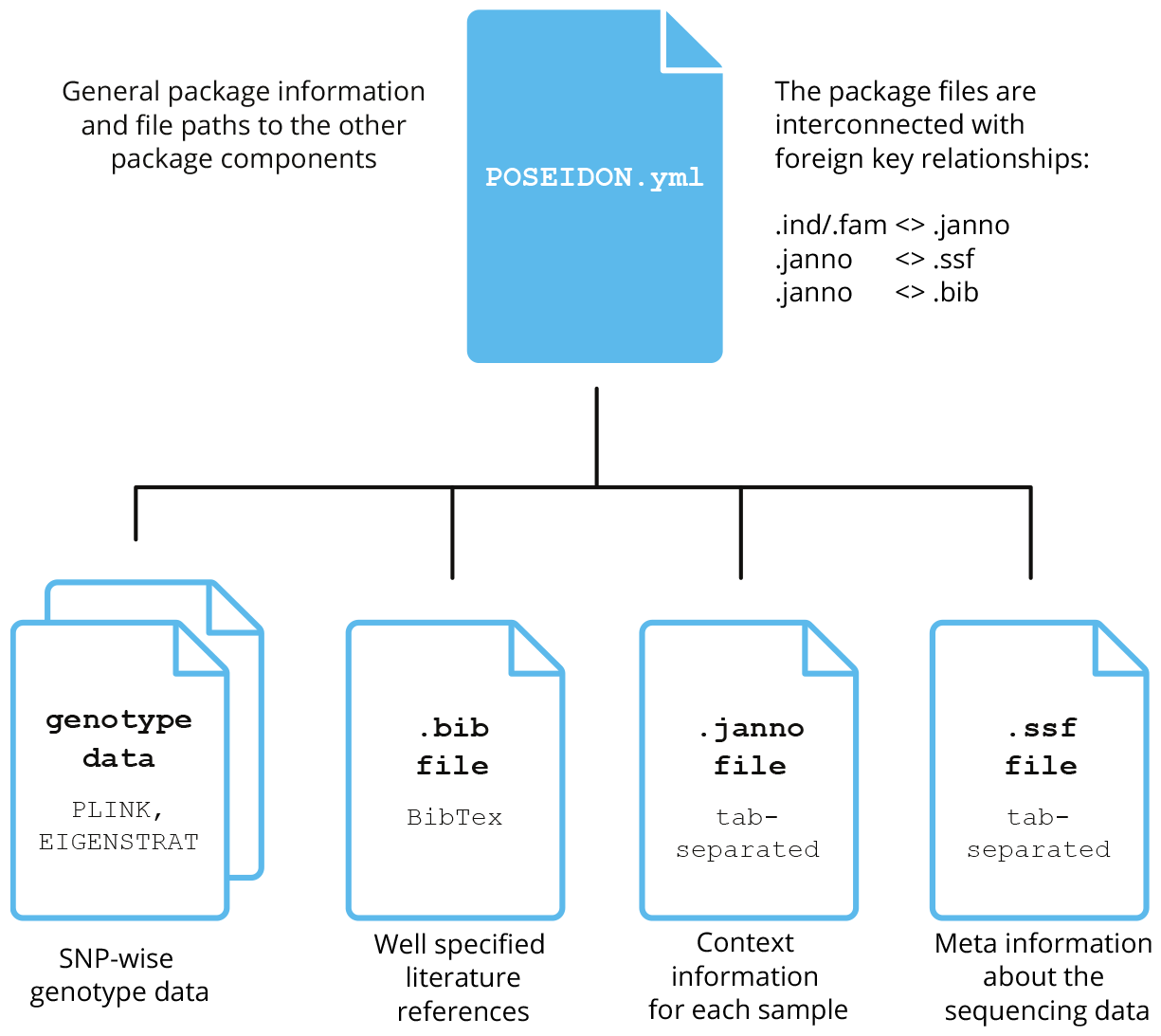
Schematic overview of the Poseidon package structure. The POSEIDON.yml file defines the package and interlinks the additional data files for genotype, context- and bibliography information in a relational structure.

The Poseidon software suite enables users to create, download, inspect, subset, merge and analyse Poseidon packages (Supplementary Text 3-7). It is mostly implemented in Haskell [20], a purely functional programming language, and split over multiple command line tools openly available as statically compiled executables for the major desktop and server operating systems. The main tool, *trident*, provides data handling functionality, including the code necessary for the client-server infrastructure of the public archives. The *xerxes* software implements commonly used genome-wide allele-frequency statistics, *qjanno* allows for SQL-like queries on .janno files and the *janno* R package offers a .janno file interface for the R programming language.

Using these tools, we conceived and implemented three public data archives for sharing and maintaining published (ancient) human DNA data packaged in the Poseidon format: The Poseidon Community Archive (PCA), the Poseidon Minotaur Archive (PMA) and the Poseidon AADR Archive (PAA). They already store considerable amounts of public genotype data (Figure 4, Figure 5 and Supplementary Text 8). Each of these archives is at their base represented by a public repository on GitHub, versioned with Git and capable of holding large genotype data files through an integration with Git’s Large File Storage (LFS) functionality. This setup allows for community driven maintenance through Git pull requests (Figure 3). A custom webserver mirrors the data from GitHub with an explicit version history, and allows to query the archive data and download packages in the browser, from the command line and through web-frontends. While the PCA focuses on author-submitted genotype data, the PMA only accepts data that went through a specific standardized genotype data processing pipeline, the Minotaur workflow, with a semi-automatic interface on GitHub and a processing queue currently hosted on computational infrastructure at the Max Planck Institute for Evolutionary Anthropology (MPI-EVA). Finally, the PAA stores releases of the AADR dataset in Poseidon format, after a light-weight curation step to ensure format compatibility.

**Figure 3:**
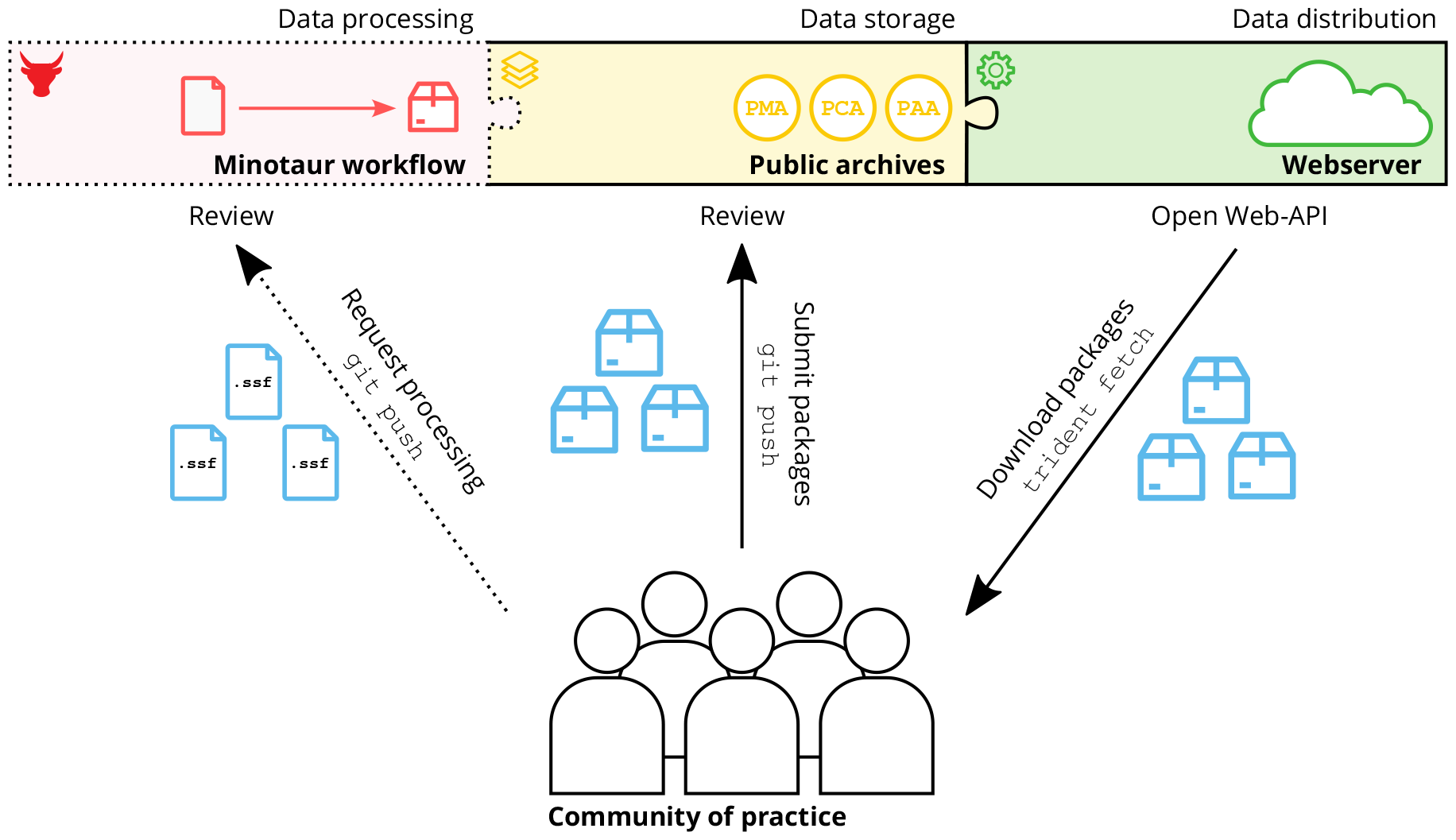
Schematic overview of the most common interaction pathways between the archaeogenetic community of practice and the Poseidon infrastructure. Community members share data either by preparing .ssf files as build-instructions for the Minotaur workflow or by directly submitting packages to the Community Archive. The data in the archives can then in turn be downloaded from the archives through the Poseidon webserver and its API.

**Figure 4:**
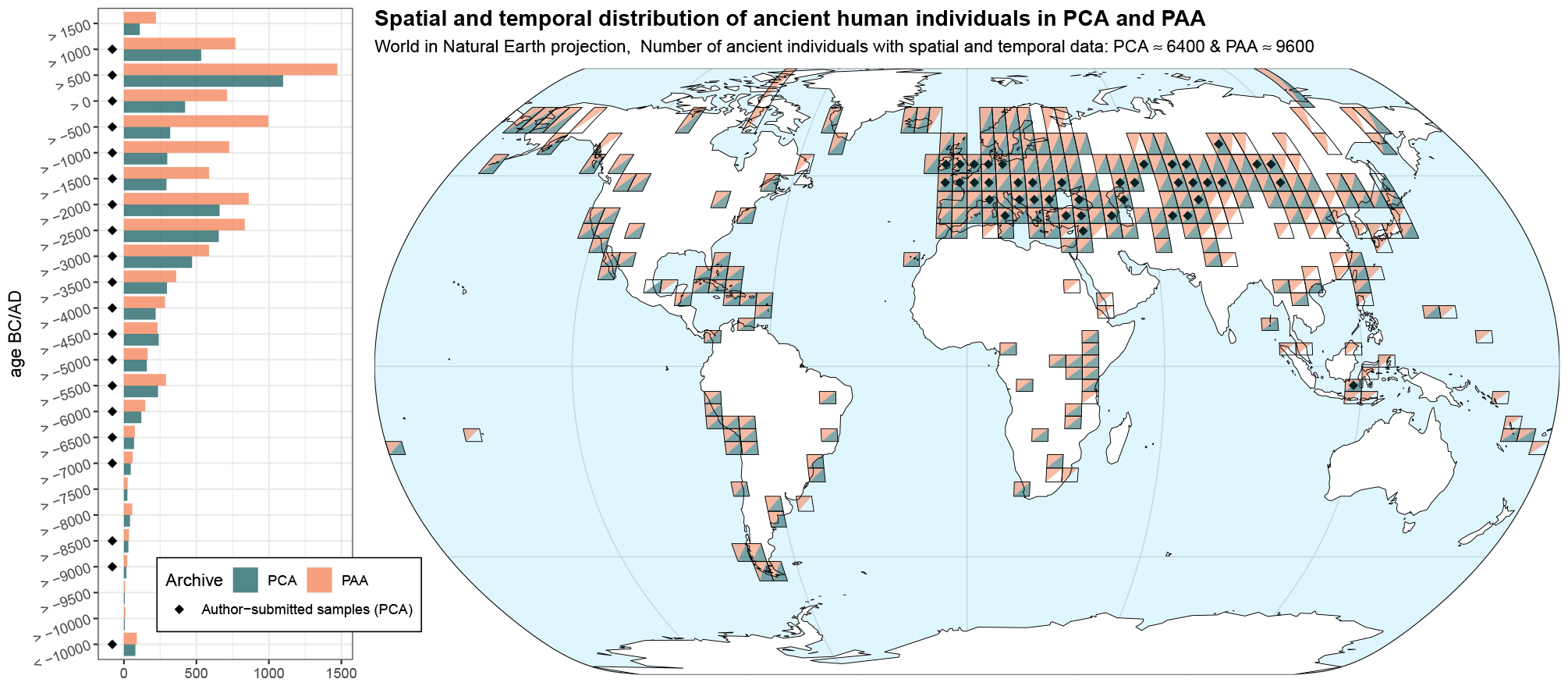
Spatiotemporal distribution of ancient individuals in the PCA and PAA (AADR v54.1.p1) public Poseidon archives. See Supplementary Text 8 for an explanation of how individuals were counted. The map shows the qualitative presence and absence of samples from both archives in a 5^*°*^-resolution grid. Especially highlighted are areas and time periods for which the PCA includes samples from currently 13 author-submitted Poseidon packages.

**Figure 5:**
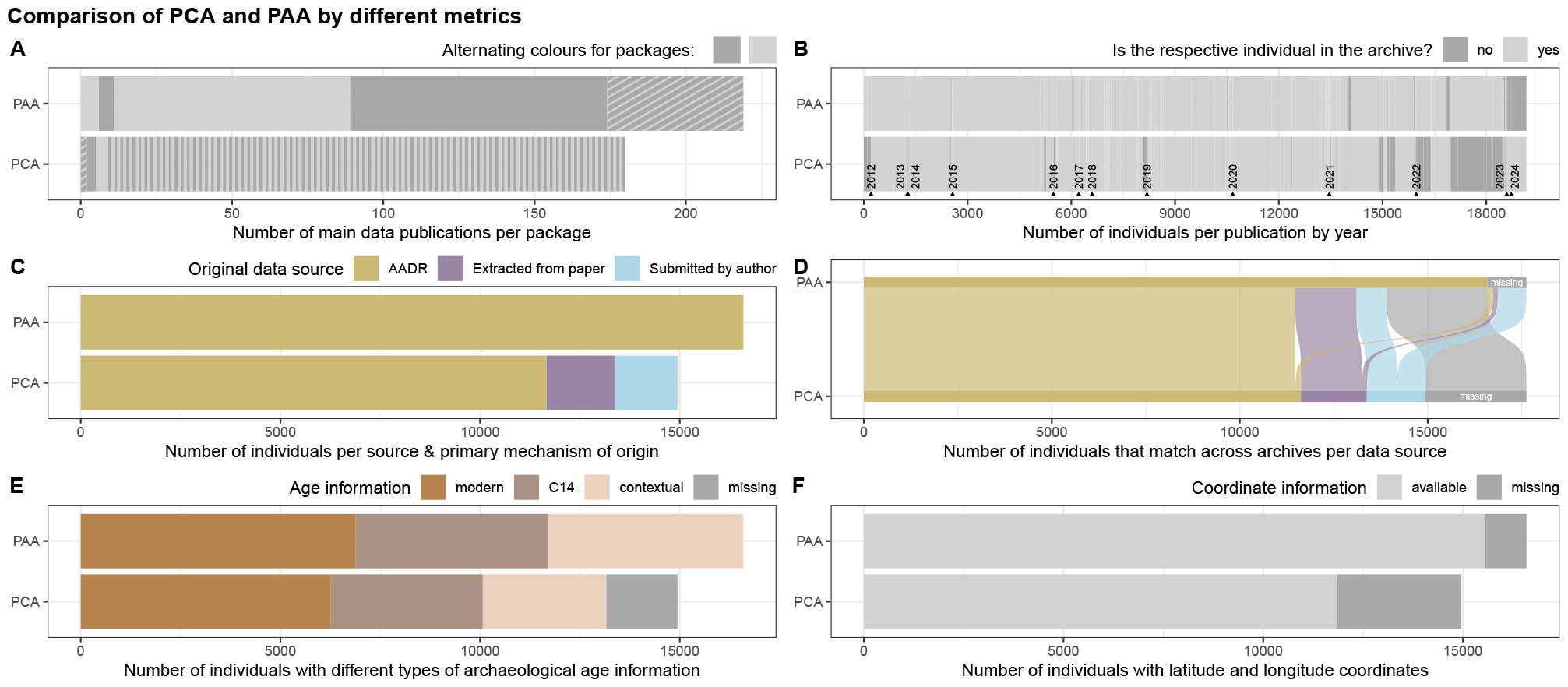
Multiple charts to compare the current content of the PCA and PAA (AADR v54.1.p1) public Poseidon archives. See Supplementary Text 8 for an explanation of how individuals were counted and more in-depth descriptions of each chart. A) Stacked barchart of publications and how they are distributed across packages in PCA and PAA. Publications represented in multiple packages are counted towards the shaded area to get a correct total, B) Barcode plot of individuals available in each archive per publication through time, C) Stacked barchart of individuals and how they were added to the archives, D) Sankey diagram of individuals matching across PCA and PAA, highlighting individuals unique to each archive on the right, E) Stacked barchart of dating information per individual available in the archives, F) Stacked barchart of spatial coordinate coverage in the archives.

## 4 Structure

The following sections explain the different components of Poseidon in detail: The Poseidon package, the software tools, and the public archives including the Minotaur processing workflow. The underlying repositories with specifications and code are available on GitHub (https://github.com/poseidon-framework) and, each in a current version, at a long-term archive: https://doi.org/10.17605/OSF.IO/ZUQGB

### 4.1 The Poseidon Package specification (v2.7.1)

The core idea of Poseidon is to organize genotype data together with relevant meta- and context data in a structured yet flexible, human- and machine-readable format. This format is the Poseidon package, defined in a semantically versioned [21] specification, openly available online (https://github.com/poseidon-framework/poseidon-schema), and also part of this publication as Supplementary Text 1. A Poseidon package must contain a POSEIDON.yml file and genotype data in PLINK or EIGENSTRAT format. It should additionally contain a .janno file to store sample-wise context information and a .bib file for literature references. Optionally, it can also contain an .ssf file with information on the underlying raw sequencing data, an unstructured README.md file for arbitrary meta information and a CHANGELOG.md file to document changes to the package.

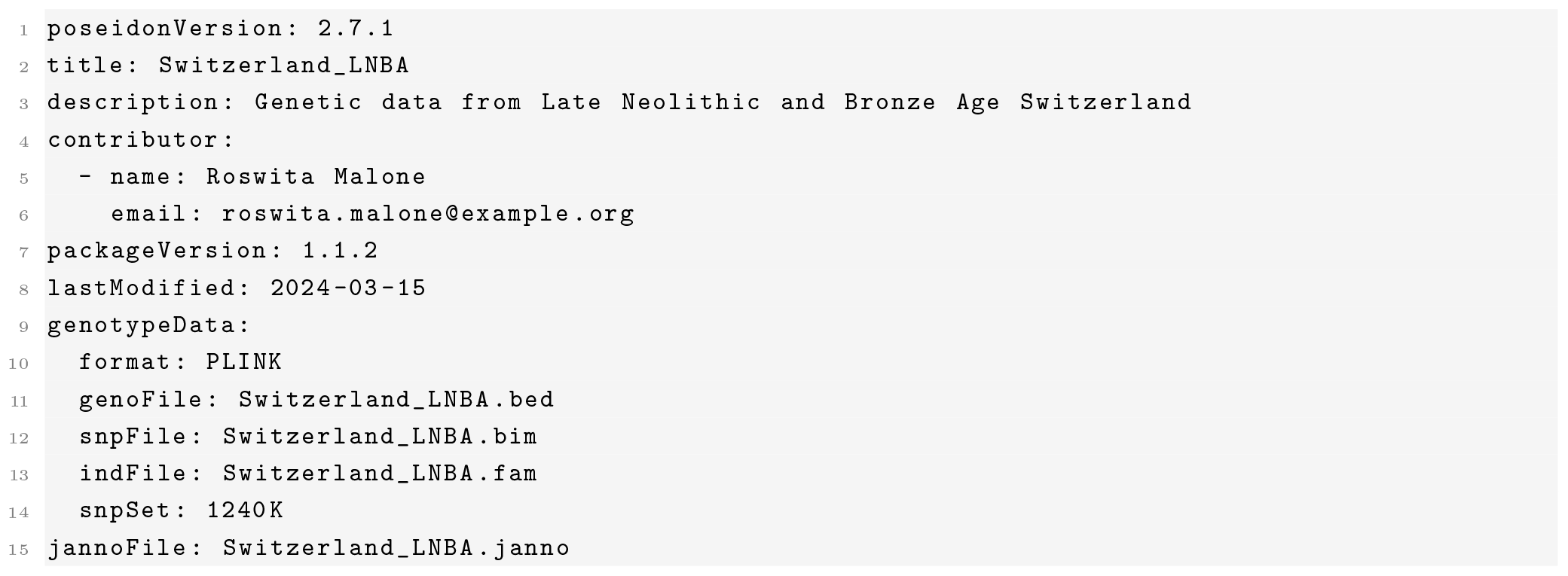
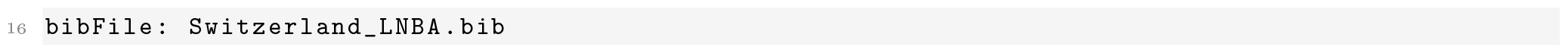

**Example 1:** A typical, neither minimal nor maximal, POSEIDON.yml file.

A Poseidon package is defined by a POSEIDON.yml file, using the YAML markup language (https://yaml.org) with a set of predefined fields, which stores some general information and, most importantly, relative paths to the other files in the package (see Example 1). A full list of the specified fields is provided as part of the schema, but only `poseidonVersion` (the schema version the package adheres to), `title` (a short package name), and `genotypeData` (context information for and relative paths to the genotype data files) are mandatory. Besides its function to define the package, the POSEIDON.yml also enables software to easily crawl for packages in a file system.

At the time of writing, Poseidon supports genotype data for single nucleotide polymorphisms (SNPs) in two common file formats: a binary-encoded format as introduced by the plink software package (PLINK) [22] and a plain-text, human-readable format defined for the eigensoft software tools (EIGENSTRAT) [17]. Both formats are structurally similar, and split into three files: A genotype table (.bed/.geno); a SNP file (.bim/.snp), defining the SNPs in the genotype table; and an individual file (.fam/.ind) for the respective samples. In the future,

Poseidon may add support for other formats, e.g. the more flexible vcf file format [23]. Arbitrary SNP sets are allowed and supported, but special keywords are reserved for the commonly used Affymetrix *HumanOrigins* array [24], and the so-called *1240k* in-solution hybridization capture reagent [25].

The .janno file accompanies the genotype data to provide context for each sample. It is designed as a tabular, tab-separated text file with a set of predefined columns. Each row corresponds to one entry in the individual file (.fam/.ind), featuring at least the following mandatory columns: `Poseidon_ID` (a sample identifier), `Genetic_Sex` (Female, Male, Unknown) and `Group_Name` (one or multiple group identifiers). Other, optional columns include standard information on the spatiotemporal provenience of a given sample, its genetic data quality metrics, and which major laboratory procedures it was subjected to. Supplementary Text 2 includes documentation for all specified columns of the .janno file. One particular column documents the scientific publication(s) within which the genetic or contextual data for a given sample was originally reported. This is a list column with BibTeX keys, which, in turn, must be specified in the .bib file of the Poseidon package, thus ensuring that each sample can be properly cited.

The .ssf file is similarly structured as the .janno file and stores sequencing source data, i.e. meta-information about the raw sequencing data behind the genotypes in a Poseidon package. The rows in this table correspond to sequencing entities, typically the set of unprocessed reads sequenced from DNA libraries or even multiple runs/lanes of the same library. At the time of writing, the specified columns mostly focus on how to download a given dataset from the INSDC databases such as ENA or SRA (https://www.insdc.org). The .ssf file does not have any mandatory variables, but entries can be linked to the package with the list column `poseidon IDs` in a many-to-many relationship.

### 4.2 Software tools

The following software tools were developed to work with Poseidon packages and other infrastructure of the Poseidon ecosystem. They can be considered reference implementations of the Poseidon schema, but are by no means exclusive nor comprehensive. The Poseidon schema is defined as an independent, versioned entity, and software developers can easily implement other tools in addition to what we as the Poseidon core team currently offer.

All Poseidon software is open-source (MIT-License) and available on GitHub (https://github.com/poseidon-framework). That is also where users should report issues.

#### 4.2.1 trident (v1.4.1.0)

*trident* is a command line software tool to create, download, inspect, merge, and subset Poseidon packages – and therefore the central tool of the Poseidon framework. It is implemented in Haskell as one executable for the *poseidon-hs* library and can be installed from source with the build systems cabal [26] or stack [27]. To ease installation we provide pre-compiled, static binary executables for Linux, Windows and macOS. We also maintain a package recipe under *poseidon-trident* on the bioconda channel [28], enabling installation of *trident* through the conda (https://docs.anaconda.com) package management system.

*trident* includes multiple sub-commands for different operations on and with Poseidon packages – Supplementary Text 3 gives a detailed overview including all specific arguments. It supports two mechanisms to obtain Poseidon packages: A user can create them from genotype data with `trident init` or download them from our community-maintained archives with `trident fetch`. The most involved and technically complex sub-command in *trident* is `trident forge`, which allows users to both subset and merge Poseidon packages. To simplify the process of package maintenance, `trident genoconvert` converts the genotype data between the formats Poseidon supports and `trident rectify` automates some common tasks such as checksum and version updates. For package inspection, *trident* includes `trident list`, which returns lists of packages, groups, or samples for a given package selection (either locally, or remotely by accessing the web API for the community-maintained online archives). `trident summarise` compiles some basic summary statistics about a package collection and `trident survey` gives an overview to which degree .janno files in a package collection are filled with data, i.e. the level of completeness for context information. The last and arguably the most important inspection subcommand is `trident validate`, which parses all components of a package to report violations of the Poseidon schema. This ensures the structural integrity of Poseidon packages and maintains machine-readability.

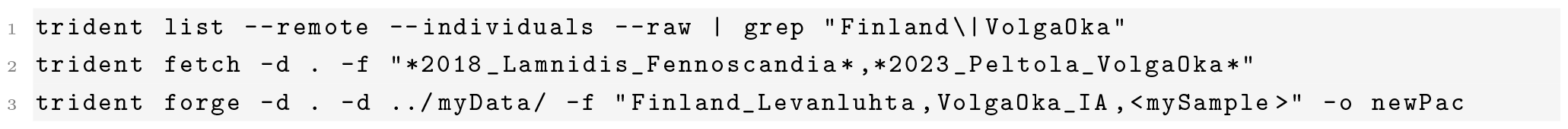

**Example 2:** A basic *trident* command line workflow to explore the community archive, download relevant packages and create a new package from the downloaded data, as well as a local/private data collection. `trident list` queries the webserver (`--remote`), and returns a tab separated (`--raw`) table of individuals/samples (`--individuals`) available in the public Community Archive (PCA). This list can then be filtered with standard command line tools (here `grep`). With `trident fetch` two packages are selected for download (`-f …`) into the current working directory (`-d .`). These two packages as well as an additional, local package collection (`-d ../myData/`) can then be read into `forge` to create a new package `newPac`, specifying (`-f …`) the groups `Finland Levanluhta`, `VolgaOka IA` and the local sample `mySample`.

A simple *trident* workflow could look like Example 2. The first two commands here require interaction with the Poseidon webserver. This server software is, in fact, also implemented in Haskell as a part of *poseidon-hs* and can be started with a hidden (and for most end users irrelevant) *trident* sub-command `trident serve`. On the client side `trident list` and `trident fetch` use this API and interact, by default, with the endpoints at https://server.poseidon-adna.org. A different server can be set with `--remoteURL`. More about the available endpoints below.

The `forge` sub-command is the technically most involved operation in *trident*. It discovers all Poseidon packages under a list of base directories (`-d`), reads them and their components into dedicated in-memory data structures, parses the query language in `-f` (or from a file with `--forgeFile`) to decide which entities (individuals, groups, packages) should be selected, and ultimately generates the new package including genotypes and context data in a single, low-memory stream processing run. All components of the Poseidon package format (i.e. the POSEIDON.yml file, the genotype data, the .janno, the .ssf, and the .bib file) are modelled in *poseidonhs* as algebraic data types with dedicated parsers. Corrupted files are rejected upon reading, including cross-file compatibility checks within each package (e.g. bibtex keys mentioned in the .janno file must be specified in the .bib file). The query language supported by `trident forge` in `-f` is a flexible domain-specific language (DSL) to describe arbitrary positive and negative entity selection scenarios. For the stream processing of genotype data in binary PLINK and EIGENSTRAT format, *poseidon-hs* relies on the the sequencing-formats library (https://github.com/stschiff/sequence-formats), which allows *trident* to read, filter and write data from a large amount of files at once, while also unifying SNPs to a consensus by recoding alleles on-the-fly. Varying SNP sets in multiple packages can be merged using either an intersection or a union operation.

#### 4.2.2 xerxes (v1.0.1.0)

*xerxes* is another command line software tool based on the *poseidon-hs* Haskell library. While *trident* is meant for data management, *xerxes* is intended for basic, every-day data analysis operations. Its most advanced and stable sub-command `xerxes fstats` makes extensive use of the genotype data stream processing introduced above to implement commonly used genome-wide statistics. These are F-Statistics (*F*_2_, *F*_3_ and *F*_4_, see Example 3, in several variants differing in bias-correction and normalisation) [24], *F*_ST_ [29], heterozygosity, and pairwise nucleotide mismatch rates for assessing pairwise relatedness. All statistics are evaluated with error-estimation using weighted block-Jackknife [30]. See Supplementary Text 4 for the *xerxes* user guide and Supplementary Text 5 for a whitepaper documenting the mathematical underpinnings of the implemented algorithms.

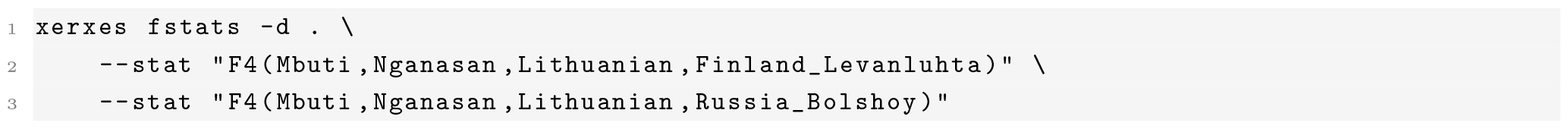

**Example 3:** Calculating two *F*_4_ statistics with *xerxes*, reproducing an analysis of Lamnidis et al. 2018 [31]. We assume the base directory (`-d .`) to include Poseidon packages with the groups `Mbuti`, `Nganasan`, `Lithuanian`,

`Finland_Levanluhta` and `Russia_Bolshoy`. These happen to be described in the Community Archive packages `2018_Lamnidis_Fennoscandia` [31], `2014_LazaridisNature` [3] and `2012_PattersonGenetics` [24], but xerxes will find those automatically if they are stored below the base directory. The individual statistics are specified with `--stats` and a dedicated domain specific language. For complex analyses *xerxes* offers a more powerful configuration file interface with the `--statConfig` argument.

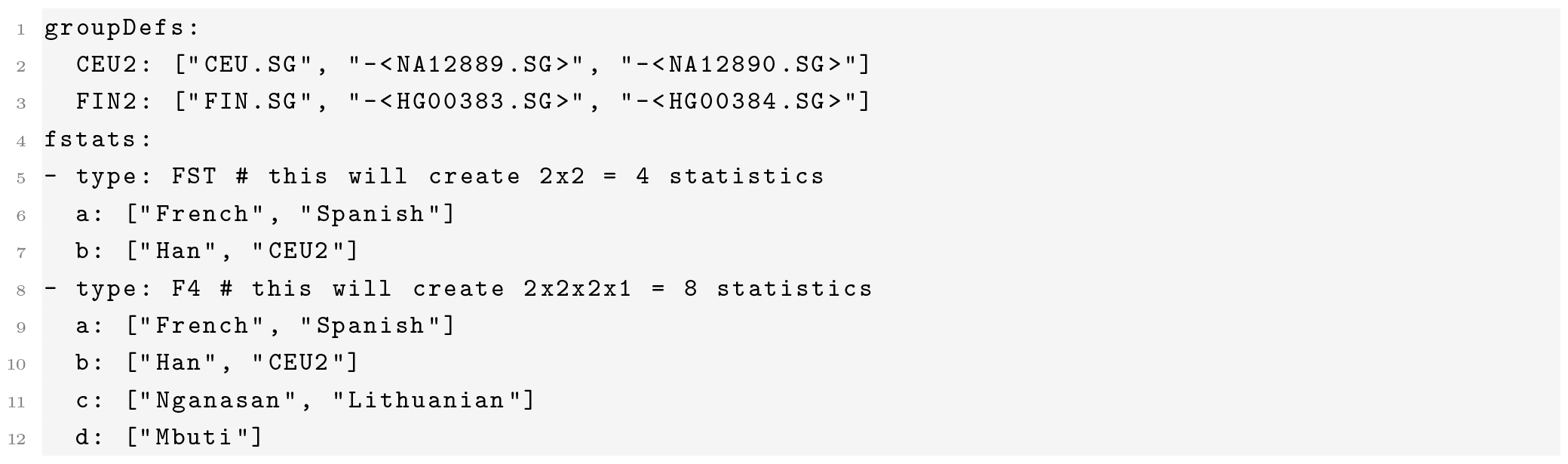

**Example 4:** An example of a configuration file for *xerxes*, specifying a total of 12 statistics of *F*_ST_ and *F*_4_ to be computed for all combinations of listed populations. In addition, the example features adhoc group definitions in the `groupDefs` section, which shows how to exclude individuals from a group using the minus-sign. More complex features of these configuration files include frequency-ascertainment, and selection of entire packages to act as groups.

We highlight three *xerxes* features that stand out: First, the user interface provides for a powerful way to specify families of statistics using combinatorical expansions. For example, users can specify in a configuration file (Example 4) that *F*_ST_ should be computed between all combinations of two lists of populations, and multiple such families can be requested simultaneously. Second, statistics can be easily adapted with custom group-definitions, in which individuals, groups or entire packages (similar to the `trident forge` selection language) can be excluded or included. If a specific individual turns out to be an outlier that should not be included in the computation of allele frequencies of its group, one can define a custom group and exclude that individual on the fly, without the need to change the underlying genotype definition files. Third, users do not have to specify the exact packages that their populations or individuals reside in, but can pass a base directory with dozens or hundreds of packages, and *xerxes* will automatically select only the relevant packages, involving the groups or individuals needed, thereby reducing memory- and run-time overhead: Similar to `trident forge`, *xerxes* uses stream-processing to merge the relevant genotype-files without the need to load large genotype matrices into memory. This is particularly important for dense genotyping data sets, for example involving tens of millions of SNPs as with the 1000 Genomes dataset [32].

#### 4.2.3 qjanno (v1.0.0.0)

*qjanno* is a command line tool implemented in Haskell to run SQL queries on .janno files (or arbitrary delimiter-separated text files) and therefore to conveniently subject the context data provided with Poseidon packages to a wide range of data transformations. It was built on top of a hard fork of v0.3.3 of the qsh package (https://github.com/itchyny/qhs, MIT-License). See Supplementary Text 6 for the *qjanno* user guide.

On startup, *qjanno* creates an SQLite [33] database in memory. It then reads the requested, well-structured text files, attributes each column a type and writes the contents of the files to tables in the in-memory database. It finally sends the user-provided SQL query to the database, waits for the result, parses it and returns it on the command line. The query gets pre-parsed to extract file names and then forwarded to an SQLite database server via the Haskell library sqlite-simple (https://github.com/nurpax/sqlite-simple). That means *qjanno* can parse and understand most SQLite3 syntax (see Example 5 and 6) with some minor exceptions (e.g. PRAGMA functions).

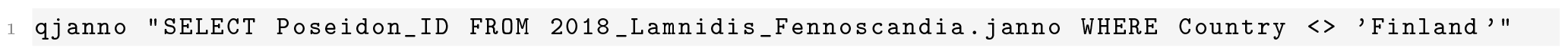

**Example 5:** A simple *qjanno* query on the command line. It extracts the columns `Poseidon_ID` and `Country` of the file `2018_Lamnidis_Fennoscandia.janno` and returns all rows where `Country` is not `‘Finland’`.

*qjanno* does not have a complete understanding of the .janno-file structure, and mostly treats it like a normal .tsv file. But .janno files are still given special consideration with a number of pseudo-functions, which allow to search Poseidon packages and .janno files recursively to load them together into one database table (see Example 6).

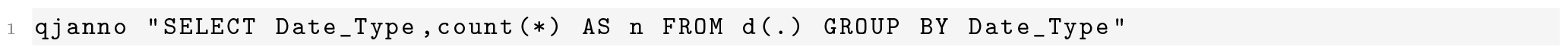

**Example 6:** Another *qjanno* query. With `‘FROM d(.)’` *qjanno* searches all latest versions of Poseidon packages under the current working directory, reads their .janno files and appends them to a common database table. This table is then grouped by the `Date_Type` column (*C14, contextual, modern*). The output features a row-count for each group in a new summary column `n`.

#### 4.2.4 janno R package (v1.0.0)

The *janno* R package simplifies loading and handling .janno files in R and the popular tidyverse [34] R package ecosystem. It provides a dedicated R S3 class `janno` that inherits from the `tibble` class to allow tidy reading and manipulating the context information in a Poseidon package (see Example 7). Supplementary Text 7 features its user guide.

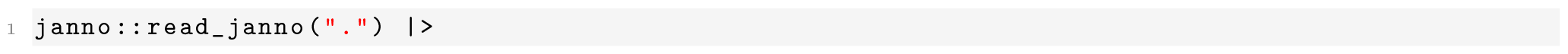
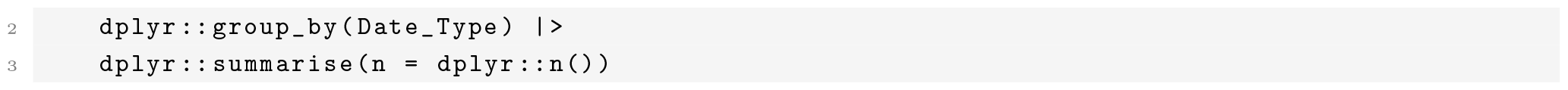

**Example 7:** A small R workflow enabled by the *janno* R package. The code is equivalent to the *qjanno* Example 6. `read_janno()` discovers and reads all .janno files under the current working directory into an object of classs `janno`. The dplyr verbs `group_by` and `summarise` can then be applied to this object to count the rows per `Date_Type` group.

As *trident* and *qjanno* (with `-d …`/`d(…)`), the package’s `read_janno()` function searches .janno files recursively under a given directory and loads them into a single data frame. The reading process includes validation according to the Poseidon schema. String list columns in .janno files are translated to true list columns in R. Beyond basic functionality (`print()`, `write_janno()`), the package features one more major function for `janno` objects: `process_age()`. This function processes the age information in the `Date_*` columns of the .janno file to derive a set of new columns useful for further chronological analysis via radiocarbon calibration: `Date_BC_AD_Prob` is a list column with the complete post-calibration probability distribution, `Date BC AD Median Derived` is the median of this distribution and `Date_BC_AD_Sample` stores *n* random samples drawn from it. The radiocarbon calibration is implemented with the *Bchron* R package [35].

### 4.3 Public archives

With a standardized, versioned data format, Poseidon does not require central infrastructure. Its software tools work independently and users can apply them locally on their own data. But Poseidon was also developed to mitigate the issue of increasingly tedious data preparation for future research projects. To that end, the Poseidon ecosystem includes three public, openly curated archives, which share implementation and infrastructure, but differ in their goals, mode of maintenance, and data content. See the Figures 4 and 5 for an overview of the current data content in the archives. Supplementary Text 8 explains this comparison in more detail.

#### 4.3.1 Technical infrastructure

All archives are hosted in dedicated Git repositories on GitHub (e.g. https://github.com/poseidon-framework/community-archive for the PCA), where each individual package is stored in a directory named after the package. This setup has a number of advantages: Git provides version control down to the individual lines of each meta- and context data file, Git and GitHub together include co-working features that allow users to submit new packages and suggest concrete changes to existing ones, and GitHub is a comparatively affordable host with advanced automation features. Git is by default not suitable for large, binary files, but GitHub offers large file storage with the Git LFS extension. The archives make use of this for all .bed and .bim files. Each change in the public archives is automatically validated using GitHub Actions on their cloud infrastructure, running a number of scripts on the new state, including `trident validate`, to ensure continuous structural integrity.

As already introduced above for `trident list --remote` and `trident fetch`, the data in the public archives is not just available on GitHub, but also and more conveniently via a web API provided by an open web server running `trident serve`. This service is hosted by the scientific IT service provider of the Max Planck Society (Gesellschaft für wissenschaftliche Datenverarbeitung mbH Göttingen, https://gwdg.de). Once per day the server fetches the latest changes to the archives on GitHub and incorporates new packages and package versions. It does so through a Git-integrated bookkeeping mechanism using the hidden *trident* subcommands `chronicle` and `timetravel`. Note that the server also provides outdated package versions (for the community archive starting from 2023-06-12) to maintain computational reproducibility. Here are the endpoints the server supports:

- https://server.poseidon-adna.org/packages returns a JSON list of all packages
- https://server.poseidon-adna.org/groups returns a JSON list of all groups
- https://server.poseidon-adna.org/individuals returns a JSON list of all samples/individuals
- https://server.poseidon-adna.org/zip_file/<package_name> returns a complete zip file of the package with the given name

The most important arguments for these are `?archive` to select the archive that should be queried, `?additionalJannoColumns` to add more detailed information to the `/individuals` response and `?package_version` to select a specific package version with `/zip_file`.

#### 4.3.2 The Community Archive

The Poseidon Community Archive (PCA) stores author-submitted, article-wise Poseidon packages. It focuses on packages prepared by the authors of the respective publication, containing the exact genotype data used for the paper, to ensure a maximum of computational reproducibility. Author submissions are also ideal for the context data in the .janno file, because the respective domain-experts are generally most knowledgeable on data quality and the spatiotemporal origin of their samples.

For historical reasons the PCA does not only contain author submissions, though. To kickstart the public archive development in 2020, we prepopulated it with packages derived from in-house data and previous versions of the AADR, which have since then been further modified and edited, as is transparent in the version history of these packages. This legacy data will remain in the PCA to maintain established workflows. Authors and other community members can take ownership, update entries if need be, and thus have the possibility to (further) improve the quality of these datasets. A contributing guide on the Poseidon webpage explain the details of how to submit a new paper-associated dataset or suggest changes to an existing one. Each submission passes through a checklist-based review process and is eventually confirmed by the Poseidon core team. Contributors and original and intermediate authors are credited via publication keys and corresponding .bib entries, as well as a dedicated `Contributor` list in the package-defining POSEIDON.yml file.

#### 4.3.3 The AADR Archive

The Poseidon AADR Archive (PAA) stores releases of the AADR dataset [13] reworked into Poseidon packages. It thus deviates from the PCA and the PMA in multiple important ways: It is not organized by individual publications, includes the versioning of the original provider on top of our own versioning, and relies to a lesser degree on community contributions. The cleaning and repackaging process is documented in an extra repository (https://github.com/poseidon-framework/aadr2poseidon) and mostly has the following goals: i) Creation of a version of the AADR that follows the Poseidon package standard and is thus directly compatible with `trident` and other Poseidon tooling, ii) increasing the machine-readability of the AADR, especially regarding the sample age information, and iii) providing clean .bib files with all references of publications in the AADR for convenient citation management.

#### 4.3.4 The Minotaur Archive

The Poseidon Minotaur Archive (PMA) mirrors the PCA in that it stores publication-wise packages, often the very same as the PCA. However, Packages in the PMA do not rely on author-submitted genotype data, but instead include genotypes consistently reprocessed from raw sequencing data, run through the Minotaur workflow (see below). The motivation for this bioinformatic reprocessing is to generate an internally consistent dataset, which optimises cross-package comparability, rather than per-author reproducibility of individual packages – like the AADR.

The submission of packages to the PMA is less direct as for the PCA and involves the preparation of a per-package recipe to parameterize the relevant processing run. Package recipes are archived in a dedicated GitHub repository, where targeted GitHub Actions guide users through the necessary steps to create and submit a new recipe.

### 4.4 The Minotaur workflow

The reproducibility of our processing behind the PMA is achieved with a semi-automatic computational workflow to generate Poseidon packages from raw sequencing data: the Minotaur workflow. The entry point for this processing pipeline is a *package recipe*, a collection of files containing all the information required to download, validate, and process the raw reads into a Poseidon package. Each recipe must contain an .ssf file, a .tsv file formatted like a valid input .tsv for nf-core/eager [36], a .config file outlining the nf-core/eager configuration parameters for processing, a .sh script that adapts the .tsv file to the local cluster at the time of processing, and finally a .txt file listing all the versions of scripts used when creating the recipe, to ensure reproducibility.

Contributors are able to request packages via GitHub issues on the dedicated minotaur-recipes repository (https://github.com/poseidon-framework/minotaur-recipes), and actively prepare package recipes by providing a configured .ssf file. This .ssf file gets validated through GitHub Actions, and then complemented to a full recipe with all the required files. Each recipe submission passes through a checklist-based review process and is finally confirmed by the Poseidon core team. This approach is designed to standardise and streamline the processing through the Minotaur workflow, while still allowing enough flexibility in its configuration to account for the heterogeneity present in raw sequencing data.

The minotaur-recipes repository is mirrored in the computational cluster of MPI-EVA, where the actual processing takes place. In the future, we may outsource this processing to a publicly accessible cloud service to make it fully independent from a particular institution. The raw data is downloaded there, and processed through nf-core/eager with the parameters specified in the .config file of the package recipe. By default, this processing includes adapter trimming and read-pair collapsing, aligning to the human reference, removal or PCR duplicates, masking of the ends of reads to mitigate aDNA damage artefacts, and genotyping. Changes to processing parameters are permitted, and can be specified, explained and recorded in the package recipe. The genotypes generated from this processing are then turned into a Poseidon package, whose .janno file is populated with descriptive statistics generated during the processing.

All the code responsible for this pipeline can be found at https://github.com/poseidon-framework/poseidon-eager. The resulting Poseidon package is finally uploaded to the PMA, where it again undergoes review before being added to the archive. During the review process, missing information for the package is filled in either manually, or pulled from the Community Archive, if appropriate.

## 5 Discussion

Archaeogenetic research, as many other fast growing, data-driven fields, is challenged by data heterogeneity, a lack of systematically applied standards, and from difficulties to discover published data. Poseidon addresses these issues on multiple levels: First, Poseidon provides a standardised, yet flexible data format for day-to-day scientific data analysis, large scale automation, and tidy data storage. Our design choice to build Poseidon around the notion of packages, together with integration of bibliography information (the .bib file), emphasises good citation practice, with any downstream merging necessarily resulting in complete bibliography files listing all original source papers. Second, the Poseidon software, such as *trident*, to discover, validate, update, merge, subset and analyse such packages, complements the standard to ease adoption. Third, perhaps the most ambitious component of Poseidon, our public archives, make use of the standard and its versioning feature to host published data and make it findable and transparently maintainable via GitHub community features.

This multi-layer architecture, with loosely coupled components, allows for a variety of adoption paths or starting points for users and data analysts: The package format alone can serve as a useful storage format for local work, even without using the software or archives. The software can help to create and work with such packages locally, even without the cloud-features and server-access. Finally, our archives can be seen as a transparent hub to download and discover data, even without our software or adoption of the package format. In light of our developing research field and its position between disciplines, we designed the Poseidon package definition to allow for flexibility, e.g. by allowing arbitrary non-schema columns, as for example used in the PAA to incorporate some fields specific to that data source. Furthermore, we have placed the standard definition itself on GitHub to enable its long-term use as a “living standard”, which can be modified and developed further given emerging use-cases and practice from the community, after open discussion and review.

While the Poseidon package format combines various standard formats (e.g. YAML, .tsv, EIGENSTRAT) into its own package specification, opportunities exist to integrate it with larger systems in the Linked Open Data (LOD) world [37]. For example, many of the concepts used in our meta-data definitions exist already in public ontologies, which could be more tightly integrated into our format in the future. Specifically, for example, our `Country` field in the .janno file definition could link to entities in Wikidata [38] or other comparable LOD databases. Our decision for a light-weight flat-file setup has given us leeway to adjust the system exactly to our preferences and the requirements of the field, but integration with the Web of Data thus remains an open tasks for the future. To find partners to establish this uplink in future versions Poseidon is part of the NFDI4Objects initiative (https://www.nfdi4objects.net).

Regarding infrastructure, Poseidon is currently very much dependent on GitHub. Following the example of other research data standards [39], all code, data, and issue-tracking is stored there, relying extensively on GitHub’s CI system (‘GitHub Actions’) for automatic code compilation and data validation. The lock-in into GitHub’s proprietary platform is slightly mitigated by the open Git format used for code- and data storage, but this is still a serious dependency, including factual drawbacks like a strict bandwidth limitation for large file data downloads.

Beyond these technical questions, finally, Poseidon is also exposed to some of the broader social challenges of scientific data management: Guiding the growth of a healthy community of developers, contributors and maintainers is not trivial. Poseidon currently depends on a small core team consisting of the authors of this paper. A growing number of active collaborators will require committing to a suitable governance system [40]. The work of the core team is currently funded by their employing institution, which also provides computational infrastructure and computing hours. If this commitment gets reduced in the middle-to long-term future, funding may become an increasingly pressing issue – a challenge shared by many research data management projects [41].

Critically the long-term success of Poseidon depends on scientists and generally practitioners in the field of archaeogenetics to reroute resources, not least time, into its development and maintenance, if they consider it valuable for their research and publications. With emerging initiatives like the HAAM Community (https://haam-community.github.io) and various working groups recently embracing Poseidon-based workflows and first papers referencing it explicitly ([42, 43] and other forthcoming work) we indeed see positive signals towards wider adoption. Regardless of whether this development subsists and a community forms around Poseidon, the Poseidon data format and the software developed for it will remain permanently and openly available for future reference.

## Supporting information

Supplementary Text 1 - Package specification

Supplementary Text 2 - Janno details

Supplementary Text 3- Trident

Supplementary Text 4 - Xerxes

Supplementary Text 5 - Xerxes Whitepaper

Supplementary Text 6 - Qjanno

Supplementary Text 7 - Janno R package

Supplementary Text 8 - Archive Comparison

## 6 Acknowledgements

This project has received funding from the Department of Archaeogenetics at the Max Planck Institute for Evolutionary Anthropology (MPI-EVA), the International Max Planck Research School for the Science of Human History at the Max Planck Institute for Geoanthropology (MPI-GEA), NFDI4Objects, so the the National research data infrastructure initiative by the German National Science foundation (DFG), and the European Research Council (ERC) under the European Union’s Horizon 2020 research and innovation program (grant agreement number 851511). The data processing with the Minotaur workflow heavily relies on computational facilities of MPI-EVA. We gratefully acknowledge insightful discussions with many current and former members of our department at MPI-EVA, most notably Selina Carlhoff, Luca Traverso, and Harald Ringbauer. Special thanks go to Michelle O’Reilly (MPI-GEA) who designed the Poseidon logo, specified the main colour palette for the website and revised the schematic overview Figures 1 and 2.

## Supplementary Texts

- Supplementary Text 1: Poseidon package specification v2.7.1
- Supplementary Text 2: .janno file details
- Supplementary Text 3: Guide for *trident* v1.4.1.0
- Supplementary Text 4: Guide for *xerxes* v1.0.1.0
- Supplementary Text 5: *xerxes* theoretical background
- Supplementary Text 6: Guide for *qjanno* v1.0.0.0
- Supplementary Text 7: Guide for the *janno* R package v1.0.0
- Supplementary Text 8: Comparison of the public archive content

