## Supplementary Text 1 - Package specification for "Poseidon – A framework for archaeogenetic human genotype data management"

### Poseidon Package Specification v2.7.1

PDF version of the standard at https://github.com/poseidon-framework/poseidon-schema
generated on 2024-03-19 based on the Git commit 904fb84

### I The Poseidon Standard v2.7.1

Poseidon is a solution for archaeogenetic genotype data organisation. This standard defines the core components of the Poseidon package.

A .pdf version of the latest instance of this document can be downloaded here.

Further details on genotype data, the .janno file and the .ssf file are documented on the Poseidon website.

The website also features a changelog documenting the changes across different schema versions here.

The key words *MUST*, *MUST NOT*, *REQUIRED*, *SHALL*, *SHALL NOT*, *SHOULD*, *SHOULD NOT*, *RECOMMENDED*, *MAY*, and *OPTIONAL* in this document are to be interpreted as described in RFC 2119.

### 1 The Poseidon package structure

A Poseidon package stores genotype data with context information for DNA samples from (ancient) (human) individuals. Packages are defined by the POSEIDON.yml file, which holds relative paths to all other files in a package.

A package therefore MUST contain:

- A `POSEIDON.yml` file to formally define the package
- Genotype data in PLINK or EIGENSTRAT format

It SHOULD additionally contain:

- A `.janno` file to store context information on spatiotemporal origin or sample quality
- A `.bib` file for literature references

It MAY also contain:

- A `README.md` file for arbitrary, additional context information
- A `CHANGELOG.md` file to document changes to the package
- A `.ssf` file with information on the underlying raw sequencing data

Here is an example of a package `Switzerland_LNBA_Roswita` in one directory:

```
Switzerland_LNBA_Roswita/POSEIDON.yml
Switzerland_LNBA_Roswita/Switzerland_LNBA.bed
Switzerland_LNBA_Roswita/Switzerland_LNBA.bim
Switzerland_LNBA_Roswita/Switzerland_LNBA.fam
Switzerland_LNBA_Roswita/Switzerland_LNBA.janno
Switzerland_LNBA_Roswita/Switzerland_LNBA.ssf
Switzerland_LNBA_Roswita/Switzerland_LNBA.bib
Switzerland_LNBA_Roswita/README.md
Switzerland_LNBA_Roswita/CHANGELOG.md
```

All text files in the package MUST be UTF-8 encoded.

### 2 The `POSEIDON.yml` file

The `POSEIDON.yml` file defines Poseidon packages by listing metainformation and relative paths in a standardised, machine-readable format.

- It MUST be a valid YAML file.
- Its mandatory and optional fields are documented in the POSEIDON_yml_fields.tsv file.

Here is an example for a `POSEIDON.yml` file:

```yaml
poseidonVersion: 2.7.1
title: Switzerland_LNBA_Roswita
description: LNBA Switzerland genetic data not yet published
contributor:
```

```yaml
  - name: Roswita Malone
   
    orcid: 1234-1234-1234-1234
  - name: Paul Panther
   
packageVersion: 1.1.2
lastModified: 2021-01-28
genotypeData:
  format: PLINK
  genoFile: Switzerland_LNBA_Roswita.bed
  genoFileChkSum: 95b093eefacc1d6499afcfe89b15d56c
  snpFile: Switzerland_LNBA_Roswita.bim
  snpFileChkSum: 6771d7c873219039ba3d5bdd96031ce3
  indFile: Switzerland_LNBA_Roswita.fam
  indFileChkSum: f77dc756666dbfef3bb35191ae15a167
  snpSet: 1240K
jannoFile : Switzerland_LNBA_Roswita.janno
jannoFileChkSum: 555d7733135ebcabd032d581381c5d6f
sequencingSourceFile: Switzerland_LNBA_Roswita.ssf
sequencingSourceFileChkSum: 19db1906240ee2f076e1a9659567dca4
bibFile: Switzerland_LNBA_Roswita.bib
bibFileChkSum: 70cd3d5801cee8a93fc2eb40a99c63fa
readmeFile: README.md
changelogFile: CHANGELOG.md
```

When a package is modified in any way (including updates of the context information in the `.janno` file), then the `packageVersion` field SHOULD be incremented and the `lastModified` field updated to the current date.

#### 2.1 Package versioning

The `packageVersion` field is a mandatory entry of the `POSEIDON.yml` file. It denotes the version of the individual package, using a three-component versioning system derived from semantic versioning.

Each version number is comprised of three numbers, separated by a `.`. For example: `0.1.0`, `1.0.0` or `2.1.3`. The first number gives the `Major`, the second the `Minor` and the third the `Patch` component of the version number. For a Poseidon package these components SHOULD be incremented when the following changes occur:

- `Major` (e.g. `1.4.2 -> 2.0.0`)
  - When samples are added to a package.
  - When samples are removed from a package.
  - When the genotype data (i.e. the contents of the `.bed`/`.bim`/`.fam` or `.geno`/`.snp`/`.ind` files) for any number of samples is changed.
- `Minor` (e.g. `1.4.2 -> 1.5.0`)
  - When larger pieces of meta- or context information are added or modified in any package file, except the

genotype data. For example:
    * An entire .janno, .bib or .ssf file is added or replaced.
    * Entire columns in the .janno or .ssf file are added or replaced.
    * Primary publications for samples in the .janno and .bib file are added or replaced.
- **Patch** (e.g. 1.4.2 -> 1.4.3)
  - When smaller pieces of meta- or context information are added or modified in any package file, except the genotype data. For example:
    * Individual entries in the .janno or .ssf file are added or replaced.
    * Secondary publications for samples in the .janno and .bib file are added or replaced.
    * BibTeX entries in the .bib file are modified.
    * The package **description** changes in the POSEIDON.yml file.
    * The CHANGELOG.md file is modified with additional information on previous entries.

When the **packageVersion** is changed, then the **lastModified** date MUST be updated and an entry to the CHANGELOG.md file SHOULD be added summarising the changes made.

Packages SHOULD start at **packageVersion** 0.1.0.

#### 3 Genotype data

Genotype data in Poseidon packages is stored either in (binary) PLINK or EIGENSTRAT format.

| | PLINK (binary) | EIGENSTRAT |
|---|---|---|
| genotype file | .bed (binary biallelic genotype table) | .geno (genotype file) |
| SNP file | .bim (extended MAP file) | .snp (snp file) |
| individual file | .fam (sample information) | .ind (indiv file) |

In addition to these files (and optionally their checksums), the POSEIDON.yml file SHOULD also provide a **snpSet** entry which determines the shape of the genotype file.

#### 4 The .janno file

The .janno file is a tab-separated text file with a header line. It holds context information (variables/columns) for each sample (objects/rows) in a package.

- A set of strictly defined core variables (defined by column name) and their possible content are documented here: janno_columns.tsv
- A .janno file MAY have all of these core variables, or only a subset of them.
- Only three columns MUST be present to make the file valid: **Poseidon_ID**, **Group_Name** and **Genetic_Sex**
- Arbitrary columns not defined here MAY be added as long as their column names do not clash with the defined ones.

- The column order is irrelevant.
- If information is unknown or a variable does not apply for a certain sample, then the respective cell(s) MAY be filled with `n/a` or simply an empty string.
- The order of the samples (rows) in the `.janno` file MUST be equal to the order in the genetic data files (`.ind`, `.fam`) in the package.
- The values in the columns **Poseidon_ID**, **Group_Name** and **Genetic_Sex** MUST be equal to the terms used in the genetic data files (`.ind`, `.fam`).
- Multiple predefined columns of the `.janno` file are list columns that can hold multiple values (either strings or numerics) separated by `;`.
- The decimal separator for all floating point numbers MUST be `.`.

#### 5 The `.bib` file

A BibTeX file with all references listed in the `.janno` file. The entry keys MUST fit the ones used in the `.janno` file.

Example:

```
@article{CassidyPNAS2015,
    doi = {10.1073/pnas.1518445113},
    url = {https://doi.org/10.1073%2Fpnas.1518445113},
    year = 2015,
    month = {dec},
    publisher = {Proceedings of the National Academy of Sciences},
    volume = {113},
    number = {2},
    pages = {368--373},
    author = {Lara M. Cassidy and Rui Martiniano and Eileen M. Murphy and Matthew D. Teasdale
 ↪  and James Mallory and Barrie Hartwell and Daniel G. Bradley},
    title = {Neolithic and Bronze Age migration to Ireland and establishment of the insular
 ↪  Atlantic genome},
    journal = {Proceedings of the National Academy of Sciences}
}
```

To connect a sample in the package to this particular literature reference, the .janno file column `Publication` would have to be filled with `CassidyPNAS2015`.

#### 6 The `README.md` file

A simple markdown file with informal, arbitrarily structured information accompanying the package.

Example:

```
This package contains a rather interesting set of samples relevant for the peopling of the
 ↪  Territory of Christmas Island in the Indian Ocean. We consider this especially relevant,
 ↪  because ...
```

#### 7 The `CHANGELOG.md` file

A markdown file to document changes in the history of a package.

Example:

```
- V 1.1.1: Fixed a spelling mistake in one site name: "Hosenacker" -> "Rosenacker"
- V 1.1.0: Added mtDNA contamination estimation to the .janno file
- V 1.0.0: Added spatial coordinates and age information to the .janno file and finalized a
↪ first stable version of the package
- V 0.2.0: Added previously restricted sample L1337
- V 0.1.0: Creation of the package
```

The structure with `- V X.X.X:` at the beginning of each line is not mandatory, but SHOULD be followed for reasons of interoperability.

#### 8 The `.ssf` file

The `.ssf` file is another tab-separated text file with a header line. It stores sequencing source data, so metainformation about the raw sequencing data behind the genotypes in a Poseidon package. The primary entities in this table are sequencing entities, typically corresponding to DNA libraries or even multiple runs/lanes of the same library.

- The predefined columns are specified here: ssf_columns.tsv
- All columns of this schema are optional, so a `.ssf` MAY have all of these core variables, only a subset of them, or even none. It SHOULD have a `poseidon_IDs` column, though, to link the sequencing entities to the Poseidon package.
- The link to the individuals listed in the `.janno`-file (and therefore to the entire Poseidon package) is made through a many-to-many foreign-key relationship between the .janno column `Poseidon_ID` and the .ssf column `poseidon_IDs`. That means each entry in the .janno file can be linked to many rows in the .ssf file and vice versa.
- As in the `.janno` file arbitrary columns not defined here MAY be added to the `.ssf` file as long as their column names do not clash with the defined ones.
- The order of columns and rows is irrelevant.
- If information is unknown or a variable does not apply, then the respective cell(s) MAY be filled with `n/a` or simply an empty string.
- Multiple predefined columns of the `.ssf` file are list columns that can hold multiple values (either strings or numerics) separated by `;`.
- The decimal separator for all floating point numbers MUST be `.`.

### II Appendix

The following tables specify individual fields/variables/columns in the `POSEIDON.yml`, the `.janno` and the `.ssf file`.

An asterisk `*` after the field name indicates a mandatory field that a given file MUST include to be valid.

#### 1 POSEIDON.yml file fields

POSEIDON.yml file fields

| Field | Description |
|---|---|
| poseidonVersion* | Poseidon package format version (e.g. 2.0.1)<br>type: String<br>format: X.Y.Z |
| title* | title of the package<br>type: String |
| description | description of the package (one or multiple sentences)<br>type: String |
| contributor | list of contributors to the package (not the publication author, but the Poseidon package creator)<br>type: Array |
| name* | name of one contributor<br>subfield of: contributor<br>type: String |
| email* | email of one contributor<br>subfield of: contributor<br>type: String<br>format: Email |
| orcid | orcid of one contributor<br>subfield of: contributor<br>type: String<br>format: ORCID |
| packageVersion* | package version (should be changed/incremented when the package is changed)<br>type: String<br>format: X.Y.Z |

POSEIDON.yml file fields *(continued)*

| Field | Description |
|---|---|
| lastModified | date of last modification of the package (should be updated when the package is changed)<br>type: Date<br>format: YYYY-MM-DD |
| genotypeData* | genotype data section |
| format* | genotype data file format<br>subfield of: genotypeData<br>type: String<br>format: EIGENSTRAT;PLINK |
| genoFile* | relative path to the geno file<br>subfield of: genotypeData<br>type: String<br>format: Path |
| genoFileChkSum | md5 checksum of the geno file<br>subfield of: genotypeData<br>type: String<br>format: md5 hash |
| snpFile* | relative path to the snp file<br>subfield of: genotypeData<br>type: String<br>format: Path |
| snpFileChkSum | md5 checksum of the snp file<br>subfield of: genotypeData<br>type: String<br>format: md5 hash |
| indFile* | relative path to the ind file<br>subfield of: genotypeData<br>type: String<br>format: Path |
| indFileChkSum | md5 checksum of the ind file<br>subfield of: genotypeData<br>type: String<br>format: md5 hash |

POSEIDON.yml file fields *(continued)*

| Field | Description |
|---|---|
| snpSet | SNP set in the genotype data<br>subfield of: genotypeData<br>type: String<br>format: 1240K;HumanOrigins;Other |
| jannoFile | relative path to the .janno file<br>type: String<br>format: Path |
| jannoFileChkSum | md5 checksum of the .janno file<br>type: String<br>format: md5 hash |
| sequencingSourceFile | relative path to the .ssf file<br>type: String<br>format: Path |
| sequencingSourceFileChkSum | md5 checksum of the .ssf file<br>type: String<br>format: md5 hash |
| bibFile | relative path to the .bib file<br>type: String<br>format: Path |
| bibFileChkSum | md5 checksum of the .bib file<br>type: String<br>format: md5 hash |
| readmeFile | relative path to the README file<br>type: String<br>format: Path |
| changelogFile | relative path to the CHANGELOG file<br>type: String<br>format: Path |

#### 2 .janno file variables

.janno file variables

| Variable | Description |
| --- | --- |
| Poseidon_ID* | sample identifier as defined by the genetics laboratory (e.g. I1234, BOT001), must fit to the values in the Poseidon package .fam/.ind file, must be unique within one package, if multiple datasets exist for the same individual different Poseidon_IDs are required<br>type: String |
| Genetic_Sex* | genetic sex of the individual derived from this sample, only F, M or U because the EIGENSTRAT and PLINK formats only support these three, edge cases (e.g. XXY, XYY, X0) are undefined and should be grouped as F, M or U, with a Note added<br>type: Char<br>allowed values: F; M; U |
| Group_Name* | meaningful population/group identifiers for the sample, should follow the geographic-temporal nomenclature proposed by Eisenmann et al. 2018 (https://doi.org/10.1038/s41598-018-31123-z), multiple entries separated by ;, the first value must be equal the group name in the .fam/.ind file<br>list column<br>type: String |
| Alternative_IDs | alternative identifiers for the same sampled individual, e.g. IDs in other databases or popular names like Ötzi/Iceman<br>list column<br>type: String |
| Relation_To | other samples (by Poseidon_ID) that are related/identical to this sample, multiple entries separated by ;<br>list column<br>type: String |
| Relation_Degree | relationship degree for relatives mentioned in Related_To, multiple values separated by ; in the same order as Related_To in case of multiple relations<br>list column<br>type: String<br>allowed values: identical; first; second; thirdToFifth; sixthToTenth; unrelated; other |

.janno file variables *(continued)*

| Variable | Description |
|---|---|
| Relation_Type | relationship type for relatives mentioned in Related_To (e.g. sister_of, child_of, nephew_of), multiple values separated by ; in the same order as Related_To in case of multiple relations<br>list column<br>type: String |
| Relation_Note | arbitrary comments about the genetic relationships of the sampled individual<br>type: String |
| Collection_ID | alternative sample identifier shared by the provider/owner of the sample (e.g. grave 40 skeleton 2)<br>type: String |
| Country | present-day political country of origin for the sample<br>type: String |
| Country_ISO | present-day political country expressed in ISO 3166-1 alpha-2 country codes<br>type: String |
| Location | unspecified location information for the sample, e.g. administrative or topographic region or mountains/rivers/lakes/cities nearby<br>type: String |
| Site | name of the archaeological site where the sample was found<br>type: String |
| Latitude | latitude where the sample was found with up to 5 places after the decimal point<br>type: Float<br>allowed range: -90 to 90 |
| Longitude | longitude with up to 5 places after the decimal point<br>type: Float<br>allowed range: -180 to 180 |

.janno file variables *(continued)*

| Variable | Description |
|---|---|
| Date_Type | type of dating information available for the sample, C14 if there is a set of radiocarbon dates in the columns Date_C14_Labnr, Date_C14_Uncal_BP and Date_C14_Uncal_BP_Err whose post-calibration probability distribution is a meaningful prior for the individual's year of death, contextual for any other age information only given in Date_BC_AD_Start, Date_BC_AD_Median and Date_BC_AD_Stop, "modern" for present-day individuals<br>type: String<br>allowed values: C14; contextual; modern |
| Date_C14_Labnr | lab numbers of C14 ages, multiple values separated by ; in case of multiple dates<br>list column<br>type: String |
| Date_C14_Uncal_BP | uncalibrated years BP (as in before 1950AD) for the C14 ages as reported by C14 labs, multiple values separated by ; in the same order as Date_C14_Labnr in case of multiple dates, only relevant if Date_Type is C14<br>list column<br>type: Integer<br>allowed range: 0 to Inf |
| Date_C14_Uncal_BP_Err | standard deviation (1-sigma ±) for the uncalibrated C14 ages as reported by the C14 labs, multiple values separated by ; in the same order as Date_C14_Labnr in case of multiple dates, only relevant if Date_Type is C14<br>list column<br>type: Integer<br>allowed range: 0 to Inf |
| Date_BC_AD_Start | lower (older) bound for the age of the sample in years BC/AD, negative numbers for BC, positive numbers for AD, in case of C14 dates 2-sigma post calibration interval, 2000 for modern samples<br>type: Integer<br>allowed range: -Inf to 2050 |

.janno file variables *(continued)*

| Variable | Description |
| --- | --- |
| Date_BC_AD_Median | median age of the sample in years BC/AD, for C14-dated samples median, for contextually dated samples simple mid-point of the archaeological intervals, 2000 for modern samples<br>type: Integer<br>allowed range: -Inf to 2050 |
| Date_BC_AD_Stop | upper (more recent) bound for the age of the sample in years BC/AD, counter point to Date_BC_AD_Start<br>type: Integer<br>allowed range: -Inf to 2050 |
| Date_Note | arbitrary comments about the dating information for the sample<br>type: String |
| MT_Haplogroup | mitochondrial haplogroup derived for the sample as specified on phylotree.org and as reported by the Haplofind or Haplogrep software tools<br>type: String |
| Y_Haplogroup | Y-chromosome haplogroup derived for the sample following a syntax with the main branch + the most terminal derived Y-SNP (e.g. R1b-P312)<br>type: String |
| Source_Tissue | skeletal element, tissue or other material sampled, the specific bone should be reported after an underscore (e.g. bone_phalanx), multiple values separated by ;<br>list column<br>type: String |
| Nr_Libraries | number of libraries produced for the sample<br>type: Integer |
| Library_Names | identifiers of the libraries used to generate the genotype data for the sample, multiple values separated by ;<br>list column<br>type: String |

| Variable | Description |
|---|---|
| Capture_Type | specifics of the data generation method (e.g. capture method) for the individual libraries generated for the sample, multiple values separated by ;<br>list column<br>type: String<br>allowed values: Shotgun; 1240K; ArborComplete; ArborPrimePlus; ArborAncestralPlus; TwistAncientDNA; OtherCapture; ReferenceGenome |
| UDG | udg treatment for the libraries, mixed in case multiple libraries with different UDG treatment were merged<br>type: String<br>allowed values: minus; half; plus; mixed |
| Library_Built | strandedness of the libraries, "mixed" in case multiple libraries with different protocols were merged<br>type: String<br>allowed values: ds; ss; mixed |
| Genotype_Ploidy | ploidy of the genotypes for the sample<br>type: String<br>allowed values: diploid; haploid |
| Data_Preparation_Pipeline_URL | url pointing to a description of the computational pipeline used to generate the genotype data from the source data<br>type: String |
| Endogenous | % endogenous DNA as estimated from SG libraries (before capture) as for example estimated by EAGER, not on target and no quality filter, in case of multiple libraries only the highest values should be reported<br>type: Float<br>allowed range: 0 to 100 |
| Nr_SNPs | number of non-missing SNPs for the sample, counted on the SNP-set stored in the Poseidon package<br>type: Integer |
| Coverage_on_Target_SNPs | average X-fold coverage across targeted SNP sites after quality filtering<br>type: Float |

.janno file variables *(continued)*

| Variable | Description |
| --- | --- |
| Damage | % damage on the 5' end for the main shotgun library used for sequencing and/or capture, in case of multiple libraries a value from the merged read alignment should be reported<br>type: Float<br>allowed range: 0 to 100 |
| Contamination | (modern) contamination of the sample as measured by the method in Contamination_Meas, multiple values separated by ; (for different methods, in case of multiple libraries report a value from the merged read alignment), the variables Contamination, Contamination_Err and Contamination_Meas must have the same number and order of (non-n/a) entries<br>list column<br>type: String |
| Contamination_Err | (modern) contamination estimate error of the sample<br>list column<br>type: String |
| Contamination_Meas | method to measure contamination, should be a software tool (ANGSD, Schmutzi, …) and the respective software versions, details should go to Contamination_Note<br>list column<br>type: String |
| Contamination_Note | arbitrary comments about the contamination estimation<br>type: String |
| Genetic_Source_Accession_IDs | ENA or SRA accession identifiers pointing to the source data used to generate the genotyping data for the sample, multiple values separated by ;, if multiple are given they should be arranged by descending specificity (e.g. project id > sample id > sequencing run id)<br>list column<br>type: String |
| Primary_Contact | project lead or first author who generated and published the data for the sample<br>type: String |

.janno file variables *(continued)*

| Variable | Description |
|---|---|
| Publication | bibtex keys for the publications where a sample was published (e.g. “AuthorJournalYear“) or “unpublished“, multiple values separated by ;, all must be present with complete BibTeX entries in the Poseidon package’s .bib file<br>list column<br>type: String |
| Note | arbitrary comments about the sample<br>type: String |
| Keywords | arbitrary tags, multiple values separated by ;<br>list column<br>type: String |

### 3 .ssf file variables

.ssf file variables

| Variable | Description |
|---|---|
| poseidon_IDs | Poseidon_IDs (in the .janno file) the sequencing entity corresponds to, multiple entries separated by ;<br>list column<br>type: String |
| udg | udg treatment applied to the library for the sequencing entity<br>type: String<br>allowed values: minus; half; plus |
| library_built | library preparation method applied for the sequencing entity (single- or double-stranded)<br>type: String<br>allowed values: ds; ss |
| sample_accession | sample accession code as used in the INSDC databases (including ENA and SRA) to identify the sequencing entity (e.g. SAMEA7050454)<br>type: String |
| study_accession | study accession code as used in the INSDC databases (e.g. PRJEB39316)<br>type: String |

.ssf file variables *(continued)*

| Variable | Description |
|---|---|
| run_accession | run accession code as used in the INSDC databases (e.g. ERR4331996), this should be a unique identifier in a Poseidon package<br>type: String |
| sample_alias | sample alias defined by the submitter in the raw sequencing data repository<br>type: String |
| secondary_sample_accession | a secondary sample accession, used in the ENA database for historical reasons (e.g. ERS4811084)<br>type: String |
| first_public | date (YYYY-MM-DD) the sequencing entity was first made public in the raw sequencing data repository<br>type: Date |
| last_updated | date (YYYY-MM-DD) the sequencing entity was last updated in the raw sequencing data repository<br>type: Date |
| instrument_model | name of the instrument used to process the sequencing entity (e.g. Illumina HiSeq 2500)<br>type: String |
| library_layout | library layout of the sequencing entity (e.g. SINGLE)<br>type: String |
| library_source | source of the DNA library (e.g. GENOMIC)<br>type: String |
| instrument_platform | platform, brand or type of the sequencer (e.g. ILLUMINA)<br>type: String |
| library_name | library identifier, so library name the submitter has entered to the raw sequencing data repository, data entries across which optical duplicates could exist should have matching library names<br>type: String |
| library_strategy | strategy used to create the library for the sequencing entity (e.g. WGS)<br>type: String |

.ssf file variables *(continued)*

| Variable | Description |
| --- | --- |
| fastq_ftp | ftp links to the FASTQ files for the sequencing entity in the raw sequencing data repository (e.g. ftp.sra.ebi.ac.uk/vol1/fastq/ERR433/009/ ERR4332639/ERR4332639.fastq.gz), multiple entries separated by ;<br>list column<br>type: URL |
| fastq_aspera | aspera links to the FASTQ files for the sequencing entity in the raw sequencing data repository (e.g. fasp.sra.ebi.ac.uk:/vol1/fastq/ERR433/009/ ERR4332639/ERR4332639.fastq.gz), multiple entries separated by ;<br>list column<br>type: URL |
| fastq_bytes | number of bytes in the FASTQ files, multiple entries separated by ;, must be in the same order as the ftp and/or aspera links<br>list column<br>type: Integer<br>allowed range: 0 to Inf |
| fastq_md5 | md5 hashes of the FASTQ files, multiple entries separated by ;, must be in the same order as the ftp and/or aspera links<br>list column<br>type: String |
| read_count | number of reads in the sequencing entity<br>type: Integer<br>allowed range: 0 to Inf |
| submitted_ftp | urls to the originally submitted files before they got converted to FASTQ in the INSDC databases, multiple entries separated by ;<br>list column<br>type: String |
