## Supplementary Text 2 - Janno details for "Poseidon – A framework for archaeogenetic human genotype data management"

### .janno file details

### Contents

**1 Background** **1**

**2 Identifiers** **1**

**3 Relations among samples/individuals** **2**

**4 Spatial position** **3**

**5 Temporal position** **3**

**7 Context information** **7**

#### 1 Background

The `.janno` file columns are specified in the Poseidon package specification here. The following documentation includes additional background information for many of the variables. This should make it more easy to compile the necessary information for both published and unpublished data. The `.pdf` version of the latest version of this document is available here.

#### 2 Identifiers

The `Poseidon_ID` column represents each sample with an ideally world-wide unique identifier string often equal to the identifier used in the respective accompanying publication. There is no central authority to issue these identifiers, so it remains in the hand of the authors to avoid duplication. We are aware of this inconsistency and hope the aDNA community will eventually come together to establish a mechanism to ensure uniqueness of identifiers. If there are multiple samples from one individual, then they have to be clearly distinguished with relevant suffixes added to the Poseidon_ID. Poseidon_IDs are also employed in the genetic data files in a Poseidon package and therefore have to adhere to certain constraints.

The column Alternative_IDs provides a way to list other IDs used for the respective individual. These might for example be names used in different publications or popular names like "Iceman", "Ötzi", "Girl of the Uchter Moor", "Tollund Man", etc.. The Relation_* columns described below allow to more precisely express the relationship type "identical" among samples in a Poseidon package.

The Collection_ID column stores an additional, secondary identifier as it is often provided by collaboration partners (archaeologists, museums, collections) that provide the specimen for archaeogenetic research. These identifiers can have a very heterogenous structure and may not be unique across different projects or institutions. The Collection_ID column is therefore a free-form text field.

The Group_Name column contains one or multiple group or population names for each individual, separated by ;. The first entry must be identical to the one used in the genotype data for the respective sample in a Poseidon package, and whitespace is not allowed in any of the entries. Assigning group and population names is a hard problem in archeogenetics [1], so the .janno file allows for more than one identifier.

### 3 Relations among samples/individuals

To systematically document biological relationships uncovered among samples/individuals in one or multiple Poseidon datasets (e.g. with software like READ [2] or BREADR [3]), the .janno file can be fit with a set of columns featuring the Relation_* prefix. Across these columns it should be possible to encode all kinds of pairwise, biological relationships an individual might have.

Relation_To is a string list column (so: multiple values are possible if separated by ;) that stores the Poseidon_IDs of other samples/individuals to which the current individual has some relationship.

Relation_Degree stores a formal description of the closeness of this relationship as measured purely from aDNA data. It is therefore also a list column that can hold the following values for each relationship:

- identical: The two samples are from the same individual or from identical twins
- first: The two individuals are closely related – a first degree relationship (e.g. siblings, parent-offspring)
- second: A second degree relationship (e.g. cousins, grandparent to grandchild)
- thirdToFifth: A third to fifth degree relationship (e.g. great-grandparent to great-grandchild)
- sixthToTenth: A sixth to tenth degree relationship
- unrelated: Unrelated – this is the default state among all individuals, which does not have to be expressed explicitly. This category will therefore probably never be used
- other: Any other kind of relationship not covered by the aforementioned categories

For each entry in Relation_To there must be a corresponding entry in Relation_Degree.

Relation_Type allows to add more verbose details about the relationship type, if it was possible to reconstruct that from the archaeological or historical context. Because there are too many possible permutations, there is no pre-defined set of values for what can and cannot be entered here. It is advisable, though, to stick to a general scheme like the following, which describes a given relationship from the point of view of the current individual:

- father_of: This individual is likely the father of the partner individual
- grandchild_of: This individual is likely the grandchild of the partner individual
- mother_or_daughter_of: This individual is likely either the mother or daughter of the partner individual (which might be unclear, in case of imprecise archaeological dating)

- `unknown`: The relationship is unclear or not yet determined. This is the default state and does not have to be expressed, unless multiple relationships are present and some but not all are known.
- ...

Unlike `Relation_Degree`, `Relation_Type` can be left empty even if there are entries in `Relation_To`. But if it is filled, then the number of values must be equal to the number of entries in both `Relation_To` and `Relation_Degree`.

The `Relation_Note` column allows to add free-form text information about the relationships of this individual. This might also include information about the method used to infer the degree and type.

### 4 Spatial position

The `.janno` file contains six columns to describe the spatial origin of an individual sample: `Country`, `Country_ISO`, `Location`, `Site` and finally `Latitude` and `Longitude`.

The `Country` column should contain a present-day political country name following the `English short name` in ISO 3166.

The `Country_ISO` column should contain the present-day political country of origin of the sample, expressed in codes using the standard ISO 3166-1 alpha-2 code, i.e. "AR" for Argentina or "NO" for Norway.

The `Location` column allows for free-form text entry and can contain further, unspecified location information. This might be the name of an administrative or geographic region, or an arbitrary unit of reference like a mountain, lake or city close to the point of discovery of the respective sample.

The `Site` column should contain a site name, ideally in the latin alphabet and ideally the name that is commonly used in publications.

The `Latitude` and `Longitude` columns should contain geographic coordinates (WGS84) in decimal degrees (DD) with a precision of not more than five places after the decimal point. This yields a precision of about 1.1132m at the equator which is sufficient to describe the position of an archaeological site. Coordinates in other formats like for example Degrees Minutes Seconds (DMS) or in completely different coordinate reference systems should be transformed. There exist many open source software solutions to do that, most based on the PROJ library e.g. the The World Coordinate Converter.

### 5 Temporal position

The temporal position of a sample is encoded with seven different columns in the `.janno` file: `Date_C14_Labnr`, `Date_C14_Uncal_BP`, `Date_C14_Uncal_BP_Err`, `Date_BC_AD_Median`, `Date_BC_AD_Start`, `Date_BC_AD_Stop`, `Date_Type`.

#### 5.1 General structure

The `Date_Type` column handles the general distinction between the most common forms of age information:

- `modern`: Applies to present-day reference samples, so not ancient DNA.
- `C14`: Applies if there is a set of radiocarbon dates explicitly listed in the columns `Date_C14_Labnr`, `Date_C14_Uncal_BP` and `Date_C14_Uncal_BP_Err` whose post-calibration probability distribution is a meaningful prior for the individual's year of death. The dates do not always have to be directly from the

individual's tissue, but they should be immediately relevant for their year of death (e.g. a date from a grain kernel recovered from the individual's grave).

- `contextual`: Applies in all other cases if the columns `Date_BC_AD_Median`, `Date_BC_AD_Start`, `Date_BC_AD_Stop` can be filled. This includes age attribution based on the archaeologically determined stratigraphy or typological information. `contextual` should also be chosen if the sample is dated very indirectly with radiocarbon dating (e.g. radiocarbon dates from other, unrelated features of the same site) or dated with other physical or chemical dating methods (e.g. dendrochronology or optically stimulated luminescence).

So `Date_C14_Labnr`, `Date_C14_Uncal_BP` and `Date_C14_Uncal_BP_Err` only go along with `Date_Type = C14`, whereas `Date_BC_AD_Median`, `Date_BC_AD_Start`, `Date_BC_AD_Stop` complement both `Date_Type = C14` and `Date_Type = contextual`. Radiocarbon dates that only serve as secondary evidence for a contextual dating should NOT be reported in `Date_C14_Labnr`, `Date_C14_Uncal_BP` and `Date_C14_Uncal_BP_Err`.

#### 5.2 The columns in detail

Each radiocarbon date has a unique identifier: the "lab number". It consists of a lab code issued by the journal Radiocarbon for each laboratory and a serial number. This lab number makes the date well identifiable and should be reported in `Date_C14_Labnr` with the lab code separated from the serial number with a minus symbol.

The uncalibrated radiocarbon measurement can be described by a Gaussian distribution with mean and standard deviation. So the column `Date_C14_Uncal_BP` holds the mean of that distribution in years before present (BP) as usually reported by radiocarbon laboratories. The age is always a positive integer value starting from a zero that corresponds to 1950 AD. The column `Date_C14_Uncal_BP_Err` holds the respective standard deviation for each date in years. This should be the 1-sigma distance, so that the probability that the actual uncalibrated age of the measured sample is within the `Date_C14_Uncal_BP`±`Date_C14_Uncal_BP_Err` range is about 68%.

`Date_C14_Labnr`, `Date_C14_Uncal_BP` and `Date_C14_Uncal_BP_Err` each can hold multiple values separated by `;` to allow for multiple radiocarbon dates for each aDNA sample. With multiple values the number and order of values in the columns must be consistent.

In the columns `Date_BC_AD_Median`, `Date_BC_AD_Start`, `Date_BC_AD_Stop` ages are reported in years BC and AD, so in relation to the zero point of the Gregorian calender. BC dates are represented with negative, AD with positive integer values.

- If radiocarbon dates are available (`Date_Type = C14`): `Date_BC_AD_Median` should report the median age after calibration. With multiple dates this can be determined either with sum calibration or more complex (e.g. bayesian) age modelling. `Date_BC_AD_Start` and `Date_BC_AD_Stop` should report the starting/ending age of a 95% probability window around the age median.
- If only contextual (e.g. from archaeological typology) age information is available (`Date_Type = contextual`): `Date_BC_AD_Start` and `Date_BC_AD_Stop` should simply report the approximate start and end date determined by the respective source of scientific authority (e.g. an archaeologist knowledgable about the relevant typological sequences). In this case `Date_BC_AD_Median` should be calculated as the mean of `Date_BC_AD_Start` and `Date_BC_AD_Stop` rounded to an integer value.
- If the sample is a modern reference sample (`Date_Type = modern`): `Date_BC_AD_Median`, `Date_BC_AD_Start`, `Date_BC_AD_Stop` should all be set to the value 2000, for 2000 AD.

The column `Date_Note` stores arbitrary free-form text information about the dating of a sample.

### 6 Genetic summary data

#### 6.1 Individual properties

The `Genetic_Sex` column should encode the biological sex as determined from the DNA read distribution on the X and Y chromosome. It only allows for the entries

- `F`: female
- `M`: male
- `U`: unknown

This limitation stems from the genotype data formats by Plink and the Eigensoft software package. Edge cases (e.g. XXY, XYY, X0, ...) can not be expressed with this format and should be reported as `U` with an additional comment in the free text `Note` field. Genetic sex determination for ancient DNA can be performed for example with Sex.DetERRmine [4].

The `MT_Haplogroup` column is meant to store the human mitochondrial DNA haplogroup for the respective individual in a simple string. The entry can be arbitrarily precise. A software tool to determine the MT haplogroup is for example Haplogrep [5].

The `Y_Haplogroup` column holds the respective human Y-chromosome DNA haplogroup in a simple string. To avoid confusion from using different haplotype naming systems, the notation should follow a syntax with the main branch + the most terminal derived Y-SNP separated with a minus symbol (e.g. R1b-P312), similar to that used by Yfull.

#### 6.2 Library properties

The `Source_Tissue` column documents the skeletal, soft tissue or other elements from which source material for DNA library preparation was extracted. If multiple samples have been taken from different elements, these can be listed separated by `;`. Specific bone names should be reported with an underscore (e.g. bone_phalanx, tooth_molar).

The `Nr_Libraries` column holds a simple integer value of the number of libraries that have been prepared for an individual.

The `Library_Names` column should list the names for the libraries as used in the publication, separated by `;`.

The `Capture_Type` column specifies the general pre-sequencing preparation methods that have been applied to the library. See [6] for a review of the different techniques (not including newer developments). This field can hold one of multiple different values, but also multiple of these separated by `;` if different methods have been applied for different libraries.

- `Shotgun`: Sequencing without any enrichment (whole genome sequencing, screening etc.).
- `1240k`: Target enrichment with hybridization capture optimised for sequences covering the 1240k SNP array [7], [8], [9].
- `ArborComplete`, `ArborPrimePlus`, `ArborAncestralPlus`: Target enrichment with hybridization capture as provided by Arbor Biosciences in three different kits branded myBaits Expert Human Affinities.
- `TwistAncientDNA`: Target enrichment with hybridization capture as provided by Twist Bioscience [10].
- `OtherCapture`: Target enrichment with hybridization capture for any other set of sequences.
- `ReferenceGenome`: Modern reference genomes where aDNA fragmentation is not an issue and other sample preparation techniques apply.

The UDG column documents if the libraries for the respective individual went through UDG (or USER enzyme) treatment. This wet lab protocol step removes molecular damage in the form of deaminated cytosines characteristic of ancient DNA.

- minus: A protocol without UDG treatment (e.g. [11]).
- half: A protocol with UDG-half treatment (e.g. [12]).
- plus: A protocol with UDG-full treatment (e.g. [13]).
- mixed: Multiple libraries that went through different UDG treatment approaches, and whose data were later merged.

The Library_Built column describes the library preparation method regarding single- or double-stranded protocols. See e.g. [14] for more information.

- ds: Double-stranded library preparation.
- ss: Single-stranded library preparation.
- mixed: If multiple libraries with different strandedness were combined. See also the Sequencing Source File in the Poseidon package as a way to provide details.

The Genotype_Ploidy column stores whether the genotype calls for this individual are originally haploid or diploid. Even for diploid organisms, it is often useful to represent genotypes by single haploid alleles (so-called pseudo-haploid genotypes), for example to generate relatively unbiased genotype calls from low coverage data. Because both the PLINK and EIGENSTRAT genotyping formats always *encode* genotype calls as diploid (by "doubling" the pseudo-haploid genotypes), the information on the original Ploidy of the call gets lost. This column is therefore used to record the underlying calling procedure. This becomes important, for example, when sample sizes are queried to compute bias-correction factors when computing F-Statistics or FST. The Genotype_Ploidy column can contain one of the following values:

- diploid: True diploid genotype calls were made.
- haploid: Haploid genotypes were called and then doubled.

The column Data_Preparation_Pipeline_URL should finally store an URL that links to a complete and human-readable description of the computational pipeline (for example a specific configuration for nf-core/eager [15]) by which the sample data was processed.

#### 6.3 Data yield

The Endogenous column holds the percentage of mapped reads over the total amount of reads that went into the mapping pipeline. That boils down to the DNA percentage of the library that matches the (human) reference. It should be determined from Shotgun libraries (so before any hybridization capture), not on target (i.e. across the whole genome, not specific positions), and before any mapping quality filtering. In case of multiple libraries only the highest value should be reported. The % endogenous DNA can be calculated for example with the endorS.py script.

The Nr_SNPs column gives the number of SNPs reported in the genotype data files for this individual.

The Coverage_on_Target_SNPs column reports the mean fold coverage on the SNP set of the genotype dataset (e.g. 1240K) for the merged libraries of this sample. To calculate the coverage it is necessary to determine which SNPs are covered how many times by the mapped reads. Individual SNPs might be covered multiple times, whereas others may not be covered at all by the highly deteriorated ancient DNA. The coverage for each SNP is therefore a number between 0 and n. The statistic can be determined for example with the QualiMap [16] software package. In case of multiple libraries, the total coverage should be given across all libraries.

#### 6.4 Data quality

The Damage column contains the % damage on the first position of the 5' end for the main Shotgun library used for sequencing or capture. This is an important statistic to verify the age of ancient DNA. In case of multiple libraries you should report a value from the merged read alignment.

##### 6.4.1 Contamination

Contamination of ancient DNA with foreign reads is a major challenge for archaeogenetics. There exist multiple competing ideas, algorithms and software tools to estimate the degree of contamination for individual samples (e.g. ANGSD [17], contamLD [18] or hapCon [19]), with some methods only applicable under certain circumstances (e.g. popular X-chromosome based approaches only work on male individuals). Also the results of different methods tend to differ both in the degree of contamination they estimate and in the way the output is usually encoded. To cover the multitude of methods in this domain, and to make the results representable in the `.janno` file, we offer the `Contamination_*` column family.

`Contamination` is a list column to represent the different contamination values estimated for a sample with one or multiple software tools. As usual multiple values are separated by `;`.

`Contamination_Err` is another list column to store the respective (standard) error term for the values in `Contamination`.

Some tools for contamination estimation do not return a mean plus a standard error. ContamMix, for example, yields a 95% confidence interval instead, to better represent assymetric output distributions. `Contamination` and `Contamination_Err` can not represent this. We suggest to derive a mean and a standard error from these alternative outputs. The latter can be calculated as the largest distance from the mean to the limits of the confidence interval.

`Contamination_Meas` finally is the third necessary list column, which contextualizes the values in `Contamination` and `Contamination_Err`. Each measure in these columns has to be accompanied by the software and software version used to calculate it. The individual entries might e.g. look like this:

- `ANGSD v0.935`
- `hapCon v0.4a1`
- `custom script`

This setup has the consequence that the columns `Contamination`, `Contamination_Err`, `Contamination_Meas` always have to have the same number of `;`-separated values.

The `Contamination_Note` column is a free text field to add additional information about the contamination estimates, e.g. which parameters where used with the respective software tools.

### 7 Context information

The `Genetic_Source_Accession_IDs` column was introduced to link the derived genotype data in Poseidon with the raw sequencing data typically uploaded to archives like the ENA [20] or SRA [21]. There, projects and individual samples are given clear unique identifiers: Accession IDs. This janno column is supposed to store one or multiple of these Accessions IDs for each individual/sample in Poseidon. If multiple are entered, then they should be arranged by descending specificity from left to right (e.g. project id > sample id > sequencing run id).

The Primary_Contact column is a free-form text field that stores the name of the main or the corresponding author of the respective paper for published data.

The Publication column holds either the value unpublished for (yet) unpublished samples or – for published data – one or multiple citation-keys of the form AuthorJournalYear without any spaces or special characters. These keys have to be identical to the BibTeX citation-keys identifying the respective entries in the .bib file of the package. BibTeX is a file format to store bibliographic information, where each entry (article, book, website, ...) is defined by a series of parameters (authors, year of publication, journal, ...). Here's an example .bib file with two entries for [22] and [23]:

```
@article{CassidyPNAS2015,
    doi = {10.1073/pnas.1518445113},
    url = {https://doi.org/10.1073%2Fpnas.1518445113},
    year = 2015,
    month = {dec},
    publisher = {Proceedings of the National Academy of Sciences},
    volume = {113},
    number = {2},
    pages = {368--373},
    author = {Lara M. Cassidy and Rui Martiniano and Eileen M. Murphy and
              Matthew D. Teasdale and James Mallory and Barrie Hartwell
              and Daniel G. Bradley},
    title = {Neolithic and Bronze Age migration to Ireland and establishment
             of the insular Atlantic genome},
    journal = {Proceedings of the National Academy of Sciences}
}

@article{FeldmanScienceAdvances2019,
    doi = {10.1126/sciadv.aax0061},
    url = {https://doi.org/10.1126%2Fsciadv.aax0061},
    year = 2019,
    month = {jul},
    publisher = {American Association for the Advancement of Science ({AAAS})},
    volume = {5},
    number = {7},
    pages = {eaax0061},
    author = {Michal Feldman and Daniel M. Master and Raffaela A. Bianco and
              Marta Burri and Philipp W. Stockhammer and Alissa Mittnik and
              Adam J. Aja and Choongwon Jeong and Johannes Krause},
    title = {Ancient {DNA} sheds light on the genetic origins of early Iron Age
             Philistines},
    journal = {Science Advances}
}
```

The string CassidyPNAS2015 is the citation-key of the first entry. To cite both publications in the Publication column, one would enter CassidyPNAS2015;FeldmanScienceAdvances2019.

When creating a new Poseidon package the .bib file should be filled together with the Publication column.

One of the most simple ways to obtain the BibTeX entries may be to request them with the doi from the doi2bib wep app. It could be necessary to adjust the result manually, though. The citation-key, for example, has to be replaced by the one used in the `Publication` column.

The `Note` column is a free-form text field that can contain small amounts of additional information that is not yet expressed in a more systematic form in the the other `.janno` file columns.

The `Keywords` column was introduced to allow for tagging individuals with arbitrary keywords. This should simplify sorting and filtering in personal Poseidon package repositories. Each keyword is a string and multiple keywords can be separated with `;`.

Arbitrary additional columns can be included in a `.janno` file, but they should be named in a way that they do not conflict with the Poseidon package specification. These columns will not be validated (assumed free-form text), but they will be preserved in the Poseidon package, and propagated during operations with `trident forge`.

---

299 [14] M.-T. Gansauge and M. Meyer, “Single-stranded DNA library preparation for the sequencing of ancient or damaged DNA,” *Nature Protocols*, vol. 8, no. 4, pp. 737–748, Mar. 2013, doi: 10.1038/nprot.2013.038.

300 [15] J. A. Fellows Yates *et al.*, “Reproducible, portable, and efficient ancient genome reconstruction with nf-core/eager,” *PeerJ*, vol. 9, p. e10947, Mar. 2021, doi: 10.7717/peerj.10947.

301 [16] K. Okonechnikov, A. Conesa, and F. García-Alcalde, “Qualimap 2: Advanced multi-sample quality control for high-throughput sequencing data,” *Bioinformatics*, vol. 32, no. 2, pp. 292–294, Oct. 2015, doi: 10.1093/bioinformatics/btv566.

302 [17] T. S. Korneliussen, A. Albrechtsen, and R. Nielsen, “ANGSD: Analysis of next generation sequencing data,” *BMC Bioinformatics*, vol. 15, no. 1, Nov. 2014, doi: 10.1186/s12859-014-0356-4.

303 [18] N. Nakatsuka, É. Harney, S. Mallick, M. Mah, N. Patterson, and D. Reich, “ContamLD: Estimation of ancient nuclear DNA contamination using breakdown of linkage disequilibrium,” *Genome Biology*, vol. 21, no. 1, Aug. 2020, doi: 10.1186/s13059-020-02111-2.

304 [19] Y. Huang and H. Ringbauer, “hapCon: Estimating contamination of ancient genomes by copying from reference haplotypes,” *Bioinformatics*, vol. 38, no. 15, pp. 3768–3777, Jun. 2022, doi: 10.1093/bioinformatics/btac390.

305 [20] J. Burgin *et al.*, “The European Nucleotide Archive in 2022,” *Nucleic Acids Research*, vol. 51, no. D1, pp. D121–D125, Nov. 2022, doi: 10.1093/nar/gkac1051.

306 [21] K. Katz, O. Shutov, R. Lapoint, M. Kimelman, J. R. Brister, and C. O’Sullivan, “The Sequence Read Archive: A decade more of explosive growth,” *Nucleic Acids Research*, vol. 50, no. D1, pp. D387–D390, Nov. 2021, doi: 10.1093/nar/gkab1053.

307 [22] L. M. Cassidy *et al.*, “Neolithic and Bronze Age migration to Ireland and establishment of the insular Atlantic genome,” *Proceedings of the National Academy of Sciences*, vol. 113, no. 2, pp. 368–373, Dec. 2015, doi: 10.1073/pnas.1518445113.

308 [23] M. Feldman *et al.*, “Ancient DNA sheds light on the genetic origins of early Iron Age Philistines,” *Science Advances*, vol. 5, no. 7, p. eaax0061, Jul. 2019, doi: 10.1126/sciadv.aax0061.
