## Supplementary Text 3- Trident for "Poseidon – A framework for archaeogenetic human genotype data management"

### Guide for trident v1.4.1.0

#### 1 Installation

See the Poseidon website (https://www.poseidon-adna.org/#/trident) or the GitHub repository (https://github.com/poseidon-framework/poseidon-hs) for up-to-date installation instructions.

### 2 The trident CLI

Trident is a command line software tool structured in multiple subcommands. If you installed it properly you can call it on the command line by typing `trident`. This will show an overview of the general options and all subcommands, which are explained in detail below.

```
Usage: trident [--version] [--logMode MODE | --debug] [--errLength INT]
               [--inPlinkPopName MODE] (COMMAND | COMMAND)

  trident is a management and analysis tool for Poseidon packages. Report issues
  here: https://github.com/poseidon-framework/poseidon-hs/issues

Available options:
  -h,--help                Show this help text
  --version                Show version number
  --logMode MODE           How information should be reported: NoLog, SimpleLog,
                           DefaultLog, ServerLog or VerboseLog.
                           (default: DefaultLog)
  --debug                  Short for --logMode VerboseLog.
  --errLength INT          After how many characters should a potential error
                           message be truncated. "Inf" for no truncation.
                           (default: CharCount 1500)
  --inPlinkPopName MODE    Where to read the population/group name from the FAM
                           file in Plink-format. Three options are possible:
                           asFamily (default) | asPhenotype | asBoth.

Package creation and manipulation commands:
  init                     Create a new Poseidon package from genotype data
  fetch                    Download data from a remote Poseidon repository
  forge                    Select packages, groups or individuals and create a
                           new Poseidon package from them
  genoconvert              Convert the genotype data in a Poseidon package to a
                           different file format
  jannocoalesce            Coalesce information from one or multiple janno files
                           to another one
  rectify                  Adjust POSEIDON.yml files automatically to package
                           changes

Inspection commands:
  list                     List packages, groups or individuals from local or
                           remote Poseidon repositories
  summarise                Get an overview over the content of one or multiple
                           Poseidon packages
  survey                   Survey the degree of context information completeness
                           for Poseidon packages
  validate                 Check Poseidon packages or package components for
```

```
                             structural correctness
```

trident allows to work directly with genotype data (see -p below), but it is optimized for the interaction with Poseidon packages, which wrap and contextualize the data. Most trident subcommands therefore have a central parameter, called --baseDir or simply -d to specify one or more base directories to look for packages. For example, if all Poseidon packages live inside a repository at /path/to/poseidon/packages you would simply say trident <subcommand> -d /path/to/poseidon/dirs/ and trident would automatically search all subdirectories inside of the repository for valid Poseidon packages (as identified by valid POSEIDON.yml files).

You can arrange a Poseidon repository in a hierarchical way. For example:

```
/path/to/poseidon/packages
    /modern
        /2019_poseidon_package1
        /2019_poseidon_package2
    /ancient
        /...
        /...
    /Reference_Genomes
        /...
        /...
```

This structure then allows to select only the level of packages you are interested in, even individual ones. -d can be given multiple times, which is particularly useful as you may have your own data to co-analyse with external reference data. In this case you simply need to provide your own genotype data as yet another Poseidon package to be added to your trident command. For example, you may have genotype data in EIGENSTRAT format (trident supports EIGENSTRAT and PLINK as formats.):

```
~/my_project/my_project.geno
~/my_project/my_project.snp
~/my_project/my_project.ind
```

Then you can transform that into a skeleton Poseidon package with the init command. You can also do it manually by simply adding a POSEIDON.yml file, with, for example, the following content:

```yaml
poseidonVersion: 2.7.1
title: My_awesome_project
description: Unpublished genetic data from my awesome project
contributor:
  - name: Stephan Schiffels
   
packageVersion: 0.1.0
lastModified: 2020-10-07
genotypeData:
  format: EIGENSTRAT
  genoFile: my_project.geno
  snpFile: my_project.snp
  indFile: my_project.ind
```

```yaml
jannoFile: my_project.janno
bibFile: sources.bib
```

Two remarks: 1) All file paths in this POSEIDON.yml file are considered *relative* to the directory in which POSEIDON.yml resides. For this example we assume that this file is added into the same directory as the three genotype files. 2) Besides the genotype data files there are two (technically optional) files referenced in this example: `sources.bib` and `my_project.janno`. Of course you can add them manually - `init` automatically creates empty dummy versions.

Once you have set up your own Poseidon package (which is really only a skeleton so far), you can add it to your `trident` analysis, by simply adding your project directory to the command using `-d`, for example:

```bash
trident list -d /path/to/poseidon/packages/modern \
  -d /path/to/poseidon/packages/ReferenceGenomes \
  -d ~/my_project \
  --packages
```

#### 2.1 General notes

##### 2.1.1 Logging and command line output

For all subcommands the general argument `--logMode` defines how `trident` reports messages (to stderr) on the command line:

- *NoLog*: Hides all messages.
- *SimpleLog*: Plain and simple output.
- *DefaultLog*: Adds the severity indicators (log levels) `Info`, `Warning` and `Error` before each message. This is the default setting.
- *ServerLog*: Additionally adds timestamps before each message.
- *VerboseLog*: Shows not just messages on the log levels `Info`, `Warning` and `Error` like the other modes, but also on the more verbose level `Debug`. Use this mostly relevant for debugging.

`--debug` is short for `--logMode VerboseLog` to activate this important log level more easily.

##### 2.1.2 Package duplicates and versions

- For `trident` multiple packages in a set of base directories can share the same `title`, if they have different `packageVersion` numbers. If the version numbers are also identical or missing, then `trident` stops with an exception.
- The `trident` subcommands `genoconvert`, `list`, `rectify`, `survey` and `validate` by default consider all versions of each Poseidon package in the given base directories. The `--onlyLatest` flag causes them to instead only consider the latest versions.
- `fetch` and `forge` generally consider all package versions. Their selection language (see below) allows for detailed version handling.
- `summarize` and `jannocoalesce` consider always only the latest package versions.

##### 2.1.3 Individual/Sample duplicates

- `Poseidon_IDs` (so individual/sample names) within one package have to be unique, or `trident` will stop.

- We also discourage sample duplicates across packages in package repositories, but `trident` will generally continue with them. `validate` will fail though, if the `--ignoreDuplicates` flag is not set.
- `forge` offers a special mechanism to resolve sample duplicates within its selection language.

###### 2.1.4 Group names in `.fam` files

The `.fam` file of PLINK-formatted genotype data is used inconsistently across different popular aDNA software tools to store group/population name information. The (global) `trident` option `--inPlinkPopName` with the arguments `asFamily` (default), `asPhenotype` and `asBoth` allows to control the reading of the population name from PLINK `.fam` files. The subcommands that write genotype data (`forge`, `genoconvert`) have a corresponding option `--outPlinkPopName` to specify this for the output.

###### 2.1.5 Whitespaces in the `.janno` file

While reading the `.janno` file `trident` trims all leading and trailing whitespaces around individual cells. Also all instances of the `No-Break Space` unicode character will be removed. This means these whitespaces will not be preserved when a package is `forge`d.

### 3 Package creation and manipulation commands

#### 3.1 Init command

`init` creates a new Poseidon package from genotype data files. It adds a POSEIDON.yml file, a dummy `.janno` file for context information and an empty `.bib` file for literature references.

Command line details

```
Usage: trident init ((-p|--genoOne FILE) | --inFormat FORMAT --genoFile FILE
                      --snpFile FILE --indFile FILE) [--snpSet SET]
                    (-o|--outPackagePath DIR) [-n|--outPackageName STRING]
                    [--minimal]

  Create a new Poseidon package from genotype data

Available options:
  -h,--help                Show this help text
  -p,--genoOne FILE        One of the input genotype data files. Expects .bed,
                           .bim or .fam for PLINK and .geno, snp or .ind for
                           EIGENSTRAT. The other files must be in the same
                           directory and must have the same base name.
  --inFormat FORMAT        The format of the input genotype data: EIGENSTRAT or
                           PLINK. Only necessary for data input with --genoFile
                           + --snpFile + --indFile.
  --genoFile FILE          Path to the input geno file.
  --snpFile FILE           Path to the input snp file.
  --indFile FILE           Path to the input ind file.
  --snpSet SET             The snpSet of the package: 1240K, HumanOrigins or
                           Other. Only relevant for data input with -p|--genoOne
```

```
                              or --genoFile + --snpFile + --indFile, because the
                              packages in a -d|--baseDir already have this
                              information in their respective POSEIDON.yml files.
                              (default: Other)
  -o,--outPackagePath DIR  Path to the output package directory.
  -n,--outPackageName STRING
                              The output package name. This is optional: If no name
                              is provided, then the package name defaults to the
                              basename of the (mandatory) --outPackagePath
                              argument. (default: Nothing)
  --minimal                   Should the output data be reduced to a necessary
                              minimum and omit empty scaffolding?
```

The command

```
trident init \
  --inFormat EIGENSTRAT/PLINK \
  --genoFile path/to/genoFile \
  --snpFile path/to/snpFile \
  --indFile path/to/indFile \
  --snpSet 1240K|HumanOrigins|Other \
  -o path/to/new_package_name
```

requires the format (`--inFormat`) of your input data (either `EIGENSTRAT` or `PLINK`), the paths to the respective files (`--genoFile`, `--snpFile`, `--indFile`), and optionally the "shape" of these files (`--snpSet`), so if they cover the `1240K`, the `HumanOrigins` or an `Other` SNP set.

A simpler interface is available with `-p (+ --snpSet)`, which only requires a path to one of the genotype data files and automatically discovers the others if they share the same base name:

```
trident init \
  -p path/to/genoFile \
  --snpSet 1240K|HumanOrigins|Other \
  -o path/to/new_package_name
```

The following file extensions are expected:

| | EIGENSTRAT | PLINK |
|---|---|---|
| genoFile | `.geno` | `.bed` |
| snpFile | `.snp` | `.bim` |
| indFile | `.ind` | `.fam` |

The output package created by `init` is located in a new directory `-o`, which should not already exist when `init` is called, and gets the package title corresponding to the basename of `-o`. You can also set the title explicitly with `-n`.

The `--minimal` flag causes `init` to create a minimal package with a very basic POSEIDON.yml and no `.bib` and `.janno` files.

#### 3.2 Fetch command

`fetch` allows to download Poseidon packages from a remote Poseidon server via a Web API. This server provides all packages in the Poseidon public archives.

Command line details

```
Usage: trident fetch (-d|--baseDir DIR)
                     (--downloadAll |
                       (--fetchFile FILE | (-f|--fetchString DSL)))
                     [--remoteURL URL] [--archive STRING]

  Download data from a remote Poseidon repository

Available options:
  -h,--help                Show this help text
  -d,--baseDir DIR         A base directory to search for Poseidon packages.
  --downloadAll            Download all packages the server is offering.
  --fetchFile FILE         A file with a list of packages. Works just as -f, but
                           multiple values can also be separated by newline, not
                           just by comma. -f and --fetchFile can be combined.
  -f,--fetchString DSL     List of packages to be downloaded from the remote
                           server. Package names should be wrapped in asterisks:
                           *package_title*. You can combine multiple values with
                           comma, so for example: "*package_1*, *package_2*,
                           *package_3*". fetchString uses the same parser as
                           forgeString, but does not allow excludes. If groups
                           or individuals are specified, then packages which
                           include these groups or individuals are included in
                           the download.
  --remoteURL URL          URL of the remote Poseidon server.
                           (default: "https://server.poseidon-adna.org")
  --archive STRING         The name of the Poseidon package archive that should
                           be queried. If not given, then the query falls back
                           to the default archive of the server selected with
                           --remoteURL. See the archive documentation at
                           https://www.poseidon-adna.org/#/archive_overview for
                           a list of archives currently available from the
                           official Poseidon Web API. (default: Nothing)
```

It works with

```
trident fetch -d ... -d ... \
  -f "*package_title_1*,*package_title_2-1.0.1*,group_name,<individual1>"
```

and the entities you want to download must be listed either in a simple string of comma-separated values, which can be passed via `-f`/`--fetchString`, or in a text file (`--fetchFile`). Entities are then combined from these sources.

Entities are specified using a special syntax (see also the documentation of `forge` below): packages are wrapped in asterisks, with or without a version number appended after a dash (e.g. `*package_title*` or `*package_title-1.2.3`), group names are spelled as is, and individual names are wrapped in angular brackets (e.g. `<individual1>`). Fetch will figure out which packages need to be downloaded to include all specified entities.

`--downloadAll`, which can be given instead of `-f` and `--fetchFile`, causes fetch to download all packages from the server. The downloaded packages are added in the first (!) `-d` directory (which gets created if it doesn't exist), but downloads are only performed if the respective packages are not already present in the latest version in any of the `-d` directories.

Note that `trident fetch` is usually used in a workflow with `trident list --remote`: First one inspects what is available on the server with `list`, to then compile a custom, targeted `fetch` command.

`fetch` has the optional arguments `--remote https:://..."` to name an alternative Poseidon server and `--archive` to select a specific Poseidon archive on the server.

#### 3.3 Forge command

`forge` creates new Poseidon packages by extracting and merging packages, populations and individuals/samples from Poseidon repositories.

Command line details

```
Usage: trident forge ((-d|--baseDir DIR) |
                       ((-p|--genoOne FILE) | --inFormat FORMAT --genoFile FILE
                         --snpFile FILE --indFile FILE) [--snpSet SET])
                     [--forgeFile FILE | (-f|--forgeString DSL)]
                     [--selectSnps FILE] [--intersect] [--outFormat FORMAT]
                     [--minimal] [--onlyGeno] (-o|--outPackagePath DIR)
                     [-n|--outPackageName STRING] [--packagewise]
                     [--outPlinkPopName MODE]

  Select packages, groups or individuals and create a new Poseidon package from
  them

Available options:
  -h,--help                Show this help text
  -d,--baseDir DIR         A base directory to search for Poseidon packages.
  -p,--genoOne FILE        One of the input genotype data files. Expects .bed,
                           .bim or .fam for PLINK and .geno, .snp or .ind for
                           EIGENSTRAT. The other files must be in the same
                           directory and must have the same base name.
  --inFormat FORMAT        The format of the input genotype data: EIGENSTRAT or
                           PLINK. Only necessary for data input with --genoFile
                           + --snpFile + --indFile.
  --genoFile FILE          Path to the input geno file.
  --snpFile FILE           Path to the input snp file.
  --indFile FILE           Path to the input ind file.
  --snpSet SET             The snpSet of the package: 1240K, HumanOrigins or
```

```
                         Other. Only relevant for data input with -p|--genoOne
                         or --genoFile + --snpFile + --indFile, because the
                         packages in a -d|--baseDir already have this
                         information in their respective POSEIDON.yml files.
                         (default: Other)
--forgeFile FILE         A file with a list of packages, groups or individual
                         samples. Works just as -f, but multiple values can
                         also be separated by newline, not just by comma.
                         Empty lines are ignored and comments start with "#",
                         so everything after "#" is ignored in one line.
                         Multiple instances of -f and --forgeFile can be
                         given. They will be evaluated according to their
                         input order on the command line.
-f,--forgeString DSL     List of packages, groups or individual samples to be
                         combined in the output package. Packages follow the
                         syntax *package_title*, populations/groups are simply
                         group_id and individuals <individual_id>. You can
                         combine multiple values with comma, so for example:
                         "*package_1*, <individual_1>, <individual_2>,
                         group_1". Duplicates are treated as one entry.
                         Negative selection is possible by prepending "-" to
                         the entity you want to exclude (e.g. "*package_1*,
                         -<individual_1>, -group_1"). forge will apply
                         excludes and includes in order. If the first entity
                         is negative, then forge will assume you want to merge
                         all individuals in the packages found in the baseDirs
                         (except the ones explicitly excluded) before the
                         exclude entities are applied. An empty forgeString
                         (and no --forgeFile) will therefore merge all
                         available individuals. If there are individuals in
                         your input packages with equal individual id, but
                         different main group or source package, they can be
                         specified with the special syntax
                         "<package:group:individual>".
--selectSnps FILE        To extract specific SNPs during this forge operation,
                         provide a Snp file. Can be either Eigenstrat (file
                         ending must be '.snp') or Plink (file ending must be
                         '.bim'). When this option is set, the output package
                         will have exactly the SNPs listed in this file. Any
                         SNP not listed in the file will be excluded. If
                         option '--intersect' is also set, only the SNPs
                         overlapping between the SNP file and the forged
                         packages are output. (default: Nothing)
--intersect              Whether to output the intersection of the genotype
                         files to be forged. The default (if this option is
```

```
                           not set) is to output the union of all SNPs, with
                           genotypes defined as missing in those packages which
                           do not have a SNP that is present in another package.
                           With this option set, the forged dataset will
                           typically have fewer SNPs, but less missingness.
  --outFormat FORMAT       The format of the output genotype data: EIGENSTRAT or
                           PLINK. (default: PLINK)
  --minimal                Should the output data be reduced to a necessary
                           minimum and omit empty scaffolding?
  --onlyGeno               Should only the resulting genotype data be returned?
                           This means the output will not be a Poseidon package.
  -o,--outPackagePath DIR  Path to the output package directory.
  -n,--outPackageName STRING
                           The output package name. This is optional: If no name
                           is provided, then the package name defaults to the
                           basename of the (mandatory) --outPackagePath
                           argument. (default: Nothing)
  --packagewise            Skip the within-package selection step in forge. This
                           will result in outputting all individuals in the
                           relevant packages, and hence a superset of the
                           requested individuals/groups. It may result in better
                           performance in cases where one wants to forge entire
                           packages or almost entire packages. Details: Forge
                           conceptually performs two types of selection: First,
                           it identifies which packages in the supplied base
                           directories are relevant to the requested forge, i.e.
                           whether they are either explicitly listed using
                           *PackageName*, or because they contain selected
                           individuals or groups. Second, within each relevant
                           package, individuals which are not requested are
                           removed. This option skips only the second step, but
                           still performs the first.
  --outPlinkPopName MODE   Where to write the population/group name into the FAM
                           file in Plink-format. Three options are possible:
                           asFamily (default) | asPhenotype | asBoth. See also
                           --inPlinkPopName.
```

`forge` can be used with

```
trident forge -d ... -d ... \
  -f "*package_name*, group_id, <individual_id>" \
  -o path/to/new_package_name
```

where the entities (packages, groups/populations, individuals/samples) you want in the output package can be denoted either as a string on the command line (`-f`/`--forgeString`), or in an input text file (`--forgeFile`). See the section below for the syntax of this selection language. Do not forget to wrap the `--forgeString` query in quotes.

Including one or multiple Poseidon packages with `-d` is not the only way to include data for a forge operation. It is also possible to consider unpackaged genotype data directly with `-p (+ --snpSet)` or `--inFormat + --genoFile + --snpFile + --indFile (+ --snpSet)`. This makes the following example possible, where we merge data from one Poseidon package and two unpackaged genotype datasets to get a new EIGENSTRAT dataset.

```
trident forge \
  -d 2017_GonzalesFortesCurrentBiology \
  -p 2018_VeeramahPNAS/2018_VeeramahPNAS.fam \
  --inFormat PLINK \
  --genoFile 2017_HaberAJHG/2017_HaberAJHG.bed \
  --snpFile 2017_HaberAJHG/2017_HaberAJHG.bim \
  --indFile 2017_HaberAJHG/2017_HaberAJHG.fam \
  -f "<STR241.SG>,<ERS1790729.SG>,Iberia_HG.SG" \
  -o testpackage \
  --outFormat EIGENSTRAT \
  --onlyGeno
```

###### 3.3.1 The forge selection language

The text in `--forgeString` and `--forgeFile` (and with a reduced syntax also in `--fetchString` and `--fetchFile`) are parsed as a domain specific query language that describes precisely which entities should be compiled in the output package of a given `forge` operation. The language has multiple syntactic elements and a specific evaluation logic.

In general a `--forgeString` query consists of multiple entities, separated by `,`. The main entities are Poseidon packages, groups/populations and individuals/samples:

- Each package title is surrounded by `*`: `*package*`. That means if you want all individuals of the Poseidon package `2019_Jeong_InnerEurasia` in the output package you would add `*2019_Jeong_InnerEurasia*` to the query.
- Groups/populations are not specially marked: `group`. So to get all individuals of the group `Swiss_Roman_Period`, you would simply add `Swiss_Roman_Period`.
- Individuals/samples are surrounded by `<` and `>`: `<individual>`. `ALA026` therefore becomes `<ALA026>`. A second way to denote individuals is with the more verbose and specific syntax `<package:group:individual>`. Such defined individuals take precedence over differently defined ones (so directly with `<individual>` or as a subset of `*package*` or `group`). This allows to resolve duplication issues precisely – at least in cases where the duplicated individuals differ in source package or primary group.
- Package versions can be appended to package names, such as `*package-1.2.3*`.
- This also works with the verbose individual syntax: `<package-1.2.3:group:individual>`.

In the `--forgeFile` each line is treated as a separate forgeString, empty lines are ignored and `#` symbols start comments. So this is a valid example of a forgeFile:

```
# Packages
*pac1*, *pac2-1.2.3*
```

```
# Groups and individuals from other packages beyond pac1 and pac2
group1, <ind1>, group2, <ind2>, <pac3:group3:ind3>

# pac2 has two outlier individuals that should be ignored
-<ind4>             # This one has very low coverage
-<pac2:group4:ind5> # This one is from a different time period
```

By prepending `-` to entities, we can exclude them from the forged package (this feature is not available for `fetch`). `forge` figures out the final list of samples to include by interpreting all forge-entities in order. So an entity list `*pac1*,-<ind1>,group1` may result in a different outcome than `*pac1*,group1,-<ind1>`, depending on whether `<ind1>` belongs to `group1` or not.

If the forge entity list starts with a negative entity, or if the entity list is empty, `forge` will implicitly assume you want to include all individuals in all **latest** versions of packages found in the base directories (except the ones explicitly excluded, of course).

The specific semantics of the various ways to include or exclude entities are as follows:

###### 3.3.1.1 Inclusion queries

- `*pac1*`: Select all individuals in the latest version of package “pac1”
- `*pac1-1.0.1*`: Select all individuals in package “pac1” with version “1.0.1”
- `group1`: Select all individuals associated with “group1” in all latest versions of all packages
- `<ind1>`: Select the individual named “ind1”, searching in all latest packages.
- `<pac1:group1:ind1>`: Select the individual named “ind1” associated with “group1” in the latest version of package “pac1”
- `<pac1-1.0.1:group1:ind1>`: Select the individual named “ind1” associated with “group1” in the package “pac1” with version “1.0.1”

###### 3.3.1.2 Exclusion queries

- `-*pac1*`: Remove all individuals in all versions of package “pac1”
- `-*pac1-1.0.1*`: Remove only individuals in package “pac1” with version “1.0.1” (but leave other versions in)
- `-group1`: Remove all individuals associated with “group1” in all versions of all packages (not just the latest)
- `-<ind1>`: Remove all individuals named “ind1” in all versions of all packages (not just the latest)
- `-<pac1:group1:ind1>`: Remove the individual named “ind1” associated with “group1”, searching in all versions of package “pac1”
- `-<pac1-1.0.1:group1:ind1>`: Remove the individual named “ind1” associated with “group1”, but only if they are in “pac1” with version “1.0.1”

If a query results in multiple individuals with the same name, forge will throw an error.

##### 3.3.2 Treatment of the genotype data while merging

Forge performs a series of steps to merge the genotype data of multiple source files:

1. Genotype data from each package is streamed in parallel. Because our packages may have different SNP locations (specified by chromosome-position pairs) listed in their `.bim/.snp` file, we first perform a zipping-

operation, whose behaviour depends on whether `--intersect` is set or not. Without `--intersect`, any SNP position listed in any package will be forwarded to the output, with missing values being filled in in all packages that do not list that particular SNP. With `--intersect`, only SNP positions that are present in all packages are considered. Note that relevant for this step is only whether a given SNP position is part of the genotype data, not whether the actual genotypes are missing or not.

2. At each SNP, the consensus alleles are selected, by collecting all reference and alternative alleles from all sources. If more than two non-dummy alleles (alleles different from `N`) are present in that collection, an error is thrown. If exactly two non-dummy alleles are present (which should be the case for binary SNPs), the two alleles are declared "reference" and "alternative" alleles for the output. If only one non-dummy allele is present, it is set to be the reference allele, and "N" is set to be the alternative.
3. All source genotype data is then read and recoded in terms of the two chosen consensus alleles. This will make sure that source data with flipped reference and alternative allele gets correctly merged in.
4. SNP IDs, as part of PLINK `.bim` files are checked across the source files. If all SNP IDs for a given SNP are missing, then the result will also be missing. If there is only one SNP ID present in some or all source packages, that ID gets forwarded to the output. In the (unusual) case that there are multiple different non-missing SNP IDs (of the form "rs" followed by a number), then a debug warning is output (which gets printed to the screen when `--debug` is selected), and simply the first value is chosen to be output into the forged `.bim` file. We decided not to throw an error in that case, because we consider the physical position of the SNP (specified by Chromosome and position) to be definitive, and the SNP ID to be of secondary importance.
5. Genetic positions, as part of PLINK `.bim` files are checked in a similar manner, with "0.0" being interpreted as missing.

##### 3.3.3 Treatment of the `.janno` file while merging

`forge` merges and subsets `.janno` files along with the genotype data. If a package lacks a `.janno` file, then a basic one will be created internally an on-the-fly based on the information in the genotype data, and used for the output. Missing columns across packages will be filled with `n/a`.

For merging two `.janno` files **A** and **B** the following rules apply regarding undefined, arbitrary additional columns:

- If **A** has an additional column which is not in **B** then empty cells in the rows imported from **B** are filled with `n/a`.
- If **A** and **B** share additional columns with identical column name, then they are treated as semantically identical units and merged accordingly.
- In the resulting `.janno` file, all additional columns from both **A** and **B** are sorted alphabetically and appended after the normal, specified variables.

The following example illustrates the described behaviour:

**A.janno**

| Poseidon_ID | Group_Name | Genetic_Sex | AdditionalColumn1 | AdditionalColumn2 |
|---|---|---|---|---|
| XXX011 | POP1 | M | A | D |
| XXX012 | POP2 | F | B | E |
| XXX013 | POP1 | M | C | F |

**B.janno**

| Poseidon_ID | Group_Name | Genetic_Sex | AdditionalColumn3 | AdditionalColumn2 |
|---|---|---|---|---|
| YYY022 | POP5 | F | G | J |
| YYY023 | POP5 | F | H | K |
| YYY024 | POP5 | M | I | L |

**A.janno + B.janno**

| Poseidon_ID | Group_Name | Genetic_Sex | AdditionalColumn1 | AdditionalColumn2 | AdditionalColumn3 |
|---|---|---|---|---|---|
| XXX011 | POP1 | M | A | D | n/a |
| XXX012 | POP2 | F | B | E | n/a |
| XXX013 | POP1 | M | C | F | n/a |
| YYY022 | POP5 | F | n/a | J | G |
| YYY023 | POP5 | F | n/a | K | H |
| YYY024 | POP5 | M | n/a | L | I |

##### 3.3.4 Treatment of the `.ssf` file while merging

The Sequencing Source File (short `.ssf` file) is forged in exactly the same way as the `.janno` file. `.ssf` files that are present are included in the forge product, following selection of those entities which are listed in the `poseidon_IDs` columns. Columns that are only present in some packages, including those not defined in the Poseidon package specification, are also included in the forged product in the same way as described for `.janno` files above.

##### 3.3.5 Treatment of the `.bib` file while merging

In the forge process all relevant samples for the output package are determined. This includes their `.janno` entries and therefore the information on the publication keys documented for them in the `.janno` `Publication` column. The output `.bib` file compiles only the relevant references for the samples in the output package. It includes the references exactly once and is sorted alphabetically by key.

##### 3.3.6 Other options

Just as for `init` the output package of `forge` is created as a new directory `-o`. The title can also be explicitly defined with `-n`.

`--minimal` allows for the creation of a minimal output package without `.bib` and `.janno`. This is especially useful for data analysis pipelines, where only the genotype data is required. Even more basic output comes with `--onlyGeno`, which means that only the genotype data is returned without any Poseidon package.

`forge` has a an optional flag `--intersect`, that defines, if the genotype data from different packages should be merged with a union or an intersect operation. See *Treatment of the genotype data while merging* above.

`--intersect` also influences the automatic determination of the `snpSet` field in the POSEIDON.yml file for the resulting package. If the `snpSets` of all input packages are identical, then the resulting package will just inherit this configuration. Otherwise `forge` applies the following pairwise merging logic:

| Input snpSet A | Input snpSet B | `--intersect` | Ouput snpSet |
|---|---|---|---|
| Other | * | * | Other |
| 1240K | HumanOrigins | True | HumanOrigins |
| 1240K | HumanOrigins | False | 1240K |

`--selectSnps` allows to provide `forge` with a SNP file in EIGENSTRAT (`.snp`) or PLINK (`.bim`) format to create a package with a specific selection. When this option is set, the output package will have exactly the SNPs listed in this file. Any SNP not listed in the file will be excluded. If `--intersect` is also set, only the SNPs overlapping between the SNP file and the forged packages are output.

With `--packagewise` the within-package selection step in forge can be skipped. This will result in outputting all individuals in the relevant packages, and hence a superset of the requested individuals/groups. It may result in better performance in cases where one wants to forge entire packages.

##### 3.4 Genoconvert command

`genoconvert` converts the genotype data in a Poseidon package to a different file format. The respective entries in the POSEIDON.yml file are changed accordingly.

Command line details

```
Usage: trident genoconvert ((-d|--baseDir DIR) |
                             ((-p|--genoOne FILE) | --inFormat FORMAT
                               --genoFile FILE --snpFile FILE --indFile FILE)
                             [--snpSet SET]) --outFormat FORMAT [--onlyGeno]
                           [-o|--outPackagePath DIR] [--removeOld]
                           [--outPlinkPopName MODE] [--onlyLatest]

  Convert the genotype data in a Poseidon package to a different file format

Available options:
  -h,--help                Show this help text
  -d,--baseDir DIR         A base directory to search for Poseidon packages.
  -p,--genoOne FILE        One of the input genotype data files. Expects .bed,
                           .bim or .fam for PLINK and .geno, .snp or .ind for
                           EIGENSTRAT. The other files must be in the same
                           directory and must have the same base name.
  --inFormat FORMAT        The format of the input genotype data: EIGENSTRAT or
                           PLINK. Only necessary for data input with --genoFile
                           + --snpFile + --indFile.
  --genoFile FILE          Path to the input geno file.
  --snpFile FILE           Path to the input snp file.
  --indFile FILE           Path to the input ind file.
  --snpSet SET             The snpSet of the package: 1240K, HumanOrigins or
                           Other. Only relevant for data input with -p|--genoOne
                           or --genoFile + --snpFile + --indFile, because the
```

```
                          packages in a -d|--baseDir already have this
                          information in their respective POSEIDON.yml files.
                          (default: Other)
  --outFormat FORMAT      the format of the output genotype data: EIGENSTRAT or
                          PLINK.
  --onlyGeno              Should only the resulting genotype data be returned?
                          This means the output will not be a Poseidon package.
  -o,--outPackagePath DIR Path to the output package directory. This is
                          optional: If no path is provided, then the output is
                          written to the directories where the input genotype
                          data file (.bed/.geno) is stored. (default: Nothing)
  --removeOld             Remove the old genotype files when creating the new
                          ones.
  --outPlinkPopName MODE  Where to write the population/group name into the FAM
                          file in Plink-format. Three options are possible:
                          asFamily (default) | asPhenotype | asBoth. See also
                          --inPlinkPopName.
  --onlyLatest            Consider only the latest versions of packages, or the
                          groups and individuals within the latest versions of
                          packages, respectively.
```

With the default setting

```
trident genoconvert -d ... -d ... --outFormat EIGENSTRAT|PLINK
```

all packages in `-d` will be converted to the desired `--outFormat` (either `EIGENSTRAT` or `PLINK`), if the data is not already in this format. This includes updating the respective POSEIDON.yml files.

The "old" data is not deleted, but kept around. That means conversion can result in a package with both PLINK and EIGENSTRAT data, but only one is linked in the POSEIDON.yml file, and that is what will be used by `trident`. To delete the old data in the conversion you can add the `--removeOld` flag.

`-p (+ --snpSet)` or `--inFormat + --genoFile + --snpFile + --indFile (+ --snpSet)` allow to directly convert genotype data that is not wrapped in a Poseidon package and store it to a directory given in `-o`. See this example:

```
trident genoconvert \
  -p 2018_Mittnik_Baltic/Mittnik_Baltic.bed \
  --outFormat EIGENSTRAT \
  -o my_directory
```

#### 3.5 Jannocoalesce command

`jannocoalesce` merges information from one or multiple source `.janno` files into a target `.janno` file.

Command line details

```
Usage: trident jannocoalesce ((-s|--sourceFile FILE) | (-d|--baseDir DIR))
                             (-t|--targetFile FILE) [-o|--outFile FILE]
                             [--includeColumns ARG | --excludeColumns ARG]
```

```
                          [-f|--force] [--sourceKey ARG] [--targetKey ARG]
                          [--stripIdRegex ARG]

  Coalesce information from one or multiple janno files to another one

Available options:
  -h,--help                Show this help text
  -s,--sourceFile FILE     The source .janno file.
  -d,--baseDir DIR         A base directory to search for Poseidon packages.
  -t,--targetFile FILE     The target .janno file to fill.
  -o,--outFile FILE        An optional file to write the results to. If not
                           specified, change the target file in place.
                           (default: Nothing)
  --includeColumns ARG     A comma-separated list of .janno column names to
                           coalesce. If not specified, all columns that can be
                           found in the source and target will get filled.
  --excludeColumns ARG     A comma-separated list of .janno column names NOT to
                           coalesce. All columns that can be found in the source
                           and target will get filled, except the ones listed
                           here.
  -f,--force               With this option, potential non-missing content in
                           target columns gets overridden with non-missing
                           content in source columns. By default, only missing
                           data gets filled-in.
  --sourceKey ARG          The .janno column to use as the source key.
                           (default: "Poseidon_ID")
  --targetKey ARG          The .janno column to use as the target key.
                           (default: "Poseidon_ID")
  --stripIdRegex ARG       An optional regular expression to identify parts of
                           the IDs to strip before matching between source and
                           target. Uses POSIX Extended regular expressions.
```

A most basic run may just include two arguments:

```
trident jannocoalesce \
  --sourceFile path/to/source.janno \
  --targetFile path/to/target.janno
```

`jannocoalesce` generally works by reading a source `.janno` file with `-s|--sourceFile` (or all `.janno` files in a `-d|--baseDir`) and a target `.janno` file with `-t|--targetFile`.

It then merges these files by a key column, which can be selected with `--sourceKey` and `--targetKey`. The default for both of these key columns is the `Poseidon_ID`. In case the entries in the key columns slightly and systematically differ, e.g. because the `Poseidon_ID`s in either have a special suffix (for example `_SG`), then the `--stripIdRegex` option allows to strip these with a regular expression to thus match the keys.

`jannocoalesce` generally attempts to fill **all** empty cells in the target `.janno` file with information from the source. `--includeColumns` and `--excludeColumns` allow to select specific columns for which this should be done. In some cases it may be desirable to not just fill empty fields in the target, but overwrite the information already there with the `-f|--force` option. If the target file should be preserved, then the output can be directed to a new output `.janno` file with `-o|--outFile`.

#### 3.6 Rectify command

`rectify` automatically harmonizes POSEIDON.yml files of one or multiple packages. This is not an automatic update from one Poseidon version to the next, but rather a clean-up wizard after manual modifications.

Command line details

```
Usage: trident rectify (-d|--baseDir DIR) [--ignorePoseidonVersion]
                       [--poseidonVersion ?.?.?]
                       [--packageVersion VPART [--logText STRING]]
                       [--checksumAll | [--checksumGeno] [--checksumJanno]
                         [--checksumSSF] [--checksumBib]]
                       [--newContributors DSL] [--onlyLatest]

  Adjust POSEIDON.yml files automatically to package changes

Available options:
  -h,--help                Show this help text
  -d,--baseDir DIR         A base directory to search for Poseidon packages.
  --ignorePoseidonVersion  Read packages even if their poseidonVersion is not
                           compatible with trident.
  --poseidonVersion ?.?.?  Poseidon version the packages should be updated to:
                           e.g. "2.5.3".
  --packageVersion VPART   Part of the package version number in the
                           POSEIDON.yml file that should be updated: Major,
                           Minor or Patch (see https://semver.org).
  --logText STRING         Log text for this version in the CHANGELOG file.
  --checksumAll            Update all checksums.
  --checksumGeno           Update genotype data checksums.
  --checksumJanno          Update .janno file checksum.
  --checksumSSF            Update .ssf file checksum
  --checksumBib            Update .bib file checksum.
  --newContributors DSL    Contributors to add to the POSEIDON.yml file in the
                           form "[Firstname Lastname](Email address);...".
  --onlyLatest             Consider only the latest versions of packages, or the
                           groups and individuals within the latest versions of
                           packages, respectively.
```

It can be called with a lot of optional arguments. Note that `rectify` by default does **not** apply any changes if none of these arguments are set. Each change requires explicit opt-in.

```
trident rectify -d ... -d ... \
  --poseidonVersion "X.X.X" \
```

```
--packageVersion Major|Minor|Patch \
--logText "short description of the update" \
--checksumAll \
--newContributors "[Firstname Lastname](Email address);..."
```

The following arguments determine which fields of the POSEIDON.yml file should be modified:

- `--poseidonVersion` allows a simple change of the `poseidonVersion` field in the POSEIDON.yml file.
- `--packageVersion` increments the package version number in the first, the second or the third position. It can optionally be called with `--logText`, which appends an entry to the CHANGELOG file for the respecitve package version update. `--logText` also creates a new CHANGELOG.md file if it does not exist yet.
- `--checksumGeno`, `--checksumJanno`, `--checksumSSF` and `--checksumBib` add or modify the respective checksum fields in the POSEIDON.yml file. `--checksumAll` is a wrapper to call all of them at once.
- `--newContributors` adds new contributors.

As `rectify` reads and rewrites POSEIDON.yml files, it may change their inner order, layout or even content (e.g. if they have fields which are not in the POSEIDON.yml specification). Create a backup of the POSEIDON.yml file before running `rectify` if you are uncertain if this might affect you negatively.

### 4 Inspection commands

#### 4.1 List command

`list` lists packages, groups and individuals of local datasets, or of packages available in the archives on the web server.

Command line details

```
Usage: trident list ((-d|--baseDir DIR) | --remote [--remoteURL URL]
                      [--archive STRING])
                    (--packages | --groups | --individuals
                      [-j|--jannoColumn COLNAME]) [--raw] [--onlyLatest]

  List packages, groups or individuals from local or remote Poseidon
  repositories

Available options:
  -h,--help                Show this help text
  -d,--baseDir DIR         A base directory to search for Poseidon packages.
  --remote                 List packages from a remote server instead the local
                           file system.
  --remoteURL URL          URL of the remote Poseidon server.
                           (default: "https://server.poseidon-adna.org")
  --archive STRING         The name of the Poseidon package archive that should
                           be queried. If not given, then the query falls back
                           to the default archive of the server selected with
                           --remoteURL. See the archive documentation at
```

```
                            https://www.poseidon-adna.org/#/archive_overview for
                            a list of archives currently available from the
                            official Poseidon Web API. (default: Nothing)
  --packages                List all packages.
  --groups                  List all groups, ignoring any group names after the
                            first as specified in the .janno-file.
  --individuals             List all individuals/samples.
  -j,--jannoColumn COLNAME List additional fields from the janno files, using
                            the .janno column heading name, such as "Country",
                            "Site", "Date_C14_Uncal_BP", etc..
  --raw                     Return the output table as tab-separated values
                            without header. This is useful for piping into grep
                            or awk.
  --onlyLatest              Consider only the latest versions of packages, or the
                            groups and individuals within the latest versions of
                            packages, respectively.
```

To list packages from your local repositories, as seen above you can run

```
trident list -d ... -d ... --packages
```

This will yield a nicely formatted table of all packages, their version and the number of individuals in them.

You can use `--remote` to show packages on the remote server. For example

```
trident list --packages --remote --archive "community-archive"
```

will result in a view of all packages available in one of the public Poseidon archives. Just as for `fetch`, the `--archive` flag allows to choose which public archive to query.

Independent of whether you query a local or an online archive, you can not just list packages, but also groups, as defined in the third column of EIGENSTRAT `.ind` files (or the first/last column of a PLINK `.fam` file), and individuals with the flags `--groups` and `--individuals` (instead of `--packages`).

The `--individuals` flag additionally provides a way to immediately access information from `.janno` files on the command line. This works with the `-j|--jannoColumn` option. For example adding `-j Country -j Date_C14_Uncal_BP` to the commands above will add the `Country` and the `Date_C14_Uncal_BP` columns to the respective output tables.

Note that if you want a less ornate table, for example because you want to load this into Excel, or pipe into another command that cannot deal with the table layout, you can use the `--raw` option to output that table as a simple tab-delimited stream.

#### 4.2 Summarise command

`summarise` prints some general summary statistics for a given Poseidon dataset taken from the `.janno` files.

Command line details

```
Usage: trident summarise (-d|--baseDir DIR) [--raw]
```

```
  Get an overview over the content of one or multiple Poseidon packages

Available options:
  -h,--help                Show this help text
  -d,--baseDir DIR         A base directory to search for Poseidon packages.
  --raw                    Return the output table as tab-separated values
                           without header. This is useful for piping into grep
                           or awk.
```

You can run it with

```
trident summarise -d ... -d ...
```

which will show you context information like – among others – the number of individuals in the dataset, their sex distribution, the mean age of the samples or the mean coverage on the 1240K SNP array in a table. `summarise` depends on complete `.janno` files and will silently ignore missing information.

You can use the `--raw` option to output the summary table in a simple, tab-delimited layout.

#### 4.3 Survey command

`survey` tries to indicate package completeness (mostly focused on `.janno` files) for Poseidon datasets.

Command line details

```
Usage: trident survey (-d|--baseDir DIR) [--raw] [--onlyLatest]

  Survey the degree of context information completeness for Poseidon packages

Available options:
  -h,--help                Show this help text
  -d,--baseDir DIR         A base directory to search for Poseidon packages.
  --raw                    Return the output table as tab-separated values
                           without header. This is useful for piping into grep
                           or awk.
  --onlyLatest             Consider only the latest versions of packages, or the
                           groups and individuals within the latest versions of
                           packages, respectively.
```

Running

```
trident survey -d ... -d ...
```

will yield a table with one row for each package. See `trident survey -h` for a legend which cell of this table means what.

Again you can use the `--raw` option to output the survey table in a tab-delimited format.

#### 4.4 Validate command

`validate` checks Poseidon packages and individual package components for structural correctness.

Command line details

```
Usage: trident validate ((-d|--baseDir DIR) [--ignoreGeno] [--fullGeno]
                          [--ignoreDuplicates] [-c|--ignoreChecksums]
                          [--ignorePoseidonVersion] |
                          --pyml FILE | (-p|--genoOne FILE) | --inFormat FORMAT
                          --genoFile FILE --snpFile FILE --indFile FILE |
                          --janno FILE | --ssf FILE | --bib FILE) [--noExitCode]
                        [--onlyLatest]

  Check Poseidon packages or package components for structural correctness

Available options:
  -h,--help                Show this help text
  -d,--baseDir DIR         A base directory to search for Poseidon packages.
  --ignoreGeno             Ignore snp and geno file.
  --fullGeno               Test parsing of all SNPs (by default only the first
                           100 SNPs are probed).
  --ignoreDuplicates       Do not stop on duplicated individual names in the
                           package collection.
  -c,--ignoreChecksums     Whether to ignore checksums. Useful for speedup in
                           debugging.
  --ignorePoseidonVersion  Read packages even if their poseidonVersion is not
                           compatible with trident.
  --pyml FILE              Path to a POSEIDON.yml file.
  -p,--genoOne FILE        One of the input genotype data files. Expects .bed,
                           .bim or .fam for PLINK and .geno, .snp or .ind for
                           EIGENSTRAT. The other files must be in the same
                           directory and must have the same base name.
  --inFormat FORMAT        The format of the input genotype data: EIGENSTRAT or
                           PLINK. Only necessary for data input with --genoFile
                           + --snpFile + --indFile.
  --genoFile FILE          Path to the input geno file.
  --snpFile FILE           Path to the input snp file.
  --indFile FILE           Path to the input ind file.
  --janno FILE             Path to a .janno file.
  --ssf FILE               Path to a .ssf file.
  --bib FILE               Path to a .bib file.
  --noExitCode             Do not produce an explicit exit code.
  --onlyLatest             Consider only the latest versions of packages, or the
                           groups and individuals within the latest versions of
                           packages, respectively.
```

You can run it with

```
trident validate -d ... -d ...
```

to check packages and it will either report a success (`Validation passed`) or failure with specific error messages.

Instead of validating entire packages with `-d` you can also apply it to individual files and package components: `--pyml` (POSEIDON.yml), `-p | --inFormat + --genoFile + --snpFile + --indFile` (genotype data), `--janno` (.janno file), `--ssf` (.ssf file) or `--bib` (.bib file). In this case `validate` attempts to read and parse the respecitve files individually and reports any issues it encounters. Note that this considers the files in isolation and does not include any cross-file consistency checks.

When applied to packages, `validate` tries to ensure that each package adheres to the Poseidon package specification. Here is a list of what is checked:

- Structural correctness of the POSEIDON.yml file.
- Presence of all files references in the POSEIDON.yml file.
- Full structural correctness of .janno, `.ssf` and `.bib` file.
- Superficial correctness of genotype data files by parsing the first 100 SNPs. A full check that parses all SNPs can be triggered with the `--fullGeno` option. `--ignoreGeno`, on the other hand, causes `validate` to ignore the genotype data entirely, which speeds up the validation significantly.
- Correspondence of BibTeX keys in `.bib` and .janno
- Correspondence of sample IDs in `.janno` and .ssf.
- Correspondence of sample and group IDs in `.janno` and genotype data files.

In fact much of this validation already runs as part of the general package reading pipeline invoked for other `trident` subcommands (e.g. `forge`). `validate` is meant to be more thorough and brittle, though, and will explicitly fail if even a single package is broken. For special cases more flexibility can be enabled with the options `--ignoreDuplicates`, `--ignoreChecksums` and `--ignorePoseidonVersion`.

Remember to run `validate` with `--debug` to get more information in case the default output is not sufficient to analyse an issue.
