## Supplementary Text 4 - Xerxes for "Poseidon – A framework for archaeogenetic human genotype data management"

### Guide for xerxes v1.0.1.0

**3 RAS command (in development)** **7**

### 1 Installation

See the Poseidon website (https://www.poseidon-adna.org/#/xerxes) or the GitHub repository (https://github.com/poseidon-framework/poseidon-analysis-hs) for up-to-date installation instructions.

### 2 Fstats command

Xerxes allows you to analyse genotype data across Poseidon packages, including your own, by pre-loading sets of packages via a `--baseDir` (or `-d`) parameter. From the pre-loaded packages it selects the ones relevant for the statistics you requested, and then streams through only these. It thus computes arbitrary F-Statistics across groups and individuals distributed in many packages, without the need to explicitly merge data first.

In this process `xerxes` takes care of merging PLINK and EIGENSTRAT data on the fly and works across different SNP sets, like Human-Origins and 1240k. It flips alleles automatically across genotype files, and throws an error if the alleles in different packages are incongruent with each other.

Here is an example command for computing several F-Statistics:

```
xerxes fstats -d ... -d ... \
  --stat "F4(<Chimp.REF>, <Altai_published.DG>, Yoruba, French)" \
  --stat "F3(<Chimp.REF>, <Altai_snpAD.DG>, Spanish)" \
  --statFile fstats.txt
  --statConfig fstats.yaml
  -f outputfile.txt
```

First, the two options `-d ...` exemplify that you need to provide at least one base directory with source Poseidon packages, but can also give multiple. Second, F-Statistics can be entered in three different ways:

1. Directly via the command line using `--stat`.
2. With a simple text file using `--statFile`.
3. With a powerful configuration file format via `--statConfig`.

These three modes of input can be mixed and matched, and even given multiple times. They are explained below.

Last, the option `-f` can be used to write the output table into a tab-separated text file, beyond just printing a table into the standard out when the program finishes. Note that there are more options, which you can view using `xerxes fstats --help`:

```
Usage: xerxes fstats (-d|--baseDir DIR) [-j|--jackknife ARG]
                     [-e|--excludeChroms ARG]
                     (--stat ARG | --statConfig ARG | --statFile ARG)
                     [--noTransitions] [-f|--tableOutFile ARG]
                     [--blockTableFile ARG]

  Compute f-statistics on groups and invidiuals within and across Poseidon
  packages

Available options:
  -h,--help                Show this help text
  -d,--baseDir DIR         A base directory to search for Poseidon packages.
  -j,--jackknife ARG       Jackknife setting. If given an integer number, this
                           defines the block size in SNPs. Set to "CHR" if you
                           want jackknife blocks defined as entire chromosomes.
                           The default is at 5000 SNPs
  -e,--excludeChroms ARG   List of chromosome names to exclude chromosomes,
                           given as comma-separated list. Defaults to X, Y, MT,
                           chrX, chrY, chrMT, 23,24,90
  --stat ARG               Specify a summary statistic to be computed. Can be
                           given multiple times. Possible options are: F4(a, b,
                           c, d), F3(a, b, c), F3star(a, b, c), F2(a, b), PWM(a,
                           b), FST(a, b), Het(a) and some more special options
                           described at
                           https://poseidon-framework.github.io/#/xerxes?id=fstats-command.
                           Valid entities used in the statistics are group names
                           as specified in the *.fam, *.ind or *.janno failes,
                           individual names using the syntax "<Ind_name>", so
```

```
                             enclosing them in angular brackets, and entire
                             packages like "*Package1*" using the Poseidon package
                             title. You can mix entity types, like in
                             "F4(<Ind1>,Group2,*Pac*,<Ind4>)". Group or individual
                             names are separated by commas, and a comma can be
                             followed by any number of spaces.
  --statConfig ARG           Specify a yaml file for the Fstatistics and group
                             configurations
  --statFile ARG             Specify a file with F-Statistics specified similarly
                             as specified for option --stat. One line per
                             statistics, and no new-line at the end
  --maxSnps ARG              Stop after a maximum nr of snps has been processed.
                             Useful for short test runs
  --noTransitions            Skip transition SNPs and use only transversions
  -f,--tableOutFile ARG      a file to which results are written as tab-separated
                             file
  --blockTableFile ARG       a file to which the per-Block results are written as
                             tab-separated file
```

#### 2.1 Allowed statistics

The following statistics are allowed in the --stat, --statFile and --statConfig options. In all of the following, the symbols a, b, c or d stand for arbitrary entities allowed in Poseidon, so groups (such as French), individuals (such as <MA1.SG>) or packages (such as *2012_PattersonGenetics*).

- F2vanilla(a,b): F2-Statistics - Vanilla version. Computed using F2vanilla(a,b) = (a-b)^2 across the genome.
- F2(a,b): F2-Statistics (bias-corrected version). Computed as F2(a, b) = F2vanilla(a,b) - hA/sA - hB/sB, where sA is the number of non-missing alleles in entity a, and hA = nA * nA' / sA * (sA - 1) is an estimator of half the heterozygosity (see Het(a)), and likewise for sB and nB etc.
- F3vanilla(a,b,c): F3-Statistics - Vanilla version, recommended if used as Outgroup-F3 statistics or with group c being pseudo-haploid: Are computed as F3(a,b,c) = (c-a)(c-b) across all SNPs.
- F3(a,b,c): F3-Statistics (bias-corrected version). Computed as F3(a,b,c) = F3vanilla(a,b) - hC/sC.
- F3star(a,b,c): F3-Statistics as defined in [1] - normalised and bias-corrected version, recommended for Admixture-F3 tests. Are computed by i) first subtracting per SNP from the vanilla-F3-Statistic a bias-correction term hC/sC, as above for F2, and ii) then normalising the genome-wide estimate by a genome-wide estimate of the heterozygosity of entity c (Het(c)), in order to make results comparable between different groups c.
- F4(a,b,c,d): F4 statistics. Are computed by averaging the quantity (a-b)(c-d) across all SNPs. No bias correction is necessary for this statistic.
- Het(a): An estimate of the heterozygosity across all SNPs, computed as 2*hA, with hA defined as above for F2.
- FST(a,b): An estimate of FST across the genome, following the estimator presented in [2] and implemented in the ADMIXTOOLS package. This amounts to a ratio of genome-wide ranges, where the numerator is an unbiased estimate of F2 (see above), and the denominator is PWM(a,b) (see below).
- FSTvanilla(a,b): Similar to FST(a,b) but without the bias correction in the numerator, mainly useful

for teaching and learning.
- `PWM(a,b)`: The pairwise mismatch rate between entities `a` and `b`, computed from allele frequencies as `a (1 - b) + (1 - a) b`.

Most of these equations can also be found in [1]. See also Appendix A of this paper for the unbiased estimators used above.

For each of the “slots” A, B, C or D, you can enter:

- Individuals, using the syntax `<Individual_Name>`
- Groups, using no special syntax `Group_Name`
- Packages, using syntax `*Package_Name*` (This can be useful, for example, if you happen to have a homogenous set of individuals from multiple groups in one package and want to consider all of these as one group.)

#### 2.2 Defining statistics directly via `--stat`

This is the simplest option to instruct the program to compute a specified statistic. Each statistic requires a separate input using `--stat` using this input method. Example:

```
xerxes fstats -d ... -d ... --stat "F3(French, Spanish, <Chimp.REF>) --stat "FST(French, Spanish)"
```

#### 2.3 Defining statistics in a simple text file

You can prepare a text file, e.g. `fstats.txt`, into which you write the above statistics, one statistic per line. Example:

```
F4(<Chimp.REF>, <Altai_published.DG>, Yoruba, French)
F4(<Chimp.REF>, <Altai_snpAD.DG>, Spanish, French)
F4(Mbuti,Nganasan,Saami.DG,Finnish)
```

You can then load these statistics using the option `--statFile fstats.txt`.

#### 2.4 Input via a configuration file

This is the most powerful way to input F-Statistics. Example:

```yaml
groupDefs:
  CEU2: ["CEU.SG", "-<NA12889.SG>", "-<NA12890.SG>"]
  FIN2: ["FIN.SG", "-<HG00383.SG>", "-<HG00384.SG>"]
  GBR2: ["GBR.SG", "-<HG01791.SG>", "-<HG02215.SG>"]
  IBS2: ["IBS.SG", "-<HG02238.SG>", "-<HG02239.SG>"]
fstats:
- type: F2
  a: ["French", "Spanish"]
  b: ["Han", "CEU2"]
  # Ascertainment is optional
- type: F3 # This will create 3x2x1 = 6 Statistics
  a: ["French", "Spanish", "Mbuti"]
  b: ["Han", "CEU2"]
```

```
  c: ["<Chimp.REF>"]
  ascertainment:
    outgroup: "<Chimp.REF>" # ascertaining on outgroup-polarised derived allele frequency
    reference: "CEU2"
    lower: 0.05
    upper: 0.95
- type: F4 # This will create 5x5x4x1 = 100 Statistics
  a: ["<I0156.SG>", "<I0157.SG>", "<I0159.SG>", "<I0160.SG>", "<I0161.SG>"]
  b: ["<I0156.SG>", "<I0157.SG>", "<I0159.SG>", "<I0160.SG>", "<I0161.SG>"]
  c: ["CEU2", "FIN2", "GBR2", "IBS2"]
  d: ["<Chimp.REF>"]
  ascertainment:
    # A missing outgroup means: ascertain on minor allele frequency
    reference: "CEU.SG"
    lower: 0.00
    upper: 0.10
```

You can save this into a text file, for example named `fstats_config.yaml`, and load it via `--statConfig fstats_config.yaml`.

The top level structure of this YAML file is an object with two fields: `groupDefs` (which is optional) and `fstats` (which is mandatory).

##### 2.4.1 Group Definitions

You can specify ad-hoc group definitions using the syntax above. Every group consists of a name (used as object key) and then a JSON- or YAML-list of signed entities, following the same syntax as `trident forge`. Briefly: Individuals, Groups and Packages can be added or excluded (prefixed by a `-`) in order. In the example above, two individuals are removed from each group.

Note that currently, groups can be defined only independently, so not incremental to each other. That means, you cannot currently use an already defined new group name in the entity list of a following group name.

##### 2.4.2 Statistic input using YAML

Each statistic defined in the `fstats` section of the YAML file, actually defines a loop over multiple populations in each statistic. In the example above, there are 6 F3-Statistics, each using a different combination of the input groups defined in each of the `a:`, `b:` and `c:` slots. There are also 100 (!) F4 statistics, following all combinations of 5x5x4x1 slots defined in `a:`, `b:`, `c:` and `d:`.

##### 2.4.3 Ascertainment (experimental feature)

In addition, every statistic section allows for a definition of an ascertainment specification, using a special key `ascertainment:`, which is optional. If given, you can specify an optional `outgroup`, a `reference` group in which to ascertain SNPs, and `lower` and `upper` allele frequency bounds. If specified, only SNPs for which the `reference` group has an allele frequency within the given bounds are used to compute the statistic (note that normalisation is still using all non-missing SNPs for that given statistic). If an `outgroup` is defined, then the outgroup-polarised derived allele frequency is used. If no `outgroup` is defined, then the minor allele frequency is used instead. If an outgroup is defined, any sites where the outgroup is polymorphic are treated as missing.

#### 2.5 Output

The final output of the `fstats` command looks like this:

```
.-----------.-----------.---------------.--------.-------.---------.
| Statistic |     a     |       b       |   c    |   d   | NrSites |
:===========:===========:===============:========:=======:=========:
| F3        | French    | Italian_North | Mbuti  |       | 593124  |
| F3        | French    | Han           | Mbuti  |       | 593124  |
| F3        | Sardinian | Pima          | French |       | 593124  |
| F4        | French    | Russian       | Han    | Mbuti | 593124  |
| F4        | Sardinian | French        | Pima   | Mbuti | 593124  |
'-----------'-----------'---------------'--------'-------'---------' ->

----------------.--------------------.------------------.---------------------.
 Estimate_Total | Estimate_Jackknife | StdErr_Jackknife |  Z_score_Jackknife  |
================:====================:==================:=====================:
 5.9698e-2      | 5.9698e-2          | 5.1423e-4        | 116.0908951980249   |
 5.0233e-2      | 5.0233e-2          | 5.0324e-4        | 99.81843057232513   |
 -1.2483e-3     | -1.2483e-3         | 9.2510e-5        | -13.493505348221081 |
 -1.6778e-3     | -1.6778e-3         | 9.1419e-5        | -18.35262346091248  |
 -1.4384e-3     | -1.4384e-3         | 1.1525e-4        | -12.481084899924868 |
----------------'--------------------'------------------'---------------------'
```

This output table lists each statistic, the slots `a`, `b`, `c` and `d`, the number of sites with non-missing data for that statistic, ascertainment information (outgroup, reference, lower and upper bound, if given), the genome-wide estimate, its standard error and its Z-score. If you specify an output file using option `--tableOutFile` or `-f`, these results are also written as a simple, tab-separated file.

Additionally, an option `--blockOutFile` can be specified. This creates a file to which a table with estimates per Jackknife block is written.

#### 2.6 Degenerate statistics

Specific cases of statistics yield zero by construction:

- `F2(a,b)`, `F2vanilla(a,b)`, `FST(a,b)` and `FSTvanilla(a,b)` where `a=b`.
- `F3(a,b,c)` and `F3vanilla(a,b,c)` where `c=a` or `c=b`
- `F4(a,b,c,d)` where `a=b` or `c=d`

Even though the bias-correction technically can result in non-zero and even negative values, we automatically detect these cases and output zero for them. This can be useful, for example, when looping over pairs of populations for a pairwise matrix of `FST`, where we then want the diagonal to be zero to yield a proper distance matrix.

#### 2.7 Ploidy and illegal cases

Genotype ploidy in input samples is important for many of the statistics, because the bias-correction terms require the number of chromosomes. Ploidy information is automatically read through the field of `Genotype_Ploidy` in the `.janno` file. A warning is printed if that information is missing, in which case we assume diploid genotypes.

But often with low-coverage data from ancient DNA we create pseudo-haploid genotypes, so in that case it is important to provide that information correctly through the `.janno` file.

In specific cases statistics are illegal with only a single haplotype. Specifically:

- `F2(a,b)` and `FST(a,b)` is undefined if either one of `a` or `b` contains only a single haplotype.
- `F3(a,b,c)` is undefined if `c` contains only a single haplotype.
- `Het(a)` unsurprisingly is undefined if `a` contains only a single haplotype.

These cases are detected and an error is thrown. For `F2`, `F3` and `FST` you can use the "vanilla" versions of the statistics if that makes sense. This is particularly relevant for so-called "Outgroup-F3-Statistics", where we sometimes use a single haploid reference genome in position `c`. Use `F3vanilla` in that case.

##### 2.8 Whitepaper

The repository comes with a detailed whitepaper that describes some more mathematical details of the methods implemented here.

#### 3 RAS command (in development)

The RAS command computes pairwise RAS statistics between a collection of "left" entities, and a collection of "right" entities. Every entity is either a group name or an individual, with similar syntax as for the F-Statistics above, so `French` is a group, and `<IND001>` is an individual.

The input of left-pops and right-pops uses a YAML file via `--popConfigFile`. Here is an example:

```
groupDefs:
  group1: a,b,-c,-<d>
  group2: e,f,-<g>
popLefts:
- <I13721>
- <I14000>
- <I13722>
- <Iceman.SG>
popRights:
- Mbuti
- Mixe
- Spanish
outgroup: <Chimp.REF>
```

In this case, two groups are defined on the fly: `group1` comprises groups `a` and `b`, but excludes group `c` and individual `d`. Note that inclusions and exclusions are executed in order. `group2` comprises of group `e` and group `f`, but excludes individual `<g>`.

As in RAScalculator [3], the allele frequency ascertainment is done across right populations only.

The are a couple of options, as specified in the CLI help (`xerxes ras --help`):

```
Usage: xerxes ras (-d|--baseDir DIR) [-j|--jackknife ARG]
                  [-e|--excludeChroms ARG] --popConfigFile ARG
                  (--minAC ARG | --minFreq ARG | --noMinFreq)
```

```
                 (--maxAC ARG | --maxFreq ARG | --noMaxFreq)
                 [-m|--maxMissingness ARG] [--blockTableFile ARG]
                 [--f4TableOutFile ARG] [--noTransitions] [--bedFile ARG]

  Compute RAS statistics on groups and individuals within and across Poseidon
  packages

Available options:
  -h,--help                Show this help text
  -d,--baseDir DIR         A base directory to search for Poseidon packages.
  -j,--jackknife ARG       Jackknife setting. If given an integer number, this
                           defines the block size in SNPs. Set to "CHR" if you
                           want jackknife blocks defined as entire chromosomes.
                           The default is at 5000 SNPs
  -e,--excludeChroms ARG   List of chromosome names to exclude chromosomes,
                           given as comma-separated list. Defaults to X, Y, MT,
                           chrX, chrY, chrMT, 23,24,90
  --popConfigFile ARG      a file containing the population configuration
  --minAC ARG              define a minimal allele-count cutoff for the RAS
                           statistics.
  --minFreq ARG            define a minimal allele-frequency cutoff for the RAS
                           statistics.
  --noMinFreq              switch off the minimum allele frequency filter
  --maxAC ARG              define a maximal allele-count cutoff for the RAS
                           statistics.
  --maxFreq ARG            define a maximal allele-frequency cutoff for the RAS
                           statistics.
  --noMaxFreq              switch off the maximum allele frequency filter. This
                           cam help mimic Outgroup-F3
  -m,--maxMissingness ARG  define a maximal missingness for the right
                           populations in the RAS statistics. (default: 0.1)
  --blockTableFile ARG     a file to which the per-Block results are written as
                           tab-separated file
  --f4TableOutFile ARG     a file to which F4 computations are written as
                           tab-separated file
  --maxSnps ARG            Stop after a maximum nr of snps has been processed.
                           Useful for short test runs
  --noTransitions          Skip transition SNPs and use only transversions
  --bedFile ARG            An optional bed file that gives sites to be included
                           in the analysis.
```

The output gives both cumulative (up to allele-count k) and per-allele-frequency RAS (for allele count k) for every pair of left and rights. The standard output contains a pretty-printed table. A tab-separated file is written to the file specified using the option `-f`.

`xerxes ras` makes a few important assumptions:

1. It assumes that the Right Populations are "nearly" completely non-missing. Any allele that is actually missing from the rights is in fact treated as homozygous-reference! A different approach would be to compute the actual frequencies on the non-missing right alleles, but then we cannot any more nicely accumulate over different ascertainment allele counts.
2. If no outgroup is specified, the ascertainment operates on minor-allele frequency (as in `fstats`)
3. If an outgroup is specified and missing from a SNP, or if the SNP is polymorphic, the SNP is skipped as missing

---

[1] N. Patterson *et al.*, "Ancient admixture in human history," *Genetics*, vol. 192, no. 3, pp. 1065–1093, Nov. 2012, doi: 10.1534/genetics.112.145037.

[2] G. Bhatia, N. Patterson, S. Sankararaman, and A. L. Price, "Estimating and interpreting FST: The impact of rare variants," *Genome Research*, vol. 23, no. 9, pp. 1514–1521, Jul. 2013, doi: 10.1101/gr.154831.113.

[3] P. Flegontov *et al.*, "Palaeo-Eskimo genetic ancestry and the peopling of Chukotka and North America," *Nature*, vol. 570, no. 7760, pp. 236–240, Jun. 2019, doi: 10.1038/s41586-019-1251-y.
