## Supplementary Text 5 - Xerxes Whitepaper for "Poseidon – A framework for archaeogenetic human genotype data management"

### Xerxes Theoretical Background - Derivations and Statistical Details of the fstats command

Stephan Schiffels

Last update: March 2024

**5 Frequency-Ascertainment** **8**

The program `xerxes fstats` within the poseidon-analysis-hs package serves as a one-stop shop for various SNP-based statistics that can be computed from genotype data, including F-Statistics, pairwise mismatch rate, $F_{\mathrm{ST}}$ and likely more statistics in the future. F-Statistics were defined in a series of papers by Nick Patterson, David Reich and others, and comprehensively formally described in [1]. Interested readers may also consult [2] as a useful reference.

### 1 F-Statistics

The most basic F-Statistics available in `fstats` are:

$$F2_{\text{vanilla}}(A,B) = \langle (a-b)^2 \rangle \tag{1}$$

$$F3_{\text{vanilla}}(A,B;C) = \langle (c-a)(c-b) \rangle \tag{2}$$

$$F4(A,B;C,D) = \langle (a-b)(c-d) \rangle \tag{3}$$

Here, capital letters A, B, C, D stand for individuals or groups of individuals, defined via a Mini-Language used in trident forge, or as group definitions in FStats-Configuration files described here: `https://www.poseidon-adna.org/#/xerxes?id=input-via-a-configuraton-file`.

Small letters a, b, c, d here stand for the observed allele frequencies (i.e. the fractions of observed non-missing alternative alleles over observed non-missing alleles within groups A, B, C, D). The average $\langle \cdot \rangle$ is an average over all available SNPs. For example, we have

$$\langle (a-b)^2 \rangle = \frac{\sum_{i=1}^{L} (a_i - b_i)^2 I(a_i, b_i)}{\sum_{i=1}^{L} i(a_i, b_i)} \tag{4}$$

where

$$I(a_i, b_i) = \begin{cases} 1 & \text{if both } a_i \text{ and } b_i \text{ are non-missing} \\ 0 & \text{else} \end{cases} \tag{5}$$

Here, $a_i$ and $b_i$ denote allele frequencies of entity A and B at SNP i. Non-missing here means that at least one allele in each groups is non-missing, such that a sample allele frequency is defined. Parts of group A and B could be missing, though.

#### 1.1 Bias-correction

Many of the population genetic use-cases and properties of F-Statistics (described in [1]) are derived from *population* allele frequencies, that is, precise frequencies of an entire population. In practice, however, we only have a finite number of samples (in the most extreme case, one individual) to estimate population frequencies.

More precisely, given *population* allele frequencies $a'$ and $b'$, we consider *sample* allele frequencies $a$ and $b$ as imperfect *estimators* of the population allele frequencies.

To reason about biases of these estimators, we first denote a noise-model. Under the assumption that the $n$ samples of population A are independent samples, the sampling probability for observing $k$ non-reference alleles within a sample of $n$ samples, given a population frequency $a'$ is a binomial distribution:

$$p_n(k|a') = \text{Binom}(k,n|a') = \binom{n}{k} (a')^k (1-a')^{n-k} \tag{6}$$

We can then define the estimator for the population allele frequency $a'$:

$$a = \frac{k}{n} \tag{7}$$

and can show, for example using Wolfram Mathematica's computer algebra system, that this estimator is indeed unbiased:

```
In[1]:= Expectation[k/n,
    Distributed[k, BinomialDistribution[n, aPrime]]]

Out[1]= aPrime
```

i.e. that its expectation is just the population allele frequency $a$' itself.

This fact can be used to show that F4-statistics using sample allele frequencies are unbiased estimators of the same statistics using population frequencies:

$$
\begin{aligned}
E\left(\mathrm{F4}(a,b,c,d)\right) &= E((a-b)(c-d)) \\
&= E(ac) - E(ad) - E(bc) + E(bd) \\
&= E(a)E(c) - E(a)E(d) - E(b)E(c) + E(b)E(d) \\
&= a'c' - a'd' - b'c' + b'd' \\
&= \mathrm{F4}(a', b', c'd')
\end{aligned}
\tag{8}
$$

In the second step we have made use of the fact that because the terms in the average-brackets are based on separate samples of individuals, the average over their product factorises. So for example we have

$$E(ac) = E(a)E(c) = a'c'. \tag{9}$$

Now, the same is *not* true for quadratic terms. Concretely, consider the vanilla-F2 statistic:

$$
\begin{aligned}
F_{2,\text{vanilla}}(a,b) &= E\left((a-b)^2\right) \\
&= E\left(a^2 - 2ab + b^2\right) \\
&= E\left(a^2\right) - 2E\left(ab\right) + E(b^2)
\end{aligned}
\tag{10}
$$

Here, the middle term is again unproblematic, see equation 9. But the quadratic terms are not unbiased. For example, we have

```
In[2]:=  Expectation[(k/n)^2,
     Distributed[k, BinomialDistribution[n, aPrime]]]

                          2              2
             aPrime - aPrime   + aPrime    n
Out[2]= ----------------------------------
                         n
```

or

$$= (a')^2 + \frac{a'(1-a')}{n} \tag{11}$$

In other words $\left\langle a^2 \right\rangle$ *overestimates* $(a')^2$ by a term inversely proportional to $1/n$, so the smaller the sample size the larger the bias.

Fortunately, we can *correct* this bias, by using a different estimator for $(a')^2$. Specifically, we can try to simply subtract the term itself from the estimator, using $a$ instead of $a'$:

```
In[3]:=  Expectation[k^2/n^2 - (k/n (1 - k/n))/n,
 k Distributed[BinomialDistribution[n, a]]
                 2      2  2
             a - a   + a  n
Out[3]= ---------------
                  2
```

```
                   n
```

which turns out not to be quite right yet. It turns out, replacing $n$ by $n-1$ in the denominator does the job:

```

In[4]:= Expectation[k^2/n^2 - (k/n (1 - k/n))/(n - 1),
 k Distributed[BinomialDistribution[n, a]]]

            2
Out[4]= a
```

So we now have an unbiased estimator for the square of the allele frequency

$$E\left(a^2 - \frac{a(1-a)}{n-1}\right) = (a')^2 \tag{12}$$

which is what we need to describe unbiased estimators for F2 and F3 statistics. The following are the estimators proposed in [1] (in Appendix A, page 1089):

$$F_2(a,b) = \left\langle (a-b)^2 - \frac{a(1-a)}{n_a - 1} - \frac{b(1-b)}{n_b - 1} \right\rangle \tag{13}$$

$$F_3(a,b,c) = \left\langle ((c-a)(c-b) - \frac{c(1-c)}{n_c - 1} \right\rangle \tag{14}$$

We can easily show that they are indeed unbiased:

$$\begin{aligned} F_2(a,b) &= E\left((a-b)^2 - \frac{a(1-a)}{n_a - 1} - \frac{b(1-b)}{n_b - 1}\right) \\ &= E\left(a^2 - 2ab + b^2 - \frac{a(1-a)}{n_a - 1} - \frac{b(1-b)}{n_b - 1}\right) \\ &= E\left(a^2 - \frac{a(1-a)}{n_a - 1}\right) + E\left(b^2 - \frac{b(1-b)}{n_b - 1}\right) - 2E(ab) \\ &= (a')^2 - 2a'b' + (b')^2 \\ &= (a' - b')^2 \\ &= F_2(a', b') \end{aligned} \tag{15}$$

where we have used the two identities defined above in equations 9 and 12.

Similarly, we have

$$\begin{aligned} F_3(a,b,c) &= E\left((c-a)(c-b) - \frac{c(1-c)}{n_c - 1}\right) \\ &= E\left(c^2 - ac - bc - ab - \frac{c(1-c)}{n_c - 1}\right) \\ &= E\left(c^2 - \frac{c(1-c)}{n_c - 1}\right) - E(ac) - E(bc) - E(ab) \\ &= (c')^2 - a'c' - b'c' - a'b' \\ &= (c' - a')(c' - b'). \end{aligned} \tag{16}$$

#### 1.2 Pseudo-haploid sample sizes and bias-correction

So, as we see, for estimators to be unbiased, we need to subtract terms from F2 and F3 statistics, which depend on the sample size. More specifically, on the number of chromosomes sampled to obtain allele frequency estimates. The usual case with high-quality diploid genetic data is that given a sample from $n_d$ diploid individuals, there are $2n_d$ chromosomes. However, with low-quality ancient DNA, a common scheme to obtain genotype estimates is a simple random-sampling scheme of sequencing reads aligning to a given SNP position.

In such a haploid sampling scheme, the resulting number of "chromosomes" that one uses for estimating allele frequencies is $n_d$ instead of $2n_d$.

In `xerxes`, we make use of the "Genotype_Ploidy" column in the .janno File, to compute the correct number of chromosomes, since the genotype file itself does not provide this information (it is always coded as pseudo-diploid, even if the calling is haploid). If this column is missing, or contains missing data, `xerxes` will output a warning and assume that the data is fully diploid!

### 2 Beyond F-Statistics

Beyond F-Statistics, we support the following additional ones:

#### 2.1 Heterozygosity

Heterozygosity is defined using population allele frequencies as:

$$\mathrm{Het}(c') = \langle 2c'(1-c') \rangle \tag{17}$$

and on sample allele frequencies:

$$\mathrm{Het}(c) = \left\langle 2c(1-c)\frac{n_c}{n_c - 1} \right\rangle \tag{18}$$

which again can be shown to be unbiased

```
In[5]:= Expectation[2 k/n (1 - k/n) n/(n - 1),
 k \[Distributed] BinomialDistribution[n, aPrime]]

                                  2
Out[5]= -2 (-aPrime + aPrime )
```

#### 2.2 F3star

In addition to `F3` and `F3vanilla`, we also have `F3star`, which is a bias-corrected F3 normalised by the heterozygosity of population C, defined on page 1071 (right) in [1]. Based on population allele frequencies we would have

$$F_3^*(a', b', c') = \frac{\langle (c-a)(c-b) \rangle}{\langle 2c'(1-c') \rangle} \tag{19}$$

and using the unbiased estimators for both the numerator and the denominator, we reproduce the estimator implemented in Patterson's ADMIXTOOLS software:

$$F_3^*(a,b,c) = \frac{\left\langle (c-a)(c-b) - \frac{c(1-c)}{n_c-1} \right\rangle}{\left\langle 2c(1-c)\frac{n_c}{n_c-1} \right\rangle} \tag{20}$$

#### 2.3 $F_{\mathbf{ST}}$

$F_{\mathrm{ST}}$ is a measure for how differentiated two populations are, taking into account internal genetic variation. It is in spirit very similar to $F_2$, in that it becomes 0 if two populations are not differentiated at all (meaning they are effectively the same), and the more differentiated, the more positive is the measure. However, $F_{\mathrm{ST}}$ is explicitly scaled in a way that exposes relative genetic drift. As derived by [3], we here use the so-called Hudson estimator, which is defined as

$$\mathrm{FST}_{\mathrm{vanilla}}(a', b') = \frac{\left\langle (a'-b')^2 \right\rangle}{\langle a'\,(1-b') + (1-a')\,b' \rangle} \tag{21}$$

As discussed in [3], there is no simple unbiased estimator for this complete expression, but there are unbiased estimators for both the numerator and denominator separately:

$$\mathrm{FST}(a,b) = \frac{\left\langle (a-b)^2 - \frac{a(1-a)}{n_a-1} - \frac{b(1-a}{n_b-1} \right\rangle}{\langle a(1-b) + b(1-a) \rangle} \tag{22}$$

and this ratio of unbiased estimators turns out to be asymptotically also unbiased.

#### 2.4 Pairwise mismatch rate

The pairwise mismatch rate (PWM) measures the rate of observing a different allele in two randomly sampled haplotypes. It is typically computed between pairs of individuals, for example to detect close relatives or identical individuals.

Its definition is:

$$\mathrm{PWM}(a,b) = \langle a(1-b) + (1-a)b \rangle$$

### 3 Summary of all supported statistics in xerxes

Here is a list of all statistics that can be computed in xerxes:

| Name (xerxes) | Formula | Bias |
|---|---|---|
| `F2vanilla` | $\left\langle (a-b)^2 \right\rangle$ | Biased |
| `F3vanilla` | $\langle (c-a)(c-b) \rangle$ | Biased |
| `F2` | $\left\langle (a-b)^2 - \frac{a(1-a)}{n_a-1} - \frac{b(1-b)}{n_b-1} \right\rangle$ | Unbiased |
| `F3` | $\left\langle ((c-a)(c-b) - \frac{c(1-c)}{n_c-1} \right\rangle$ | Unbiased |
| `F3star` | $\frac{\left\langle (c-a)(c-b) - \frac{c(1-c)}{n_c-1} \right\rangle}{\left\langle 2c(1-c)\frac{n_c}{n_c-1} \right\rangle}$ | Asymptotically unbbiased |
| `F4` | $\langle (a-b)(c-d) \rangle$ | Unbiased |
| `Het` | $\left\langle 2c(1-c)\frac{n_c}{n_c-1} \right\rangle$ | Unbiased |
| `FSTvanilla` | $\frac{\left\langle (a'-b')^2 \right\rangle}{\langle a'(1-b')+(1-a')b' \rangle}$ | Biased |
| `FST` | $\frac{\left\langle (a-b)^2 - \frac{a(1-a)}{n_a-1} - \frac{b(1-a}{n_b-1} \right\rangle}{\langle a(1-b)+b(1-a) \rangle}$ | Asympotically unbiased |
| `PWM` | $\langle a(1-b) + (1-a)b \rangle$ | Unbiased |

### 4 Jackknife-Estimation of the standard error

#### 4.1 Theory

All of the above statistics are computed genome-wide, but we use a Block-Jackknife approach to estimate errors. This approach was also popularised in Patterson's software ADMIXTOOLS, and is based on a method proposed by [4].

The key idea is to evaluate the statistics not only genome-wide, but also for subsets of the genome. For these subsets, consider the entire data being divided into $g$ consecutive blocks, of equal or different sizes of $m_j$ sites in each block, where $j = 1 \dots g$.

Let

$$n = \sum_{j=1}^{g}$$

be the total number of sites.

Let $\theta_n$ be the genome-wide estimate based on all $n$ sites, then we define $n$ partial estimates, each with one block removed. We use the notation $\theta_{-j}$ to denote the estimate of the statistics applied a partial dataset with the $j$th block removed.

We then first define the Jackknife estimate:

$$\theta_J = g\theta_n - \sum_{j=1}^{g} (1 - m_j/n)\theta_{-j}$$

Note that for many statistics, one can show that $\theta_J = \theta_n$, but that is not in general the case, and in fact not for all of the above statistics. In `xerxes fstats`, we report both the genome-wide estimate $\theta_n$ (called "Estimate_Full") and the Jackknife estimate $\theta_J$ (called "Estimate_Jackknife") in the output table. In general, for ease of communication and clarity of methods, the genome-wide estimate should be reported as the actual estimate.

We also can derive an estimate of the standard error $\sigma_J$ of the Jackknife estimate (which we take to be also the standard error of the total estimate). As derived in [4], it is given by:

$$
\begin{aligned}
h_j &= \frac{n}{m_j} \\
p_j &= h_j \theta_n - (h_j - 1)\theta_{-j} \\
\sigma_J^2 &= \frac{1}{g} \sum_{j=1}^{g} \frac{1}{h_j - 1} (p_j - \theta_J)^2
\end{aligned}
\tag{23}
$$

Finally, we compute Z-scores for statistics as

$$Z = \frac{\theta_J}{\sigma_J}$$

#### 4.2 Implementation

We implement two ways to define blocks in `xerxes fstats`. First, simply by chunks of a given number of SNPs (by default 5000), as for example specified using the parameter `--jackknife 5000`. Second, by entire chromosomes, as specified using the special declaration `--jackknife CHR`. In practice, we have not seen great differences between the two approaches, and the default is to chunk by 5000 SNPs.

The program takes the *total number of sites* in each block as the basis for the weights $m_j$, not just the sites contributing to a given statistic, which could exclude sites due to missing data. This choice was made to make the computation more reproducible and comparable across multiple statistics. Note that this does not mean that missingness is ignored. Of course, missingness still contributes to variance seen throughout the genome, and this gets reflected in the Jackkknife estimation of the standard error. We just decided to fix the weights themselves by the total number of sites, ignoring missingness for the specific couting of $m_j$.

### 5 Frequency-Ascertainment

This is an experimental feature that will be described properly in a forthcoming preprint, and then will be added also to this whitepaper.
