## Supplementary Text 6 - Qjanno for "Poseidon – A framework for archaeogenetic human genotype data management"

### Guide for qjanno v1.0.0.0

#### Contents

**1 Background** **1**

**2 How does this work?** **1**

**3 Installation** **2**

**4 The CLI interface** **2**

**5 Query examples** **5**

### 1 Background

Qjanno started as a fork of the qhs software tool, which was, in turn, inspired by the command line tool q. All of them enable SQL queries on delimiter-separated text files (e.g. .csv or .tsv). For `qjanno`, we copied the source code of qhs v0.3.3 (MIT-License) and adjusted it to provide a smooth experience with a special kind of .tsv file: The Poseidon `.janno` file.

Unlike `trident` or `xerxes`, `qjanno` does not have a complete understanding of the `.janno` file structure, and (mostly) treats `.janno` files like normal .tsv files. It does not validate them upon reading and takes them at face value. Still `.janno` files are given special consideration: With a set of pseudo-functions in the `FROM` field of the SQL query they can be searched recursively and loaded together into one table.

`qjanno` still supports most features of qhs, so it can still read arbitrary .csv and .tsv files independently or in conjunction with `.janno` files (e.g. for `JOIN` operations).

### 2 How does this work?

On startup, `qjanno` creates an SQLite [1] database in memory. It then reads the requested, structured text files and attributes each column a type (either character or numeric). With this annotation it can write the contents of the files to tables in the in-memory database. `qjanno` finally sends the user-provided SQL query to said database, waits for the result, parses it and returns it on the command line.

The SQL query gets pre-parsed to extract file names and then forwarded to an SQLite database server via the

Haskell library sqlite-simple. That means `qjanno` can parse and understand basic SQLite3 syntax, though not everything. `PRAGMA` functions, for example, are not available. The examples below show some of the available syntax, but they are not exhaustive. Trial and error is recommended to see what does and what does not work. Please report missing expected functionality at our issue board on GitHub.

#### 3 Installation

See the Poseidon website (https://www.poseidon-adna.org/#/qjanno) or the GitHub repository (https://github.com/poseidon-framework/qjanno) for up-to-date installation instructions.

#### 4 The CLI interface

This is the CLI interface of `qjanno`:

```
Usage: qjanno [--version] [QUERY] [-q|--queryFile FILE] [-c|--showColumns]
              [-t|--tabSep] [--sep DELIM] [--noHeader] [--raw] [--noOutHeader]

  Command line tool to allow SQL queries on .janno (and arbitrary .csv and .tsv)
  files.

Available options:
  -h,--help                Show this help text
  --version                Show qjanno version
  QUERY                    SQLite syntax query with paths to files for table
                           names. See the online documentation for examples. The
                           special table name syntax 'd(path1,path2,...)' treats
                           the paths (path1, path2, ...) as base directories
                           where .janno files are searched recursively. All
                           detected .janno files are merged into one table and
                           can thus be subjected to arbitrary queries.
  -q,--queryFile FILE      Read query from the provided file.
  -c,--showColumns         Don't run the query, but show all available columns
                           in the input files.
  -t,--tabSep              Short for --sep $'\t'.
  --sep DELIM              Input file field delimiter. Will be automatically
                           detected if it's not specified.
  --noHeader               Does the input file have no column names? They will
                           be filled automatically with placeholders of the form
                           c1,c2,c3,...
  --raw                    Return the output table as tsv.
  --noOutHeader            Remove the header line from the output.
```

This help can be accessed with `qjanno -h`. Running `qjanno` without any parameters does not work: The `QUERY` parameter is mandatory and without it the tool will fail with the exception `Query cannot be empty`.

#### 4.1 A basic example

A basic, working qjanno query could look like this:

```
$ qjanno "SELECT package_title,Poseidon_ID,Country \
          FROM d(2010_RasmussenNature,2012_MeyerScience)"
.----------------------.----------------------.-----------.
|    package_title     |     Poseidon_ID      |  Country  |
:======================:======================:===========:
| 2010_RasmussenNature | Inuk.SG              | Greenland |
| 2012_MeyerScience    | A_Mbuti-5.DG         | Congo     |
| 2012_MeyerScience    | A_Yoruba-4.DG        | Nigeria   |
| 2012_MeyerScience    | A_Sardinian-4.DG     | Italy     |
| 2012_MeyerScience    | A_French-4.DG        | France    |
| 2012_MeyerScience    | A_Dinka-4.DG         | Sudan     |
| 2012_MeyerScience    | A_Ju_hoan_North-5.DG | Namibia   |
'----------------------'----------------------'-----------'
```

Running qjanno with this query triggers the following process:

1. With d(...) in the FROM field, qjanno searches recursively for package-defining POSEIDON.yml files in the given base directories 2010_RasmussenNature and 2012_MeyerScience.
2. It finds the .yml files and reads some of their fields, including the title, the packageVersion and the jannoFile path. It then selects the latest version of each package.
3. With the relevant .janno file paths available, qjanno reads them, appends the package_title, package_version and source_file columns, merges them (with a simple row-bind), and orders their columns.
4. It then writes the resulting .janno table to the SQLite database in memory.
5. Now the actual query gets sent to the database server to execute it. In this case the SELECT statement includes three variables (column names): package_title, Poseidon_ID and Country. The database server returns these three columns for the merged .janno table.
6. qjanno finally prints the result in a clean, human readable format to the standard output.

#### 4.2 The .janno-crawling pseudo-functions

d(...) is one of four pseudo-functions to search and load .janno files in the FROM field of the query:

- d(<path_to_directory1>,<path_to_directory2>,...): With d(), qjanno (recursively) searches all package-defining POSEIDON.yml files in all listed directories and reads them to determine the latest package version. It then reads the .janno files associated with these latest package versions.
- da(<path_to_directory1>,<path_to_directory2>,...): da() behaves just as d(), but it does not filter for the latest package version: It loads all packaged .janno files.
- j(<path_to_directory1>,<path_to_directory2>,...): j() simply searches for files with the extension .janno in all listed directories and loads them regardless of whether they are part of a Poseidon package or not.
- <path_to_one_janno_file>.janno: Specific .janno files can be listed individually. They are identified as such by their .janno extension.

Multiple of these methods can be combined as a comma-separated list. Each respective mechanism then yields a list of `.janno` file paths, and the list of lists is flattened to a simple list of paths. `qjanno` then reads all files in this combined list, merges them and makes them available for querying in the in-memory SQLite database.

Note that the `FROM` field must not include any spaces – even in a comma-separated list. `qjanno` parses the `QUERY` using space as a separator.

#### 4.3 CLI details

`qjanno` can not just read `.janno` files, but also arbitrary .csv and .tsv files. This option is triggered by providing file names (relative paths) in the `FROM` field of the query.

```
$ echo -e "Col1,Col2\nVal1,Val2\nVal3,Val4\n" > test.csv
$ qjanno "SELECT * FROM test.csv"
.-------------.------.------.
| source_file | Col1 | Col2 |
:=============:======:======:
| test.csv    | Val1 | Val2 |
| test.csv    | Val3 | Val4 |
'-------------'------'------'
```

For these non-`.janno` files `qjanno` tries to automatically determine the relevant delimiter. With `--sep` a delimiter can be specified explicitly, and the shortcut `-t` sets `--sep $'\t'` for tab-separated files.

```
$ echo -e "Col1\tCol2\nVal1\tVal2\nVal3\tVal4\n" > test.tsv
$ qjanno "SELECT * FROM test.tsv" -t
.--------------.------.------.
| source_file  | Col1 | Col2 |
:==============:======:======:
| test_tab.csv | Val1 | Val2 |
| test_tab.csv | Val3 | Val4 |
'--------------'------'------'
```

The `--noHeader` option allows to read files without headers, so column names. The columns are then automatically named *c1,c2,... cN*:

```
$ echo -e "Val1,Val2\nVal3,Val4\n" > test.csv
$ qjanno "SELECT c1,c2 FROM test.csv" --noHeader
.------.------.
|  c1  |  c2  |
:======:======:
| Val1 | Val2 |
| Val3 | Val4 |
'------'------'
```

The remaining options concern the output: `--raw` returns the output table not in the ornate, human-readable ASCII table layout, but in a simple tab-separated format. `--noOutHeader` omits the header line in the output.

```
$ echo -e "Col1,Col2\nVal1,Val2\nVal3,Val4\n" > test.csv
$ qjanno "SELECT * FROM test.csv" --raw --noOutHeader
test.csv  Val1  Val2
```

```
test.csv  Val3  Val4
```

Note that these output options can be combined to directly prepare individual lists in `trident`'s `forge` selection language format:

```
$ qjanno "SELECT '<'||Poseidon_ID||'>' FROM d(2012_MeyerScience)" --raw --noOutHeader
<A_Mbuti-5.DG>
<A_Yoruba-4.DG>
<A_Sardinian-4.DG>
<A_French-4.DG>
<A_Dinka-4.DG>
<A_Ju_hoan_North-5.DG>
```

##### 4.4 The `-c|--showColumns` option

`-c|--showColumns` is a special option that, when activated, makes `qjanno` return not the result of a given query, but an overview table with the columns available in all selected files for said query. That is helpful to get an overview what could actually be queried.

```
$ echo -e "Col1,Col2\nVal1,Val2\nVal3,Val4\n" > test.csv
$ qjanno "SELECT * FROM test.csv" -c
.-------------.----------.-------------------.
|   Column    |   Path   | qjanno Table name |
:=============:==========:===================:
| source_file | test.csv | test              |
| Col1        | test.csv | test              |
| Col2        | test.csv | test              |
'-------------'----------'-------------------'
```

This summary also includes the artificial, structurally cleaned table names assigned by `qjanno` before writing to the SQLite database. Often we can not simply use the file names as table names, because SQLite has strict naming requirements. File names or relative paths are generally invalid as table names and therefore need to be replaced with an adjusted string. These artificially generated names are mostly irrelevant from a user perspective – except a query involves multiple files, e.g. in a `JOIN` operation. See below for an example.

#### 5 Query examples

The following examples show some of the functionality of the SQLite query language available through `qjanno`. See the SQLite syntax documentation for more details. The examples were prepared and tested in a clone of the Poseidon community archive.

###### Sub-setting with `WHERE`

Get all individuals/samples (.janno rows) in two Poseidon packages where UDG is set to 'minus'. That means the underlying aDNA libraries were subjected to a lab protocol without UDG (USER enzyme) treatment before sequencing.

```
$ qjanno " \
SELECT package_title,Poseidon_ID,UDG \
```

```
FROM d(2010_RasmussenNature,2012_MeyerScience) \
WHERE UDG = 'minus' \
"
.-------------.-------.
| Poseidon_ID |  UDG  |
:=============:=======:
| Inuk.SG     | minus |
'-------------'-------'
```

Note that rows where the UDG entry is missing (`NULL`) are silently dropped here.

Get all individuals where Genetic_Sex is not ‘F’ (female) **and** Country is ‘Sudan’.

```
$ qjanno " \
SELECT Poseidon_ID,Country \
FROM d(2010_RasmussenNature,2012_MeyerScience) \
WHERE Genetic_Sex <> 'F' AND Country = 'Sudan' \
"
.--------------.---------.
| Poseidon_ID  | Country |
:==============:=========:
| A_Dinka-4.DG | Sudan   |
'--------------'---------'
```

Get all individuals where the UDG column is not `NULL` (not missing) **or** the Country is ‘Sudan’.

```
$ qjanno " \
SELECT Poseidon_ID,Country \
FROM d(2010_RasmussenNature,2012_MeyerScience) \
WHERE UDG IS NOT NULL OR Country = 'Sudan' \
"
.--------------.-----------.
| Poseidon_ID  |  Country  |
:==============:===========:
| Inuk.SG      | Greenland |
| A_Dinka-4.DG | Sudan     |
'--------------'-----------'
```

Get all individuals where Nr_SNPs is equal to or bigger than 600,000.

```
$ qjanno " \
SELECT Poseidon_ID,Nr_SNPs \
FROM d(2010_RasmussenNature,2012_MeyerScience) \
WHERE Nr_SNPs >= 600000 \
"
.-------------.---------.
| Poseidon_ID | Nr_SNPs |
:=============:=========:
| Inuk.SG     | 1101700 |
```

```
'-------------'---------'
```

**Ordering with `ORDER BY`**

Order all individuals by Nr_SNPs.

```
$ qjanno " \
SELECT Poseidon_ID,Nr_SNPs \
FROM d(2010_RasmussenNature,2012_MeyerScience) \
ORDER BY Nr_SNPs \
"
.----------------------.---------.
|     Poseidon_ID      | Nr_SNPs |
:======================:=========:
| A_French-4.DG        | 592535  |
| A_Ju_hoan_North-5.DG | 593045  |
| A_Mbuti-5.DG         | 593057  |
| A_Dinka-4.DG         | 593076  |
| A_Yoruba-4.DG        | 593097  |
| A_Sardinian-4.DG     | 593109  |
| Inuk.SG              | 1101700 |
'----------------------'---------'
```

Order all individuals by Date_BC_AD_Median in a descending (`DESC`) order. Date_BC_AD_Median includes missing values.

```
$ qjanno " \
SELECT Poseidon_ID,Date_BC_AD_Median \
FROM d(2010_RasmussenNature,2012_MeyerScience) \
ORDER BY Date_BC_AD_Median DESC \
"
.----------------------.-------------------.
|     Poseidon_ID      | Date_BC_AD_Median |
:======================:===================:
| Inuk.SG              | -1935             |
| A_Sardinian-4.DG     |                   |
| A_Yoruba-4.DG        |                   |
| A_Dinka-4.DG         |                   |
| A_Mbuti-5.DG         |                   |
| A_Ju_hoan_North-5.DG |                   |
| A_French-4.DG        |                   |
'----------------------'-------------------'
```

**Reducing the number of return values with `LIMIT`**

Only return the first three result individuals.

```
$ qjanno " \
SELECT Poseidon_ID,Group_Name \
```

```
FROM d(2010_RasmussenNature,2012_MeyerScience) \
LIMIT 3 \
"
.---------------.-----------------------------.
|  Poseidon_ID  |         Group_Name          |
:===============:=============================:
| Inuk.SG       | Greenland_Saqqaq.SG         |
| A_Mbuti-5.DG  | Ignore_Mbuti(discovery).DG  |
| A_Yoruba-4.DG | Ignore_Yoruba(discovery).DG |
'---------------'-----------------------------'
```

##### Combining tables with `JOIN`

For `JOIN` operations, SQLite requires table names to specify which columns are meant when combining multiple tables with overlapping column names. See the option `-c|--showColumns` to get the relevant table names as generated from the input file paths.

```
$ echo -e "Poseidon_ID,MoreInfo\nInuk.SG,5\nA_French-4.DG,3\n" > test.csv

$ qjanno "SELECT * FROM d(2010_RasmussenNature,2012_MeyerScience)" -c
.------------------------------.-------------------------------------------.
|            Column            |                   Path                    |
:==============================:===========================================:
| package_title                | d(2010_RasmussenNature,2012_MeyerScience) |
| package_version              | d(2010_RasmussenNature,2012_MeyerScience) |
| source_file                  | d(2010_RasmussenNature,2012_MeyerScience) |
| Poseidon_ID                  | d(2010_RasmussenNature,2012_MeyerScience) |
...
--------------------------------------.
          qjanno Table name           |
======================================:
 d2010RasmussenNature2012MeyerScience |
 d2010RasmussenNature2012MeyerScience |
 d2010RasmussenNature2012MeyerScience |
 d2010RasmussenNature2012MeyerScience |
...

$ qjanno "SELECT * FROM test.csv" -c
.-------------.----------.-------------------.
|   Column    |   Path   | qjanno Table name |
:=============:==========:===================:
| source_file | test.csv | test              |
| Poseidon_ID | test.csv | test              |
...
```

Join the `.janno` files with the information in the test.csv file (by the `Poseidon_ID` column).

```
$ qjanno " \
SELECT d2010RasmussenNature2012MeyerScience.Poseidon_ID,Country,MoreInfo \
FROM d(2010_RasmussenNature,2012_MeyerScience) \
INNER JOIN test.csv \
ON d2010RasmussenNature2012MeyerScience.Poseidon_ID = test.Poseidon_ID \
"
.---------------.-----------.----------.
|  Poseidon_ID  |  Country  | MoreInfo |
:===============:===========:==========:
| Inuk.SG       | Greenland | 5        |
| A_French-4.DG | France    | 3        |
'---------------'-----------'----------'
```

##### Grouping data and applying aggregate functions

SQLite provides a number of aggregation functions: `avg(X)`, `count(*)`, `count(X)`, `group_concat(X)`, `group_concat(X,Y)`, `max(X)`, `min(X)` and `sum(X)`. See the documentation here. These functions shine especially when combined with the `GROUP BY` operation.

Determine the minimal number of SNPs across all individuals.

```
$ qjanno "SELECT min(Nr_SNPs) AS n FROM d(2010_RasmussenNature,2012_MeyerScience)"
.--------.
|   n    |
:========:
| 592535 |
'--------'
```

Count the number of individuals per Date_Type group and calculate the average Nr_SNPs for both groups.

```
$ qjanno " \
SELECT Date_Type,count(*),avg(Nr_SNPs) \
FROM d(2010_RasmussenNature,2012_MeyerScience) \
GROUP BY Date_Type \
"
.-----------.----------.--------------.
| Date_Type | count(*) | avg(Nr_SNPs) |
:===========:==========:==============:
| C14       | 1        | 1101700.0    |
| modern    | 6        | 592986.5     |
'-----------'----------'--------------'
```

---

[1] K. P. Gaffney, M. Prammer, L. Brasfield, D. R. Hipp, D. Kennedy, and J. M. Patel, "SQLite: Past, present, and future," *Proceedings of the VLDB Endowment*, vol. 15, no. 12, pp. 3535–3547, Aug. 2022, doi: 10.14778/3554821.3554842.
