## Supplementary Text 7 - Janno R package for "Poseidon – A framework for archaeogenetic human genotype data management"

### Guide for the janno R package v1.0.0

#### Contents

**1 Installation** 1

**2 Read `.janno` files** 1

**3 Validate `.janno` files** 2

**4 Write `janno` objects back to `.janno` files** 2

**5 Process age information in `janno` objects** 2

**6 General helper functions** 3

### 1 Installation

See the Poseidon website (https://www.poseidon-adna.org/#/janno_r_package) or the GitHub repository (https://github.com/poseidon-framework/janno) for up-to-date installation instructions.

### 2 Read `.janno` files

You can read `.janno` files with

```
my_janno_object <- janno::read_janno(
  path = "path/to/my/janno_file.janno",
  to_janno = TRUE,
  validate = TRUE
)
```

The path argument takes one or multiple file paths or directory paths. `read_janno()` searches recursively for `.janno` files in these directory paths.

Before loading the `.janno` files they are validated with `janno::validate_janno()`. You can avoid this potentially time consuming step with `validate = FALSE`.

Usually the `.janno` files are first loaded as normal `.tsv` files with every column type set to `character` and then the columns are transformed to the specified types. This transformation can be turned off with `to_janno =`

FALSE.

`read_janno()` returns an object of class `janno`. This class is derived from the `tibble` class, which integrates well with the tidyverse [1] and its packages, e.g. `dplyr` or `ggplot2`.

#### 3 Validate .janno files

You can validate `.janno` files with

```
my_janno_issues <- janno::validate_janno("path/to/my/janno_file.janno")
```

`validate_janno` returns a `tibble` with issues in the respective `.janno` files. For edge cases this validation may yield slightly different results than `trident validate`.

#### 4 Write janno objects back to .janno files

`janno` objects usually contain list columns, that can not directly be written to a flat text file like the `.janno` file. The function `write_janno` solves that. It employs a helper function `flatten_janno()`, which translates list columns to the string list format in `.janno` files (so: multiple values for one cell separated by `;`).

This only works for vector list columns, so when each cell contains a vector of values. If a list column contains other data structures, e.g. `data.frame`s, they will be dropped and replaced with the NULL value `n/a` in the resulting `.janno` file.

```
janno::write_janno(
  my_janno_object,
  path = "path/to/my/new/janno_file.janno"
)
```

#### 5 Process age information in janno objects

`.janno` files contain age information in multiple different columns. See the `.janno` file specification and documentation for a list and detailed explanations of these variables. The function `janno::process_age()` works with this age information to calculate different derived columns, which are then added to the input `janno` object.

You can run it with

```
janno::process_age(
  my_janno_object,
  choices = c("Date_BC_AD_Prob", "Date_BC_AD_Median_Derived", "Date_BC_AD_Sample"),
  n = 100,
  cal_curve = "intcal20"
)
```

`process_age()` includes calibration of radiocarbon dates with the Bchron R package [2]. The calibration curve set in `cal_curve` is applied for every date in the `janno` object. If there are multiple radiocarbon dates for one sample they are automatically combined as the normalized sum of all individual post-calibration probability distributions.

The `choices` argument contains the list of columns that should be calculated and added by `process_age()`. `n` is the number of samples that should be drawn for `Date_BC_AD_Sample`.

##### 5.1 Output column `Date_BC_AD_Prob`

`Date_BC_AD_Prob` is a list column with a `data.frame` for each `janno` row, so each individual/sample. This `data.frame` stores a density distribution (`sum_dens`) over a set of years BC/AD (`age`). Additionally the boolean column `two_sigma` documents if a given year is within the 2-sigma high-density regions of the distribution. `center` is also a boolean column with only one `TRUE` value for the year that corresponds to the calibrated median age of the sample.

| age | sum_dens | two_sigma | center |
|---|---|---|---|
| -1506 | 0.00000456 | FALSE | FALSE |
| -1505 | 0.00000622 | FALSE | FALSE |
| -1504 | 0.00000907 | FALSE | FALSE |
| ... | ... | ... | ... |

The density distributions are either the result of (sum) calibration on radiocarbon dates or – for samples that are only contextually dated – a uniform distribution over the archaeologically determined age range.

##### 5.2 Output column `Date_BC_AD_Median_Derived`

`Date_BC_AD_Median_Derived` is a simple integer column with the median age (in years BC/AD) as determined from `Date_BC_AD_Prob`.

##### 5.3 Output column `Date_BC_AD_Sample`

`Date_BC_AD_Sample` is again a list column with a vector of `n` ages (in years BC/AD) for each `.janno` file individual/sample. These ages are randomly drawn with `base::sample(prob = ...)` using the probability distribution calculated for `Date_BC_AD_Prob`.

#### 6 General helper functions

When you are preparing a `.janno` file and want to determine the entries for the columns `Date_BC_AD_Median`, `Date_BC_AD_Start` and `Date_BC_AD_Stop` from radiocarbon dates, then `janno::quickcalibrate()` might come in handy.

```
janno::quickcalibrate(ages, sds)
```

`ages` takes a list of uncalibrated C14 ages BP and `sds` a list of the respective standard deviations. If multiple ages are provided for one sample, then the function automatically performs a sum calibration.

`quickcalibrate(list(1000, c(2000, 2200)), list(20, c(30, 40)))` for example returns a `data.frame` like this:

| Date_BC_AD_Start_2Sigma | ... | Date_BC_AD_Median | ... | Date_BC_AD_Stop_2Sigma |
|---|---|---|---|---|
| 994 | ... | 1029 | ... | 1149 |

| Date_BC_AD_Start_2Sigma | ... | Date_BC_AD_Median | ... | Date_BC_AD_Stop_2Sigma |
|---|---|---|---|---|
| -383 | ... | -88 | ... | 117 |

This output can be copied to a `.janno` file, where `Date_BC_AD_Start_2Sigma` corresponds to `Date_BC_AD_Start`, and `Date_BC_AD_Stop_2Sigma` to `Date_BC_AD_Stop`.

---

[1] H. Wickham *et al.*, "Welcome to the Tidyverse," *Journal of Open Source Software*, vol. 4, no. 43, p. 1686, Nov. 2019, doi: 10.21105/joss.01686.

[2] J. Haslett and A. Parnell, "A simple monotone process with application to radiocarbon-dated depth chronologies," *Journal of the Royal Statistical Society Series C: Applied Statistics*, vol. 57, no. 4, pp. 399–418, May 2008, doi: 10.1111/j.1467-9876.2008.00623.x.
