## Supplementary Text 8 - Archive Comparison for "Poseidon – A framework for archaeogenetic human genotype data management"

### Supplementary Text: Comparison of the public archive content

### 1 Comparing PCA and PAA

Comparing the content of the Poseidon Community Archive (PCA) and the AADR Archive (PAA) is a non-trivial task, given the nature of ancient DNA data and specific peculiarities of each archive. The following sections explain the subfigures A-F of the main text Figure 5 and highlight some of the immanent complexity of the comparison. The code behind this analysis is available on GitHub here: `https://github.com/nevrome/poseidon.analysis.2024` and in a long-term archive here: `https://doi.org/10.17605/OSF.IO/ZUQGB`. The following R packages were used: bib2df [1], cowplot [2], ggpattern [3], ggrepel [4], ggsankey [5], giscoR [6], hash [7], janno, sf [8], wesanderson [9] and various packages from the tidyverse [10].

We here consider the state of the archives on 2024-03-15. For the PAA this equates to v54.1.p1 of the AADR [11], which it is split in the PAA into four Poseidon packages.

#### 1.1 Mapping of publications to Poseidon packages

One fundamental difference between the PAA and the PCA emerges from the package structure. The PAA combines many publications in a small number of thematic packages, whereas the PCA (and PMA) generally feature packages that each correspond to exactly one publication. This is well visible in Supplementary Figure 1. Two notable exceptions in the PCA include reference genomes for archaic human populations and special (e.g. non-human) modern reference genomes.

The AADR in version 54.1.p1 and previous versions includes two big datasets: A *1240K* dataset where all samples are provided with genotype data mapped to the 1240k [12] SNP-set and a *HO* dataset where the genotype data only includes the SNPs on the HumanOrigins [13] array. The HO dataset is a superset of the 1240K one, as it includes all samples of the latter, plus a large number of modern human reference samples for which only HumanOrigins data is publicly available. For the PAA these two datasets were split into four Poseidon packages. This was mainly done due to a technical limitation of the GitHub LFS system on which we rely for the public archives: It only allows a maximal single file size of 2GB (`https://docs.github.com/en/repositories/working-with-files/managing-large-files/about-git-large-file-storage` [2024-03-26]). Among these four packages, one includes modern reference data for which 1240k data is available, one for which only HumanOrigin SNPs are known, and two feature ancient data split by geography (Europe vs. beyond Europe).

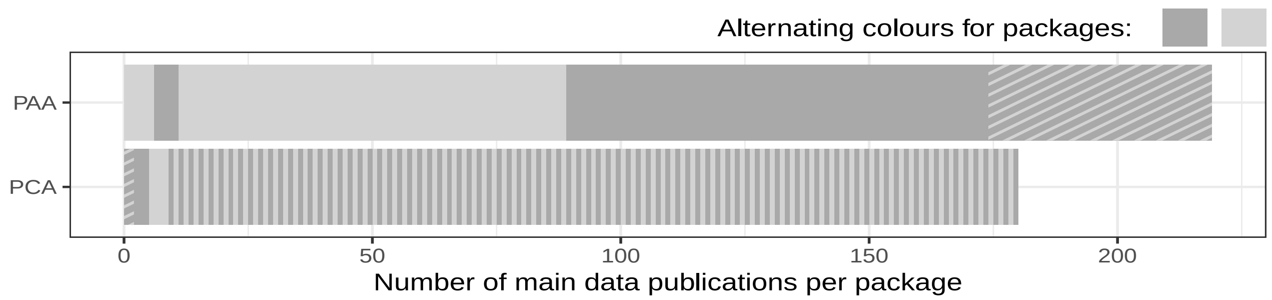

**Supplementary Figure 1:** Corresponds to Figure 5A in the main text.

This setup produces a lot of overlap between packages, thus potentially inflating the total number of publications in the figure, where publications are counted by packages. To correct for that the diagonally hatched section summarises publications that appear in more than one package to count them only once. As a consequence of their construction the double-counting in the PAA is high, and low for the PCA.

Note that this figure only considers the main publication per sample, so the primary publication listed in the `Publication` column of the .janno file for each sample.

#### 1.2 Samples, individuals and identifiers

Where the comparison in section 1.1 only considers publications, the following figures count on the level of human individuals and samples. When we speak about the main operational entities in a Poseidon package we use either *individual* or *sample* throughout the software, its documentation and even this publication, but both of these terms are actually inaccurate. This requires some clarification:

Generally, archaeogenetics operates on burial contexts, e.g. graves with one or multiple ancient human individuals. Usually, though not always, it is possible to attribute the skeletal remains within these graves to individuals based on the archaeological context and physical-anthropological analysis. Each individual can get sampled one or multiple times, either by directly probing their preserved tissue, mostly bones, or by sampling any reagent that contains their DNA (through whatever pathway or taphonomic process). From one such sample one or multiple extracts can be derived, which can be transformed into one or multiple libraries, which may or may not be subjected to a DNA capture protocol and then sequenced one or multiple times. The raw sequencing data can undergo various different forms of computational processing and eventually genotyping to produce the data relevant for most derived analyses and thus stored in Poseidon. While the wetlab-processes can be understood as a relatively predictable tree of separate physical and digital products for any given ancient individual, the computational data-processing finally breaks the conceptual tree-ness by allowing for arbitrary conflation of sequencing data obtained through potentially separate means: Data from different libraries can very well be merged if they are from the same individual, even if they are not from the same sample.

A Poseidon_ID, and therefore the identifier for the main singular entity in a Poseidon package, could approximately be described as representing one end-point in the data preparation graph laid out above. Typically this end-point corresponds to an optimal result, consciously selected for a given individual, research question and publication. Unfortunately, in reality a Poseidon_ID is not suited to uniquely identify exactly one such end-point. The reality in the Poseidon ecosystem is rather that slightly different end-points can have the same Poseidon_ID, e.g. across package versions or public Poseidon archives. A single endpoint can only be uniquely identified from a combination of Poseidon_ID, Poseidon package and package version.

As a consequence, comparing the data coverage of PCA and PAA is difficult. The same end-points can have different Poseidon_IDs, different Poseidon_IDs can describe the same or a very similar end-points and end-points may not even be a particularly helpful entity for comparison. A general identifier for the underlying ancient human individuals would mitigate the issue, but Poseidon does not include or specify such an identifier yet.

For the following analysis we address this issue as follows: Each entity in each of the archives features a Poseidon_ID, which is unique within the archive. Within the archives the Poseidon_IDs are thus sufficient for a reliable tally of the number of entities stored in them. We will refer to these as *sample*-wise counts. For the comparison across the archives we compiled a simplified identifier string from the Poseidon_IDs, which avoids many of the idiosyncracies paper authors and the authors of the AADR dataset introduced to express a specific processing end-point on top of the basic sample/individual identifier. For example we removed suffixes like `.HO`, `.SG`, `_noUDG`, `_petrous`, which highlight different aspects of the data preparation process, `_contam` or `.cont`, which point to data quality issues, and `_published` and `_in.preparation` which communicate something about the a sample's publication history. See the code for a comprehensive list of suffixes and other modifications removed in our processing. We used the thus generated, simplified identifier as a proxy for the underlying individuals and visualize its count explicitly in the following figures. It enables an approximate, potentially individual-level comparison between the archives for the figures Supplementary Figure 3 and Supplementary Figure 5, so we will refer to this as an *individual*-level count.

Figure 5 of the main text only shows the *individual*-level count, whereas the versions here in the supplement also include the larger *sample*-wise counts by visualizing the difference between both with diagonally hatched sections.

#### 1.3 Publication coverage

To understand to which degree PCA and PAA include publications with ancient human genotype data, and thus how well they each cover the field of archaeogenetics, we constructed Supplementary Figure 2 and Supplementary Figure 3.

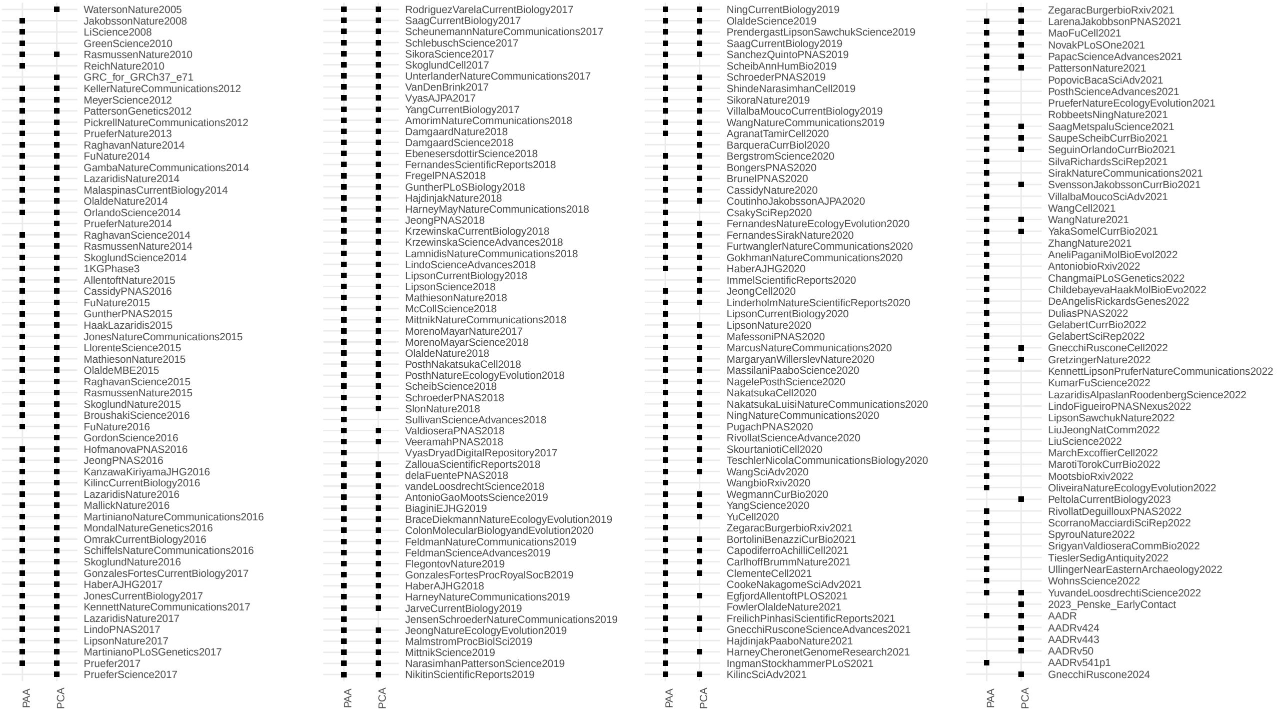

**Supplementary Figure 2:** Publications covered by PCA and PAA. The identifiers are taken from the PAA, except a given publication is only available in the PCA. The publications are ordered by year from top to bottom and from left to right.

PCA and PAA do not use an identical set of keys to identify publications. To match them, we first created a lookup table to find the PAA key for the corresponding PCA key. So Supplementary Figure 2 shows for each PAA key (except the publication only occurs in the PCA) if the respective publication is referenced in either archive and thus most likely included there. Note that here, unlike for Supplementary Figure 1, not only the primary, but all publications are considered. This means also older and secondary publications have their appearance.

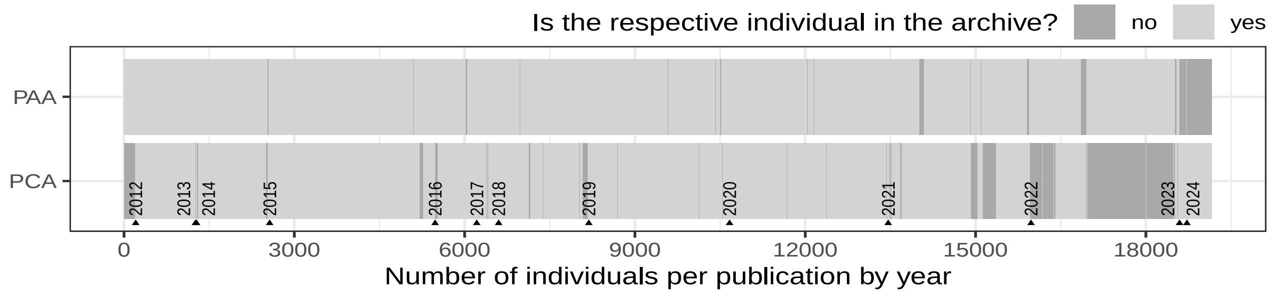

**Supplementary Figure 3:** Corresponds to Figure 5B in the main text.

Supplementary Figure 3 provides a different perspective on the same question. Along the x-axis we sum all individuals, aggregated from the Poseidon_IDs as described in section 1.2 for both archives. The individuals are ordered by publications, which are in turn ordered by the year they were published in. This allows to append a secondary x-axis highlighting the starting points of the individual years. The grey colour scale indicates if a given individual is in an archive or not. Because this is a tally by publication (including secondary publications), some individuals may appear twice or even more often in this figure, if they are linked to multiple publications.

Both figures together show how the PAA is overall more complete, but almost exclusively regarding later publications in 2022. It also references some very old publications from before 2012. For 2023 and 2024 the PCA already includes data which was not yet available at the time when the AADR version represented in the PAA was compiled.

#### 1.4 Data sources and individual-level overlap

The PCA is now meant for author-submitted Poseidon packages, but previously went through a consolidation process, where it originally acted as a catch-all archive for any published archaeogenetic genotype data. Over the last three years we transferred data from the AADR into the PCA multiple times. We documented this process on the package level, and can now count how many data points originate from which AADR version. We can also count how many were added directly from the respective publication or were proper author submissions as intended for the future of the PCA. Supplementary Figure 4 shows the outcome of this tally.

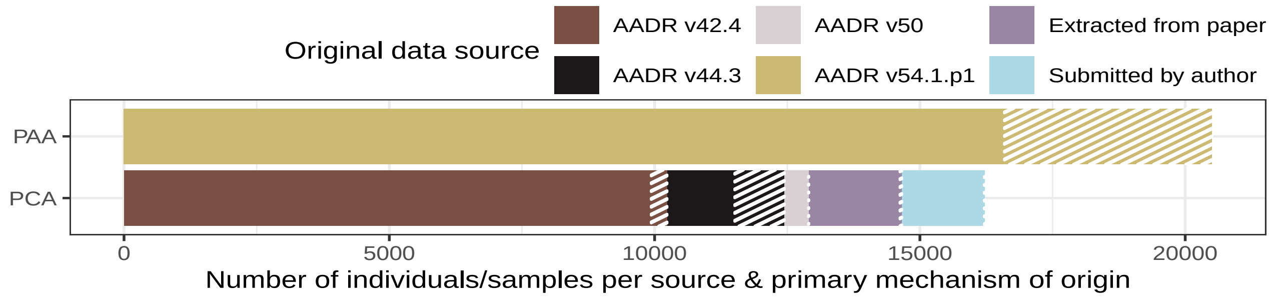

**Supplementary Figure 4:** Corresponds to Figure 5C in the main text.

As laid out in section 1.2 we distinguish two different counts: The number of entities with unique Poseidon_IDs per archive and the aggregated, simplified IDs that serve as a proxy for the number of individuals. In Supplementary Figure 4, and also 6 and 7 below, we mark the individual-proxy-level count with a solid fill, whereas the difference to counting all Poseidon_IDs is highlighted with a hatched fill.

Note that the PAA here exclusively represents the AADR v54.1.p1. A large number of individuals are represented by multiple samples across its four packages, explaining the difference between the two count types. The largest contributing factor to this double-counting is modern reference data (see also Supplementary Figure 6). The PCA consists of a large set of samples originally extracted from the AADR v42.4, another one from v44.3 and then finally v50. Beyond that another set of samples was directly extracted from publications by various contributors within and beyond the Poseidon core team. For these packages the genotype data had to be generated from the raw sequencing data uploaded by the respective authors to the large data archives ENA or SRA. A final and now growing set of samples was directly shared by the paper authors.

This summary begs the question, which of these samples overlap between PAA and PCA. As explained in section 1.2 a comparison on the sample-level (so via Poseidon_ID) is not meaningfully possible, but the individual-level aggregation enables it to some degree.

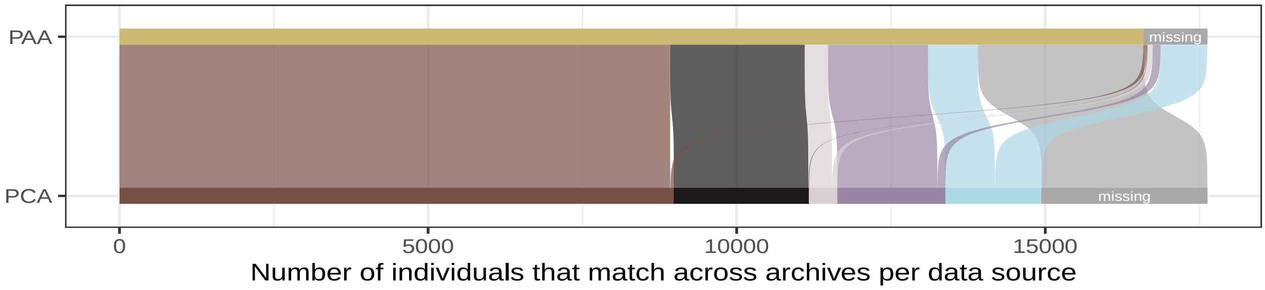

**Supplementary Figure 5:** Corresponds to Figure 5D in the main text.

Supplementary Figure 5 is a basic Sankey diagram that links individuals from PCA and PAA as good as automatically possible. A darkgrey fraction in the figure makes mismatches between the archives explicit.

This figure confirms what already emerged above: The PCA is mostly a subset of the PAA in terms of covered individuals. Only with the latest (during and after 2023) author-submissions the PCA includes a relevant amount of data that is not included in the PAA. The small number of mismatches between the old and the new AADR version is most likely an effect of small name changes and data reorganization either in the AADR over time or in the individual publication-wise packages of the PCA.

#### 1.5 Temporal and spatial information availability

The last two sub-figures in main text Figure 5 summarize the coverage and distribution for two of the most important context information variables: the spatial and temporal origin of a given sample. Both figures include the two counts introduced in section 1.2 with the individual-count shown with a solid and the difference to the sample-count with a hatched fill.

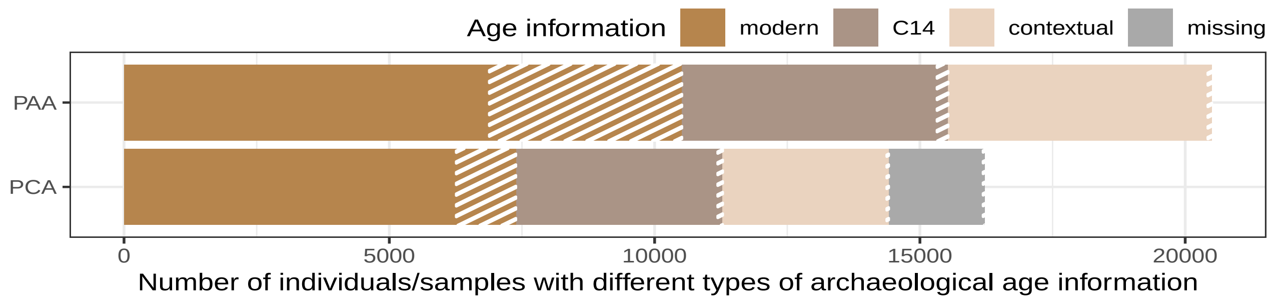

**Supplementary Figure 6:** Corresponds to Figure 5E in the main text.

Poseidon features the .janno column `Date_Type` with the possible values `modern`, `C14` and `contextual` (and of course `n/a` for missing values). In both datasets modern reference samples make up the biggest individual group of data, but ancient data with C14 or contextual age information together surpassed this set in size. Where the PAA features age information for all its samples, the PCA lacks some at the time of writing.

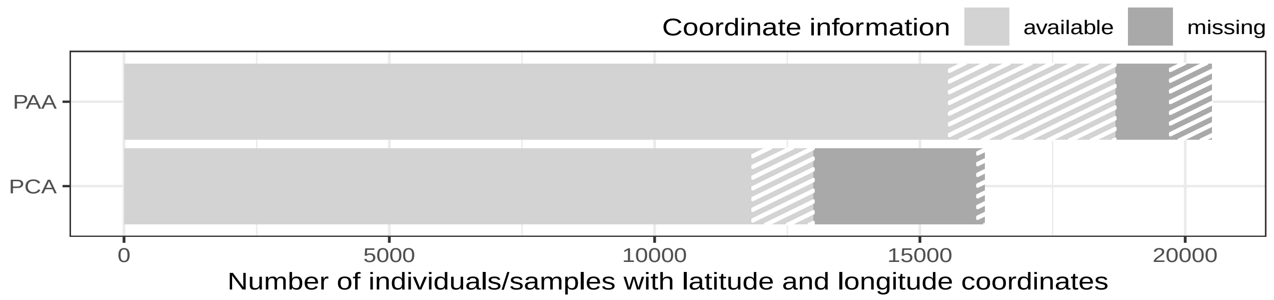

**Supplementary Figure 7:** Corresponds to Figure 5F in the main text.

The .janno columns `Latitude` and `Longitude` include spatial coordinates for each sample. Both archives have gaps here, the PCA more severe ones, though.

[4] Kamil Slowikowski. *ggrepel: Automatically Position Non-Overlapping Text Labels with 'ggplot2'*. R package version 0.9.4. 2024.

[5] David Sjoberg. *ggsankey: Sankey, Alluvial and Sankey Bump Plots*. R package version 0.0.99999. 2024.

[6] Diego Hernangómez. *giscoR: Download Map Data from GISCO API - Eurostat*. Version 0.4.0. 2024. DOI: `10.5281/zenodo.4317946`.

[7] Christopher Brown. *hash: Full Featured Implementation of Hash Tables/Associative Arrays/Dictionaries*. R package version 2.2.6.3. 2023.

[8] Edzer Pebesma. "Simple Features for R: Standardized Support for Spatial Vector Data". In: *The R Journal* 10.1 (2018), pp. 439–446. DOI: `10.32614/RJ-2018-009`.

[9] Karthik Ram and Hadley Wickham. *wesanderson: A Wes Anderson Palette Generator*. R package version 0.3.7. 2023.

[10] Hadley Wickham et al. "Welcome to the Tidyverse". In: *Journal of Open Source Software* 4.43 (2019), p. 1686. DOI: 10.21105/joss.01686.

[11] Swapan Mallick and David Reich. *The Allen Ancient DNA Resource (AADR): A curated compendium of ancient human genomes, 54.1.p1 data release [March 6, 2023]*. 2023. DOI: 10.7910/DVN/FFIDCW.

[12] Iain Mathieson et al. "Genome-wide patterns of selection in 230 ancient Eurasians". In: *Nature* 528.7583 (2015), pp. 499–503. DOI: 10.1038/nature16152.

[13] Nick Patterson et al. "Ancient Admixture in Human History". In: *Genetics* 192.3 (2012), pp. 1065–1093. DOI: 10.1534/genetics.112.145037.
